# Structural and Spectroscopic Basis for Catalysis by a Class C Radical *S*-adenosylmethionine Methylase Involved in Nosiheptide/Nocathiacin Biosynthesis

**DOI:** 10.64898/2026.01.19.700444

**Authors:** Bo Wang, Hayley L. Knox, Nicholas J. York, Matthew I. Radle, Alexey Silakov, Squire J. Booker

## Abstract

Nosiheptide (NOS) is a ribosomally synthesized and post-translationally modified peptide (RiPP) natural product that exhibits potent antibiotic activity against multiple bacterial pathogens. NOS features a core macrocyclic peptide containing thiazoles, dehydrated serine and threonine residues, and a 3-hydroxypyridine ring. In addition to the macrocycle, NOS possesses a side-ring system formed by a 3-methyl-2-indolic acid (MIA) bridge that connects to glutamyl and cysteinyl residues on the core peptide via ester and thioester linkages. This unique side-ring is installed by the class C radical *S*-adenosylmethionine (SAM) methylase NosN. Here, we report three X-ray crystal structures of the NosN homolog, NocN, at resolutions of 1.4 Å, 1.84 Å, and 1.78 Å under anaerobic conditions, representing the first structural characterization of a class C radical SAM methylase. The structures reveal clear electron density for two bound SAM molecules. Remarkably, the C5′ atom of SAM^I^, which coordinates to the [Fe_4_S_4_] cluster, lies 3.5 Å from the methyl group of SAM^II^ and is properly positioned for direct hydrogen atom abstraction. A structure containing a product mimic illustrates how NocN engages its substrate and identifies Tyr276 as a key catalytic residue. The structure further suggests that the sulfonium center of SAM^II^ may undergo epimerization to facilitate radical attack. Finally, electron paramagnetic resonance spectroscopy identifies a paramagnetic species consistent with the addition of the SAM^II^-derived methylene radical to the MIA substrate.

## Introduction

Class C radical *S*-adenosylmethionine (SAM) methylases catalyze chemically challenging transformations on non-nucleophilic sp²-hybridized carbons, most commonly in the biosynthetic pathways of complex natural products. Characterized members of this family carry out one-carbon transfer reactions—typically methylation, lactonization, or cyclopropanation (**Figure 1a**).^1–4^ These enzymes, which belong to the HemN family of radical SAM (RS) proteins, share a distinctive mechanistic feature: the use of two simultaneously bound SAM molecules (SAM^I^ and SAM^II^) during catalysis.^3,4^ Despite their diverse chemical outcomes, all characterized class C radical SAM methylases (RSMs) are proposed to share common initial steps. The first SAM molecule (SAM^I^), coordinated to the unique iron of the [Fe_4_S_4_] cluster cofactor, undergoes a reductive cleavage to yield methionine and a 5′-deoxyadenosyl radical (5′-dA•)—a hallmark step of RS enzymes, with the exception of the cobalamin-dependent RS methylase TsrM.^5,6^ The 5′-dA• abstracts a hydrogen atom (H•) from the methyl group of the second SAM molecule (SAM^II^), generating a methylene radical. This reactive intermediate attacks the target sp²-hybridized carbon of the substrate to form a covalent SAM–substrate adduct containing an unpaired electron. Although this radical intermediate has been postulated for several class C methylases, it has not yet been directly characterized. The ensuing steps vary among enzymes and yield diverse outcomes, including methylated, lactonized, or cyclopropanated products.^7–14^

**Figure 1.**
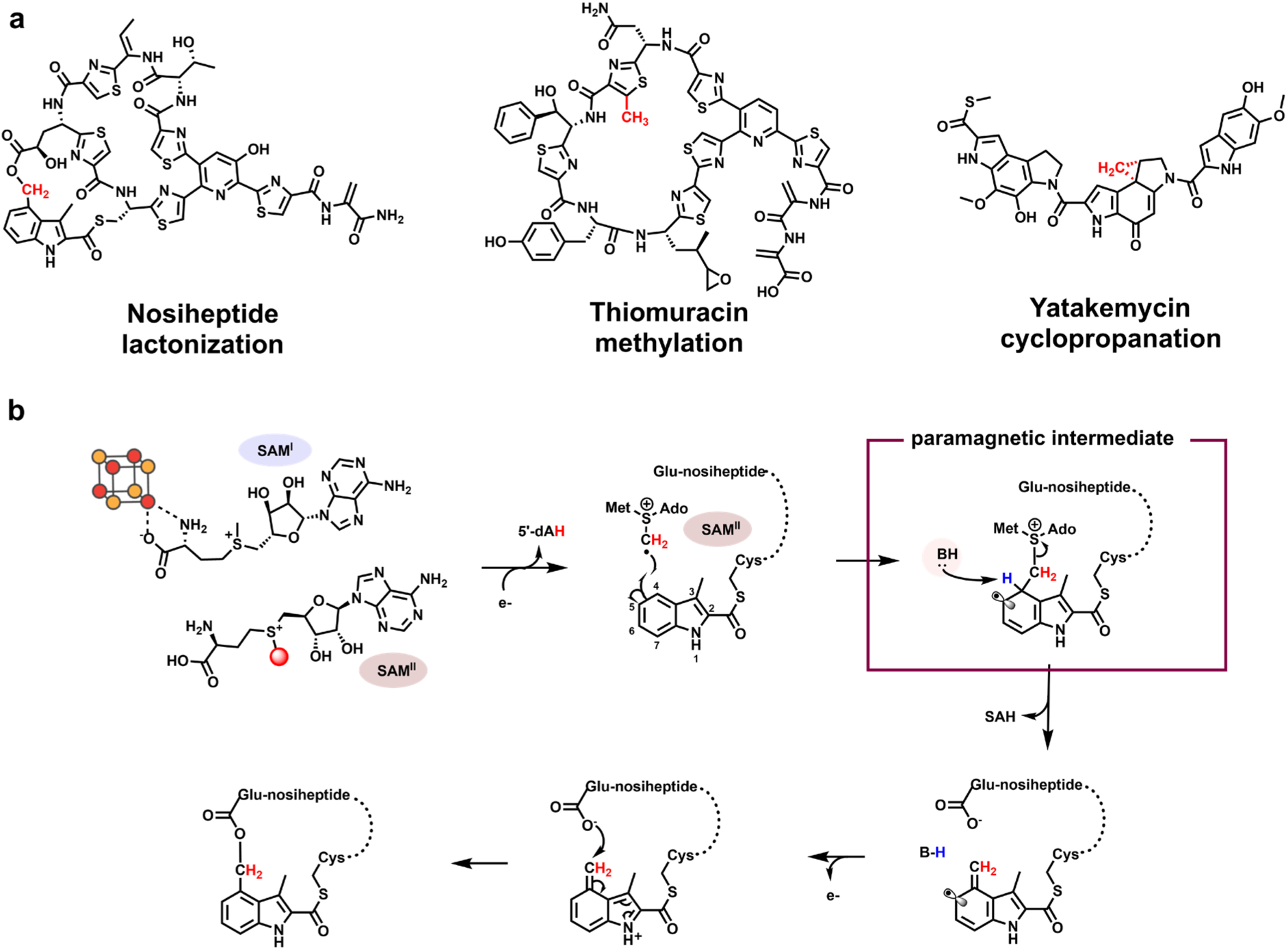
**a.** Chemical structures of nosiheptide, thiomuracin, and yatakemyin. The methyl or methylene groups, highlighted in red, are introduced by the class C RSMs NosN, TbtI, and C10P, respectively; **b.** Proposed catalytic mechanism of side-ring closure catalyzed by NosN. Initial steps, shared by all characterized class C RSMs, include generating a 5’-dA• through the reductive cleavage of SAM^I^ and abstracting a H• from the methyl group of SAM^II^ by the 5’-dA•.

To date, three class C RSMs—NosN,⁹ TbtI,¹⁰ and C10P¹¹—have been investigated in detail through in vitro studies. NosN participates in the biosynthesis of nosiheptide (NOS), a ribosomally synthesized and post-translationally modified peptide (RiPP) antibiotic with potent activity against multiple human bacterial pathogens. Although NosN can catalyze methylation in vitro under certain conditions, its physiological role is to form the lactone linkage between the side chain of Glu6 and C4 of 3,4-dimethylindolic acid thioesterified to Cys8, thereby generating the side-ring of NOS.^9,15^ Mechanistic studies support the reaction pathway illustrated in **Figure 1b**. SAM^I^ is reductively cleaved to generate a 5′-dA•, which abstracts a hydrogen atom (H•) from the methyl group of SAM^II^. The resulting methylene radical adds to C4 of the 3-methyl-2-indolic acid (MIA) moiety to yield a SAM–MIA adduct. Subsequent deprotonation at C4 triggers the release of *S*-adenosylhomocysteine (SAH), followed by electron loss and lactone formation through attack by the side chain of Glu6, completing the NOS side-ring closure.

In this work, we provide both structural and spectroscopic evidence supporting this mechanism. We report three X-ray crystal structures of NocN at 1.4 Å, 1.84 Å, and 1.78 Å resolution under anaerobic conditions, the first representative structures of a class C RSM. NocN shares 75% sequence identity with NosN and catalyzes a nearly identical transformation in the biosynthesis of the related antibiotic nocathiacin (NOC). The NocN structure reveals clear electron density for two bound SAM molecules, with the 5′ carbon of SAM^I^ positioned 3.5 Å from the methyl group of SAM^II^ in an orientation consistent with direct H• abstraction. A second structure containing a product mimic reveals two binding sites—one in the C-terminal region and one within the presumed active site. The latter suggests that the sulfonium center of SAM^II^ may epimerize upon radical formation to facilitate radical attack, a hypothesis that is supported by density functional theory (DFT) calculations. Additionally, the structure and associated biochemical studies indicate that Tyr276 is a key catalytic residue. Finally, electron paramagnetic resonance (EPR) spectroscopy provides experimental evidence for the transient radical intermediate formed upon the addition of the SAM-derived methylene radical to C4 of MIA.

## RESULTS

### NocN catalyzes side-ring formation on a substrate peptide mimic

We initially attempted to determine the structure of *Streptomyces actuosus* NosN (UniProt ID: C6FX53) but were unable to obtain crystals suitable for diffraction. We therefore turned our attention to NocN from *Nocardia* sp. ATCC 202099 (uniprot ID: E5DUI5), which shares 75% sequence identity with NosN. NocN is involved in the biosynthesis of nocathiacin, a thiopeptide natural product that exhibits a chemical structure like that of NOS (**Figure S1)**.^16^ The NOS core precursor peptide sequence is SCTTCEC<u>C</u>CSCSS, while that for NOC is SCTTCEC<u>S</u>CSCSS. The only difference is that MIA is attached to the side chain of the indicated cysteine in NOS and the indicated serine in NOC via thioester and ester linkages, respectively. Compound **1**, a truncated substrate lacking a leader peptide that had previously been used to probe NosN’s substrate specificity, was also expected to serve as a substrate surrogate for NocN.^9^ In a reaction containing NocN, compound **1**, dithionite (DT), and SAM, the expected ring-closed product (*m/z* 1440) is observed by mass spectrometry (**Figure S2ab**). When *S*-adenosyl-[*methyl*-^2^H_3_]methionine (*d_3_*-SAM) is substituted for SAM in the reaction, a product exhibiting an increase of two mass units is observed (**Figure S2cd)**, indicating that NocN transfers a –CH_2_– unit from the methyl group of SAM during the formation of NOC’s side ring. Identical results were obtained in similar studies of NosN.^9,15^ Interestingly, two additional peaks are also observed, corresponding to products containing one (M-1) or no (M-2) deuteriums (**Figure S2d**). A similar isotopic pattern was also observed in our previous studies of NosN using the smaller tripeptide substrate **3** (**Figure S3a**). This unexpected isotope distribution suggests that deuterium exchange occurs during catalysis. To determine whether exchange involves the C5′ hydrogens of SAM, reactions were conducted with *d_7_*-SAM, which contains one deuterium at C3′ and C4′, two deuteriums at C5′, and three deuteriums on the methyl group (**Figure S3b**).^10^ In a reaction using *d_7_-*SAM and compound **3**, the product exhibits an *m/z* of 698, corresponding to the ring-closed species with a CD_2_ group (**Figure S3c**). Neither M-1 nor M-2 peaks are observed, indicating that hydrogen exchange involves C5′ of SAM.

### The NocN Radical SAM Domain Binds Two SAM Molecules Simultaneously

NocN was crystallized under anoxic conditions in the presence of AzaSAM (PDB ID: 9P2U), SAM (PDB ID: 9P3B), or a combination of SAH and a synthetic side-ring-closed (SRC) product mimic (PDB ID: 9P3C), and the structures were determined to resolutions of 1.40 Å, 1.84 Å, and 1.78 Å, respectively (**Table S1**). NocN is composed of two modular domains: a RS domain (residues 21-267) and a C-terminal domain (residues 268-420) (**Figure 2a**).^17^ The overall structure of NocN shows marked similarities to that of HemN, despite their low sequence identity (25.4%) and different catalytic reactions. The two structures exhibit a root-mean-square deviation (RMSD) of 1.8 Å over the α-carbons of their entire polypeptide chains (**Figure S4**).^18^ HemN has an additional domain, a loosely bound "trip-wire" at the N-terminus (residues 4-35), which may be involved in the association or dissociation of substrates, intermediates, and products.^12,18^ NocN lacks this domain, although its first 20 amino acids are disordered.

**Figure 2.**
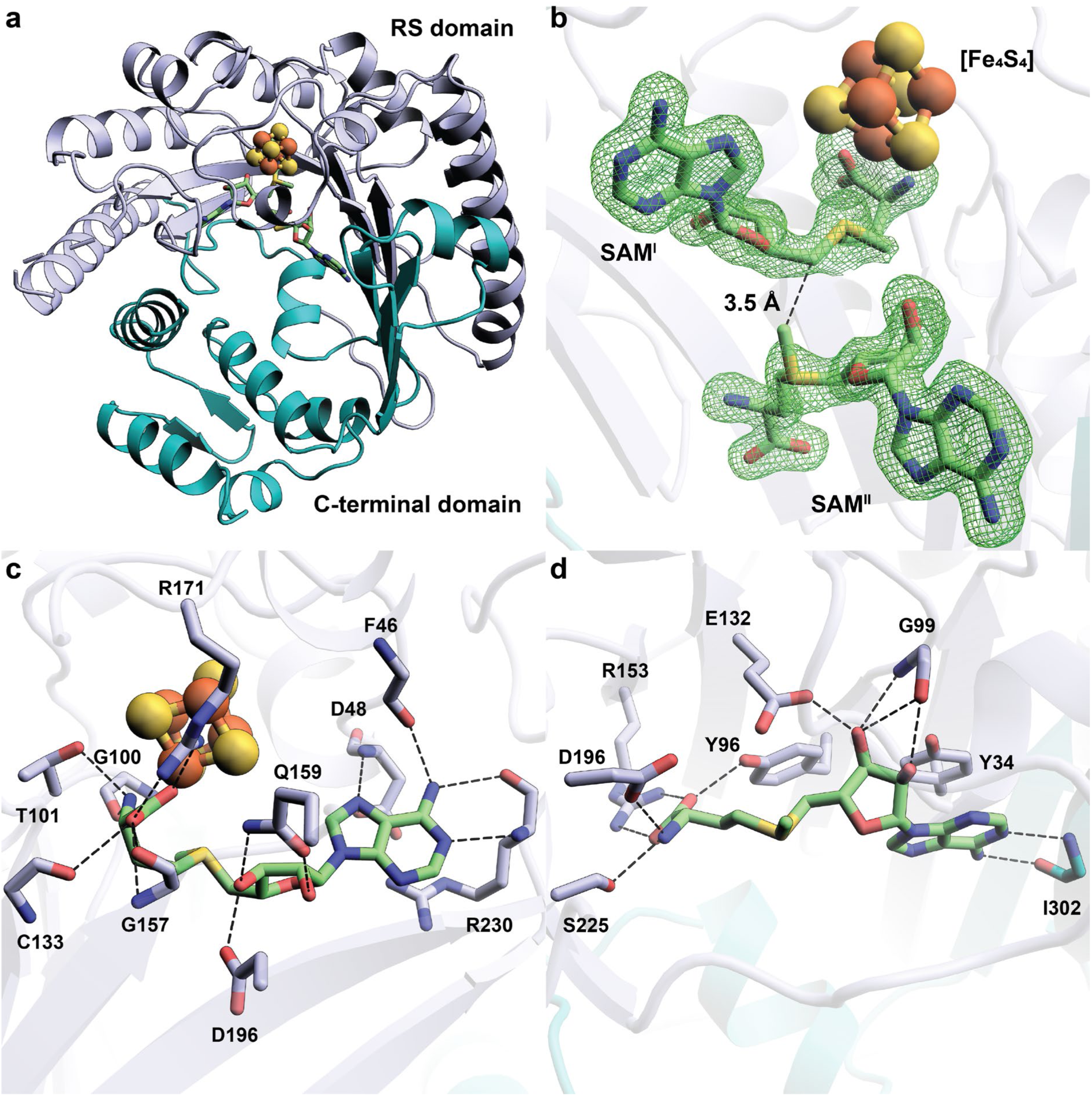
**a.** The overall structure of NocN shown as a ribbon diagram and colored by domains. The RS domain is shown in light blue. The C-terminal domain is shown in teal; **b**. Closeup of SAM^I^, SAM^II^, and the [Fe_4_S_4_] cluster within the active site of the NocN structure. Fo-Fc omit maps are shown for SAM^I^ and SAM^II^ (green mesh, contoured at 3.0 σ). The distance between C5′ of SAM^I^ and the methyl group of SAM^II^ is shown as a black dashed line; **c.** Interaction network between NocN and SAM^I^; **d.** Interaction network between NocN and SAM^II^.

The RS domain forms the core of the NocN structure, comprising a shortened (ß⍺)_6_ triosephosphate isomerase (TIM) barrel. Like most other RS enzymes, NocN’s RS domain houses a [Fe_4_S_4_] cluster ligated by cysteines in the canonical ^40^CysX ^44^CysX ^47^Cys RS motif. NocN’s and HemN’s shortened TIM barrels are similar to that of pyruvate formate-lyase activating enzyme (PFL-AE), but are more splayed (**Figure S5**).^19–21^ Both NocN and HemN must bind two molecules of SAM simultaneously, along with their substrates for catalysis, whereas PFL-AE binds only one SAM molecule and its protein substrate. We observe clear electron density for two SAM-related molecules in the RS domain of NocN (**Figure 2b**). SAM^I^ binds to the unique iron of the [Fe_4_S_4_] cluster via its amino and carboxylate moieties, which is required for its reductive cleavage to yield a 5’-dA• and methionine. The binding is facilitated by residues from NocN’s RS domain through an interaction network observed in other RS enzyme structures (**Figure 2c, Figure S6**).^17,22^ For example, the amino group coordinating the unique iron forms H-bonds with Gly100 and Thr101, which form a glycine-rich "GGE" motif.^22^ In addition to H-bonding with Cys133 and Gly157, the carboxylate group that coordinates the unique iron also forms a salt bridge with the guanidinium sidechain of Arg171. The polar and hydrophobic contacts with the two hydroxyl groups of the ribose moiety and the adenine moiety are similar to those in other RS enzyme structures.^23–26^ A π-cation interaction between the adenine ring and the guanidium sidechain of R230 is observed in the NocN structure but not in the structure of other RS enzymes, including HemN.^18^ In our NocN structure with the SRC product bound, R230 also establishes a π-cation interaction with a thiazole moiety of the substrate, indicating the importance of R230 in both SAM and substrate recognition through π-cation-π stacking (**Figure S7**). The methyl group of SAM^II^ points at C5′ of SAM^I^ with a distance of 3.5 Å between the two carbon atoms, consistent with H• abstraction from SAM^II^’s methyl group by the 5′-dA• (**Figure 2b**). Most of the interactions with SAM^II^ are from the RS domain of NocN. The only exception is the H-bonds between the adenine ring of SAM^II^ and the main chain of Ile302, which is from the C-terminal domain (**Figure 2d**). Asp196 is the only residue that establishes direct H-bonds with both SAM^I^ (through the 3′-OH of the ribose moiety) and SAM^II^ (through the amino group of the methionine moiety). The binding of the adenine and ribose moieties of SAM^II^ is identical to that of SAM^II^ observed in HemN, with almost the same interaction networks (**Figure S6**). In particular, we observe a π-stacking interaction between Tyr34 and the adenine ring of SAM^II^, which is reminiscent of the π-stacking observed for SAM^II^ in HemN. In HemN, this residue is Tyr56, which has been shown to be crucial for catalysis. Tyr56 has been suggested to be involved in cluster stabilization, as slight cluster degradation occurs upon substituting this residue; however, the Tyr residue is not proximal to the cluster.^18,27^ Furthermore, as shown in **Figure S8**, this Tyr residue appears to be conserved among HemN-like RSMs, suggesting that its π-stacking interaction is important for SAM^II^ binding. The polypeptide chains and cofactors/co-substrates of NocN and HemN overlay well. However, the methionine moieties of SAM^II^ from NocN and SAM^II^ from HemN show distinct binding poses (**Figure S9**). It should be noted that SAM^II^ in the HemN structure adopts an R,S configuration (namely epimerized SAM), whereas naturally occurring SAM has the S,S configuration. In NocN, the methionine moiety of SAM^II^ points toward β5 and β6 of the TIM barrel and forms a tight H-bonding network with residues from the TIM barrel, including Tyr96 (2.8 Å), Arg153 (3.0 and 3.1 Å), Asp196 (2.7 Å), and Ser225 (2.9 Å) (**Figure 2d**). However, the methionine moiety of epimerized SAM^II^ from HemN points toward the outside of the active site and does not form any direct H-bond interactions with any HemN residues. The different arrangements of SAM^II^ in NocN and HemN may reflect how SAM^II^ engages its substrate and its subsequent fate upon H• abstraction from its methyl group. A recently revised mechanism for HemN shows that the methylene radical generated on SAM^II^ abstracts the pro-(*S*) hydrogen from the β-carbon of the propionate moiety of coproporphyrinogen III to promote decarboxylation.^12^ However, the same radical species attacks an sp^2^-hybridized carbon in the NosN reaction instead of abstracting an H• from the substrate. In the NocN structure with AzaSAM bound, the two AzaSAM molecules bind in slightly different poses from those of the two SAM molecules observed. The difference is mainly reflected by the distance of 5.9 Å from C5′ of AzaSAM^I^ to the methyl group of AzaSAM^II^ (**Figure S10**). AzaSAM is a SAM mimic with a nitrogen atom replacing the sulfonium atom. Thus, we believe that differences in bond lengths account for the slight difference between the two structures.

### Substrate-Bound Structures of NocN

A structure of NocN containing SAH^I^, SAH^II^, and two SRC products (SRC1 and SRC2) was also determined at 1.78 Å resolution. The SRC product is a product mimic containing a truncation of three amino acids at the C-terminus and lacking the leader peptide (37 amino acids) at the N-terminus (structure and numbering of thiazole rings are shown in **Figure S11**). Our biochemical studies have shown that NocN and NosN can catalyze their reactions on the corresponding substrate mimic lacking the leader peptide. NocN’s C-terminal domain exhibits three structural features (**Figure S12a**). Feature I is a loop-helix-loop (α_7_ and two loops on both ends); Feature II consists of four antiparallel ß sheets aligning with the core (ß⍺)_6_ fold; and Feature III is a bundle of three α-helices (α_8_-α_10_), which form a shallow cavity. Continuous electron density for SRC2 is observed in the cavity (**Figure S12b**). SRC2 establishes direct polar interactions with either the backbones or sidechains of residues from these three α-helices of Feature III (**Figure S12c**). The cavity’s environment is hydrophobic, primarily due to the presence of numerous leucine residues (**Figure S12d**). However, very few aromatic residues, which can form π-π stacking interactions with the aromatic-rich SRC2, are observed. C4′ of MIA, the carbon attacked by the methylene radical, is 19.1 Å away from the sulfur atom of SAH^II^. Thus, SRC2 is not likely to be associated with NocN’s catalytic activity. Why SRC2 binds in this location, and the function of the shallow cavity in the C-terminal domain, is unclear.

The active site of NocN, into which two SAH molecules and SRC1 bind, is formed between the RS domain and the C-terminal domain (**Figure 3a**). The two SAM molecules interact primarily with the RS domain, whereas most interactions with SRC1 occur in the C-terminal domain. The electron density for SRC1 in the active site is not as continuous as for SRC2. However, the density around the MIA moiety, where the chemistry occurs, is sufficient for modeling it (**Figure 3b**). The NocN active site is narrow, into which the A-shaped SRC1 molecule inserts its MIA moiety. The C-terminus of SRC1 exits the active site along the antiparallel ß-sheets (Feature II), while the N-terminus of SRC1 exits the active site along α_10_ from Feature III. The sulfur atom of SAH^II^ is 3.4 Å from the appended methylene group and 4.1 Å from C4′ of MIA. If we project a methyl group onto SAH^II^, it is positioned between C5′ of 5′-dA and C4′ of MIA. However, this methyl group points away from C4′ of MIA (**Figure 3c**). If the substrate binds similarly to SRC1 in our structure, SAM^II^ would require a significant conformational change for the methylene radical to approach C4′ of MIA, such as rotating along the C5′-S bond (**Figure S13**). However, such a rotation is unlikely because it would require the breaking of several H-bonds to NocN. Another possibility is that the sulfonium epimerizes upon forming the methylene radical, thereby making the radical attack much more favored (**Figure 3d**). SAM is well known to epimerize in solution in a pH-independent manner. The rate constant for this process has been estimated to be ∼3.2 × 10^-6^ s^-1^ at 37 °C, pH 7.5, and constant ionic strength.^28–30^ It has also been reported that the electronegativity of substituents around the sulfonium atom can affect epimerization rates.^30–33^ Thus, the methylene radical, formed upon H• abstraction by the 5′-dA•, may accelerate epimerization.

**Figure 3.**
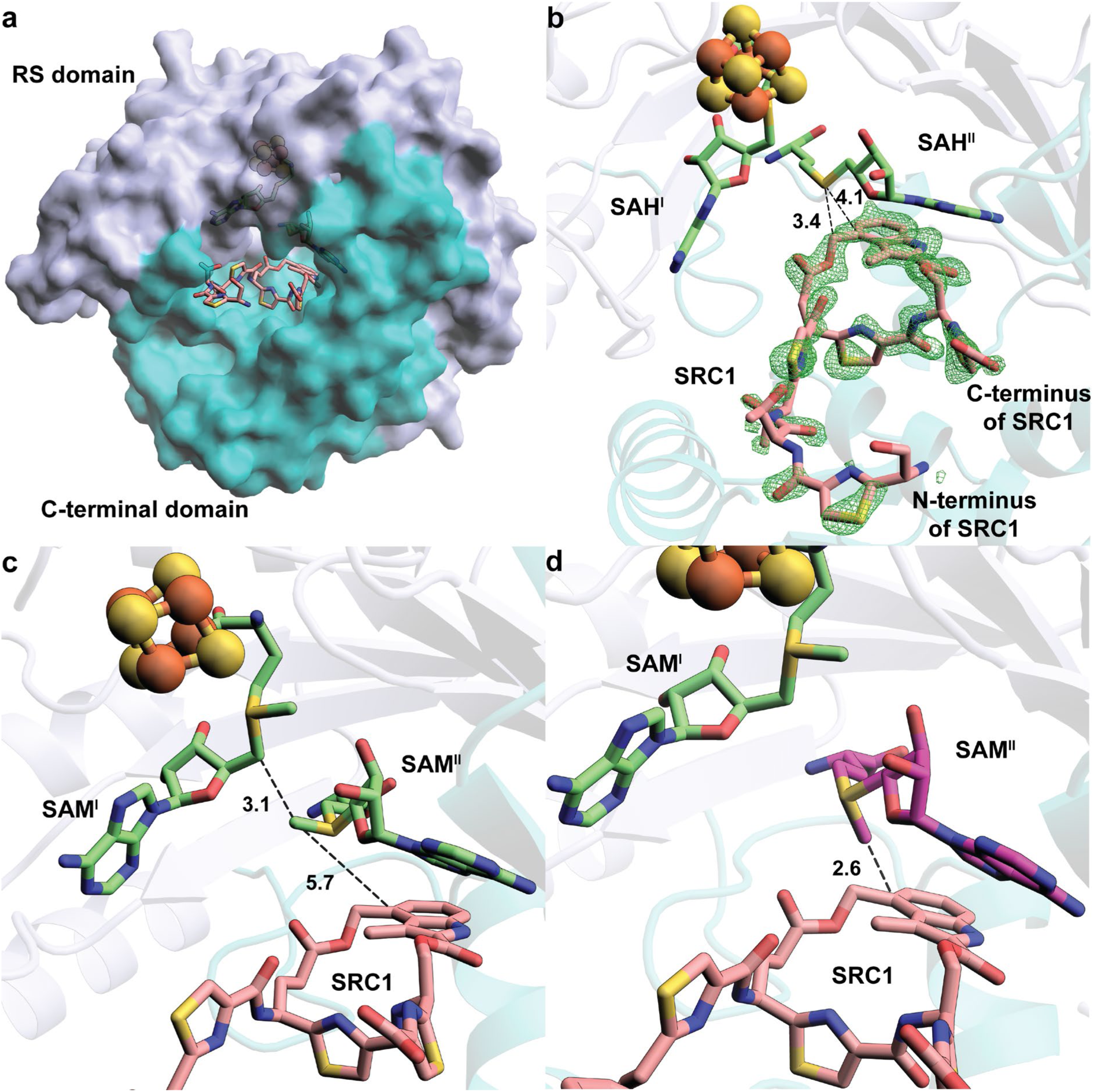
**a.** The overall structure of NocN is illustrated as a surface diagram and colored by domains. The RS domain is shown in light blue. The C-terminal domain is shown in teal; **b.** Closeup of SAH^I^, SAH^II^, and the SRC1 product within the active site of the NocN structure. Fo-Fc omit maps are shown for SRC1 (green mesh, contoured at 3.0 σ). The distances between the sulfur atom of SAH^II^ and the appended methylene group and C4′ of MIA are shown as black dashed lines; **c.** Orientation of SAM molecules in the SRC-bound structure of NocN. The methyl group of SAM^II^ is between C5′ of SAM^I^ and C4′ of MIA, but pointing in the opposite direction of C4′ of MIA; **d.** If the sulfonium epimerizes upon H• abstraction from the methyl group of SAM^II^, the distance and orientation of the methylene radical become feasible for the radical attack step.

To test whether epimerization is a reasonable catalytic strategy, we used DFT to perform a relaxed-surface scan of both SAM and the SAM methylene radical epimerizing from *S* to *R* at the sulfonium (**Figure S14**). Starting coordinates were obtained from our NocN structure, and the methyl/methylene carbon was incrementally shifted by ten degrees during the change in sulfonium stereochemistry, affording a planar transition state (**Figure S14a**). The energy diagram of this scan (**Figure S14b**) reveals 27.3 and 20.7 kcal/mol activation barriers for SAM and SAM methylene radical epimerization, respectively. Therefore, generating the methylene radical lowers the epimerization activation barrier by 6.6 kcal/mol. We note that these values may not reflect the epimerization activation barriers in the protein’s active site. However, our results indicate that a methylene radical on SAM will substantially enhance SAM epimerization. A possible explanation for this stabilization is the delocalization of the radical onto the sulfonium cation. At the lowest-energy *S*-configuration calculated for the sulfonium methylene radical, the Mulliken spin population is 0.94 on the methylene carbon and 0.03 on sulfur. At the highest-energy planar transition state, these values shift to 0.85 for carbon and 0.18 for sulfur, indicating delocalization of the radical onto sulfur. This distribution is also evident in the visualization of the singly occupied molecular orbital (SOMO) of the transition state, which shows a typical π* molecular orbital between the methylene carbon and sulfur (**Figure S14c**).

### Tyr276 Plays an Important Role in NocN Catalysis

The active site of NocN is rich in aromatic residues, and π-π stacking with Phe283, Tyr287, Phe288, and the adenine ring of SAH^II^ plays an important role in positioning the MIA moiety of the SRC1 species (**Figure 4a**). Interestingly, all three aromatic residues are located in the α-helix of Feature I, indicating the importance of this secondary structure in engaging the substrate. Surprisingly, only two direct H-bonds are observed between NocN and the SRC1 species. These include bonds between the side chain of Tyr276 with both oxygen atoms of the carboxyl side chain of the thiazolyl glutamate (ThzGlu) and Tyr287 with the carbonyl group of the same ThzGlu. Tyr276 is close to the carboxyl side chain of ThzGlu and the appended methylene carbon from SAM^II^. Potentially, it could serve as the base that deprotonates the aromatic radical intermediate, facilitating the elimination of SAH in our proposed mechanism (**Figure 1b**). It may also serve as a proton shuttle to assist the nucleophilic attack of the carboxyl side chain of ThzGlu upon forming the electrophilic methylene in our proposed mechanism. Tyr227 does not interact directly with the carboxyl side chain of ThzGlu, but it may assist Tyr276 in either deprotonation. To study their potential roles in catalysis, we performed mutagenesis studies on NosN, which is isolated in better yields than NocN. Tyr202 (corresponding to Tyr227 of NocN) and Tyr251 (corresponding to Tyr276 of NocN) were changed to phenylalanine. As shown in **Figure 4b and Figure S15**, wild-type NosN shows a rate constant of 0.024 ± 0.00099 min^-1^, and the Y202F variant shows slightly diminished activity (red trace) with a rate constant of 0.018 ± 0.0029 min^-1^. However, the Y251F variant and the Y202/251F double variant show significantly lower activities (0.0033 ± 0.00047 min^-1^ for Y251F, 0.0013 ± 0.000045 min^-1^ for Y202/251F), suggesting the importance of Tyr251 (Tyr276 for NocN) during catalysis.

**Figure 4.**
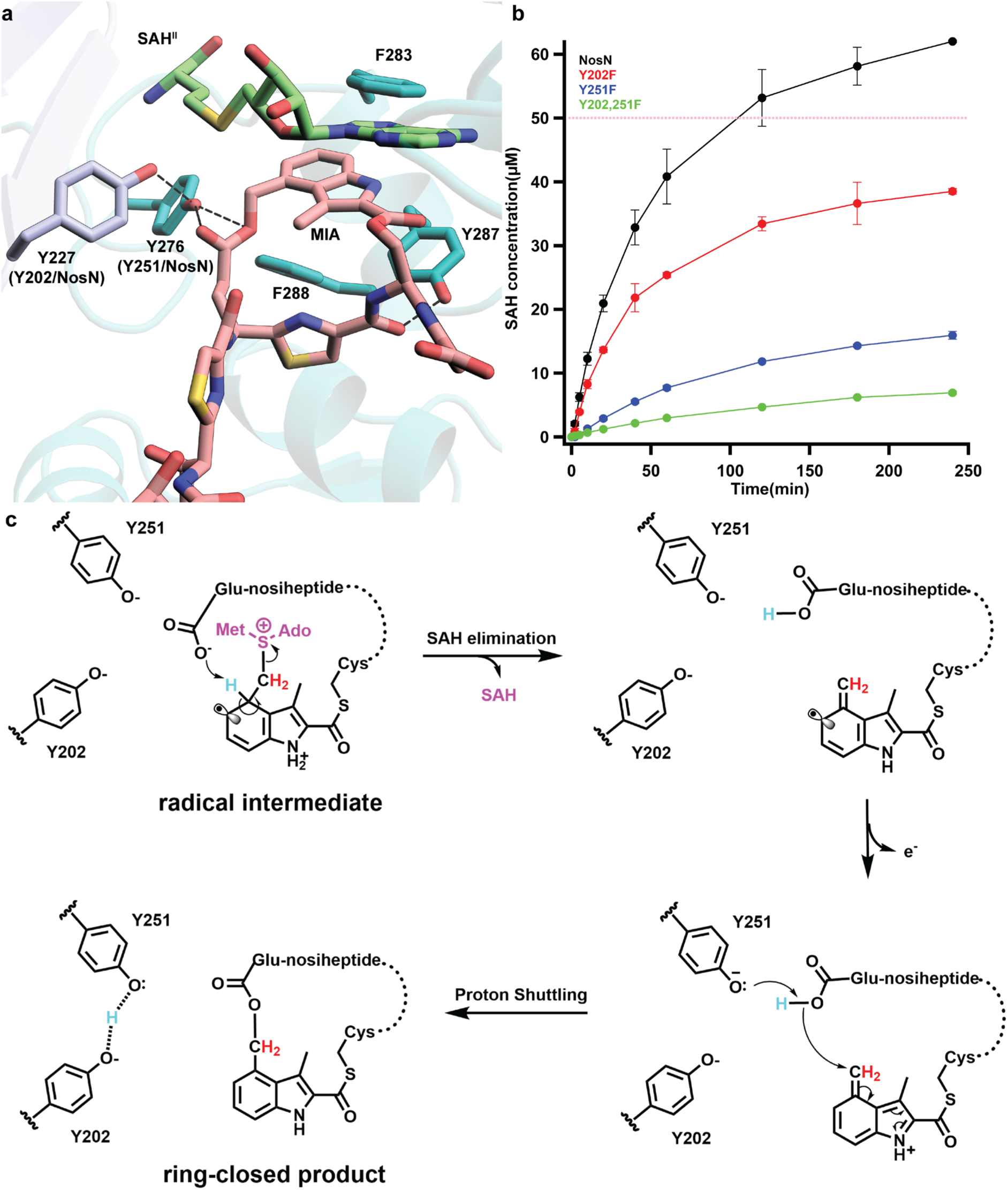
**a.** Interaction network between NocN and SRC1; **b.** Time-course reactions performed using NosN (black), Y202F (red), Y251F (blue), or Y202,251F (green), tracking the formation of SAH. Each reaction is performed in triplicate. The dashed pink line indicates the concentration of enzyme (50 µM) used in the reaction. Turnover numbers derived from the initial rates of these time courses are summarized in Figure S15; **c.** Replenished catalytic mechanism of NosN.

Recent studies by Xiong et al. showed that NosN’s in vivo substrate contains the leader peptide, thiazoles, dehydrated alanines, and butyrine.^34^ In a previous study, we detected two SAM adducts using compound **5,** a shortened version of NosN’s substrate that lacks the leader peptide but contains all thiazoles and three dehydrated amino acids. We believe that the rigid structure of compound **5** disfavors the deprotonation of the radical intermediate, leading to its oxidation or reduction to form the SAM adducts (**Figure S16**).^9^ Thus, the detection of SAM adducts correlates with the slow deprotonation of the radical intermediate. If Tyr251 (Tyr276 for NocN) or Tyr202 (Tyr227 for NocN) is the base that deprotonates the radical intermediate, the accumulation of the SAM adducts in reactions using the Y251F or Y202F variants should be observed. In a reaction of wild-type NosN using compound **3**, a small amount of the SAM adduct is detected at various time points throughout the 4 h incubation, suggesting that the radical species is long-lived, which is supported by the slow decay of the EPR signal (see below) (**Figure S17a**). This slow decay is likely due to a rate-limiting deprotonation of the radical intermediate during catalysis. Hence, upon terminating the reaction by adding acid, the radical species are quenched, yielding the SAM adducts. Quantification of the SAM adducts shows a time-dependent increase in concentration; however, the ratio of SAM adducts to SAH remains constant at ∼14% (**Figure S17b, c**). Contrastingly, in the reaction of the NosN Y202F and Y251F variants, no accumulation of SAM adducts is observed, suggesting that neither residue acts as a direct general base.

The carboxyl group of ThzGlu could also be the base that deprotonates the radical intermediate. Compound **6**, which has an amide in place of the carboxylate side chain, was synthesized as an analog that should not be able to deprotonate the radical intermediate (**Figure S18a**). As shown in **Figure S18b**, this analog affords a substantial increase in SAM adduct formation as compared to NosN reactions with compound **3**. Quantification of the SAM adduct shows a rapid burst followed by a slow decay (**Figure S18c**). The ratio of SAM adducts versus SAH peaks at 60% and decreases to 30% (**Figure S18d**). However, methylated products rather than ring-closed products are detected when using compound **6**. These methylated products were verified using *d_3_-*SAM and [*methyl*-*^13^C*]-SAM (**Figure S19**). These data suggest that the carboxyl group of ThzGlu likely serves as the base that deprotonates the radical species, exemplifying substrate-assisted catalysis. However, the observation of methylated products indicates that deprotonation can occur in the absence of the carboxylate of ThzGlu, which is likely mediated by a water molecule. Methylated products were also observed in a previous study using MIA connected to *N*-acetylcysteamine as a substrate, further supporting the hypothesis that water can mediate this deprotonation.^15,35^ Our studies allow us to add more detail to our proposed NosN mechanism of catalysis (**Figure 4c**). The carboxyl side chain of ThzGlu is likely the base that deprotonates the radical intermediate, with the proton being transferred to Tyr251. Upon SAH departure, the resulting carboxylate can attack the exocyclic methylene of MIA, forming the side-ring-closed product.

### Characterization of the Aryl Radical Intermediate by EPR Spectroscopy

A key step in NosN catalysis is the addition of a SAM^II^-derived methylene radical to C4 of the 3-methylindolyl group, generating a paramagnetic species in which SAM^II^ is crosslinked to the substrate via a methylene bridge. This step was examined by EPR spectroscopy. **Figure 5a** shows EPR spectra of NosN samples containing SAM and either compound **1** or compound **5**, with sodium dithionite (DT) used as the reductant to initiate catalysis. The sample with compound **1** was frozen after 5 min at room temperature, whereas the sample with compound **5** was frozen after 40 min. Compounds **1** and **5** are structurally similar; the only difference is that several serine residues in compound **1** are replaced by dehydroalanines in compound **5**, imparting greater rigidity. The EPR spectra recorded at 80 K for both substrates are nearly identical and consistent with a single unpaired electron experiencing multiple hyperfine (HF) interactions. However, the rates of radical formation and decay differ markedly. Samples containing compound **1** reach maximum signal intensity at 5 min, which persists until 10 min and remains detectable for at least 1 h (**Figure S20**). In contrast, samples containing compound **5** require up to 40 min to reach maximum intensity, which persists for at least 90 min (**Figure S21**). This difference likely reflects the increased rigidity of compound **5**, which may hinder the approach of a general base needed to abstract the C4 proton and quench the radical by electron addition or electron loss. The similar EPR spectra for compounds **1** and **5** are consistent with radicals of comparable structure, as expected from the proposed mechanism. The distinct kinetics suggest that the more rigid dehydroalanine residues in compound **5** slow radical decay by restricting the base’s access to the C4 hydrogen of MIA, thereby stabilizing the intermediate until quenching and formation of the SAM adduct.

**Figure 5.**
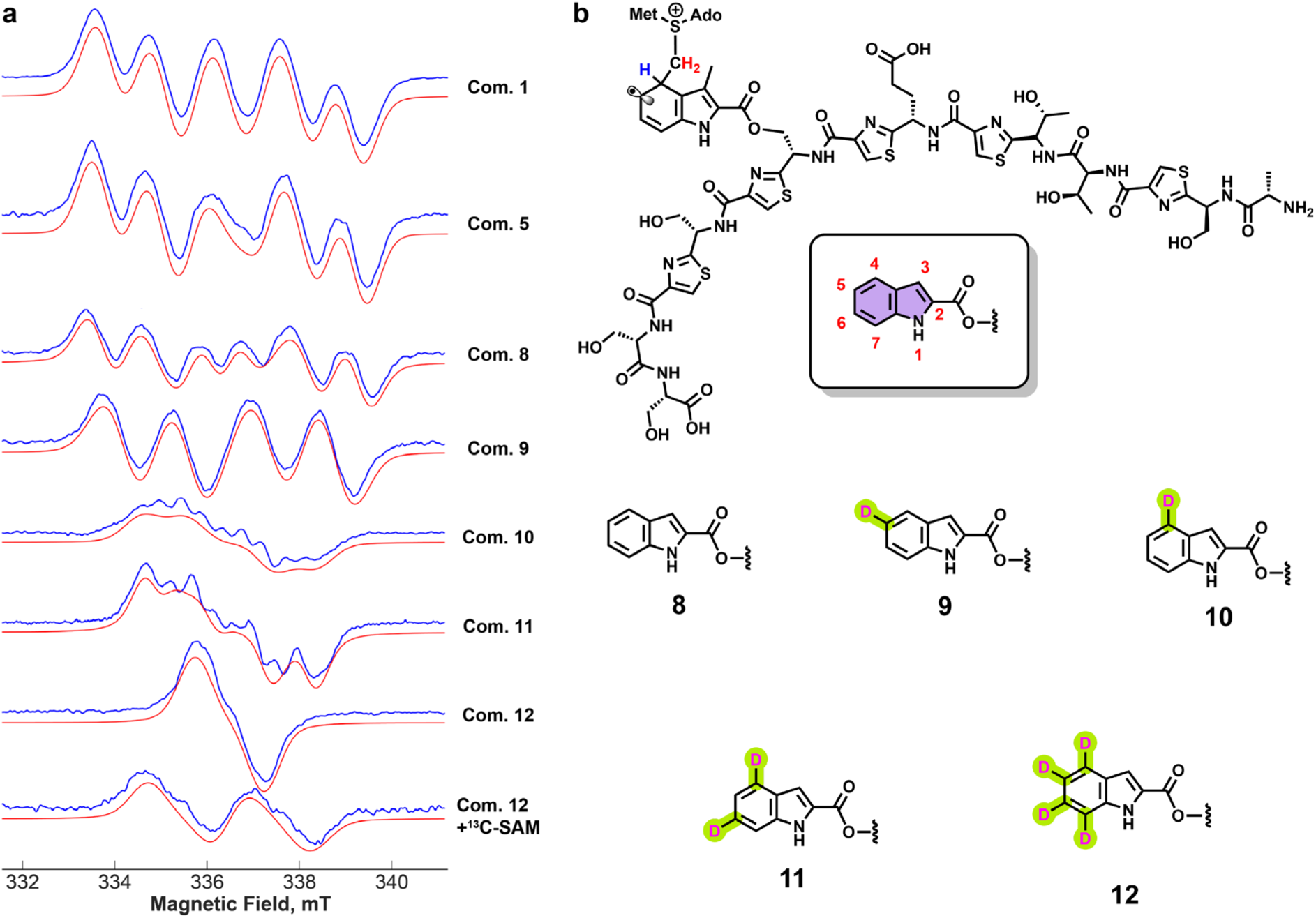
**a.** Comparison of EPR spectra of a paramagnetic species in the NosN reaction with compound **1** at 5 min, compound **5** at 60 min, compounds **8**-**12** at 2 min, and compound **12** + ^13^C-SAM at 2 min. In each EPR spectrum, the blue traces are the experimental spectra, and the red traces are simulations; **b.** Chemical structures of the radical species based on the chemical structure of compound **1**. The structure in the rounded rectangle shows how indolic acid is numbered. The chemical structures below show compound **8** and its deuterated analogs **9**-**12**.

To probe the location of the radical, several isotopically labeled derivatives of compound **1** were synthesized (**Figure 5b**). Compound **8**, an analog lacking the C3-methyl group of 3- methylindolic acid, was synthesized because it more easily allowed regioselective deuterium incorporation. The EPR spectrum of the NosN reaction with compound **8** shows a well-defined six-line pattern similar to that of compound **1**, with minor line-shape differences. Deuterium substitution at C5 (compound **9**) narrows the spectrum to four lines, while labeling at C4 (compound **10**) markedly reduces the fine structure. These results support the assignment of the radical intermediate, with most of the spin density localized on C5 and a strong HF interaction at the C4 hydrogen via hyperconjugation. Deuteration at both C4 and C6 (compound **11**) causes further narrowing, and the isotopologue labeled at C4, C5, C6, and C7 (compound **12**) exhibits an almost complete loss of fine structure, consistent with spin delocalization across the ring. To confirm that the radical arises from SAM crosslinking, perdeuterated compound **12** was reacted with [^13^C-*methyl*]-SAM. The resulting EPR spectrum shows an additional splitting absent in the natural-abundance SAM sample, consistent with isotropic ^13^C HF coupling at C4 due to hyperconjugation.

To aid interpretation, DFT calculations of the EPR parameters for the aryl radical intermediate were performed (**Figure S22**, **Table S2**). The calculated parameters were used as starting points to fit the experimental spectra from isotope-labeling studies. The best-fit parameters deviated only slightly from the computed values, supporting the assignment of the observed spectra to the proposed radical intermediate. Nearly identical HF couplings were obtained for methylated and unmethylated substrates, suggesting minimal structural perturbation from the C3-methyl group. Some deviations between calculated and experimental coupling constants are expected, given that the protein environment was not explicitly modeled. Using parameters derived from compound **8** and its isotopologues, spectra for compounds **1** and **5** could be simulated with only minor adjustments to the HF constants. The primary difference among these analogs lies in the magnitude of the C4-H HF coupling, reflecting subtle changes in spin distribution likely caused by the absence of the C3-methyl group, which may influence substrate positioning within the active site.

## DISCUSSION and CONCLUSION

Class C RSMs, members of the HemN subfamily of RS enzymes, catalyze unusually complex transformations, yet only a few representatives have been characterized to date. Although HemN itself is not a methylase, its crystal structure revealed two SAM molecules bound simultaneously in the active site—a feature now considered diagnostic for HemN-like proteins, including Class C RSMs. This dual-SAM arrangement has been proposed for enzymes such as NosN, TbtI, and C10P, but direct structural evidence has been lacking.

In this study, we obtained a structure of NocN showing two SAM molecules bound in close proximity, providing direct evidence for this defining feature. The 5′-carbon of SAM^I^ lies approximately 4.5 Å from the sulfur and 3.5 Å from the projected methyl group of SAM^II^, strongly suggesting a direct interaction between the two molecules—consistent with the proposed Class C RSM mechanism. Additional support comes from experiments using *d₃-*SAM, in which deuterium from SAM^II^ exchanged with hydrogens at the 5′-position of 5′-dA. As shown in **Figure S3d**, abstraction of a H• from SAM^II^’s methyl group by a 5′-dA• generates a methylene radical, which can re-abstract a hydrogen (or deuterium) from 5′-dAD. Repeating this process leads to partial or complete D–H exchange, yielding CHD• or CD₂• radicals. Subsequent addition of these radicals to the substrate yields ring-closed products showing M–1 and M–2 peaks in the mass spectrum. The predominance of the M–1 peak indicates fewer exchange cycles relative to M–2, consistent with a rapid, reversible exchange within a tightly organized active site. Together, these data strongly support the model of two SAM molecules bound simultaneously in Class C RSMs.

We also obtained a structure of NocN with an SRC species bound. Unexpectedly, the SRC species was observed in both the active site and the C-terminal region. The basis for C-terminal binding is unclear. The N-terminus of the SRC species—corresponding to its leader peptide—lies near a cavity where SRC2 binds. This cavity may represent a RiPP recognition element (RRE), which typically binds leader peptides;^36^ however, bioinformatic analysis using the HHpred Web tool (https://toolkit.tuebingen.mpg.de/) revealed very low sequence similarity to known RREs.^37^ The SRC molecule located in the active site illustrates how the two SAM molecules and the substrate interact. Notably, the methyl group of SAM^II^ in our current structure points **away** from the C4 of MIA, the position expected to receive the methylene radical generated upon H• abstraction by the 5′-dA•. This misalignment suggests that conformational rearrangements may occur during catalysis to reposition the reactive center, although how this would happen is not evident in our current structure. Alternatively, epimerization of the SAM^II^ sulfonium center during catalysis could account for the required reorientation. DFT calculations support the feasibility of this epimerization, although racemic SAM was not detected during turnover. While sulfonium epimerization is known as a degradation pathway for SAM, its involvement as a catalytic step has not been reported previously.

As a comparison, RlmN, is a Class A RSM that methylates the C2 positions of adenosine 2503 (A2503) and adenosine 37 (A37) in ribosomal and transfer RNAs, respectively.^38–41^ Like NosN, RlmN consumes two SAM molecules per turnover, but it operates through a distinct ping-pong mechanism.^40^ The first SAM molecule is used to methylate Cys355 to form *S*-methylcysteine (mCys355) and SAH, which then dissociates. A second SAM molecule subsequently binds and undergoes reductive cleavage to form a 5′-dA•, which abstracts an H• from the methyl group of mCys355. The resulting methylene radical attacks C2 of A2503, and deprotonation by Cys118 yields the methylated product. In this system, mCys355 plays a role analogous to SAM^II^ in NosN.

A comparison of the crystal structures of RlmN (PDB 5HR7) and NocN (with SAM and the SRC species modeled) reveals that the overall architectures do not superimpose well (RMSD = 3.8 Å for 794 aligned atoms), reflecting differences in domain organization.^38^ However, alignment of the RS domains shows that the [Fe₄S₄] clusters overlay closely. Superimposing the two [Fe₄S₄] clusters positions the SAM molecules in nearly identical locations (**Figure 6a**). Remarkably, this alignment places the methylthiol group of mCys355 from RlmN near the methyl group of SAM^II^ in NocN.

**Figure 6.**
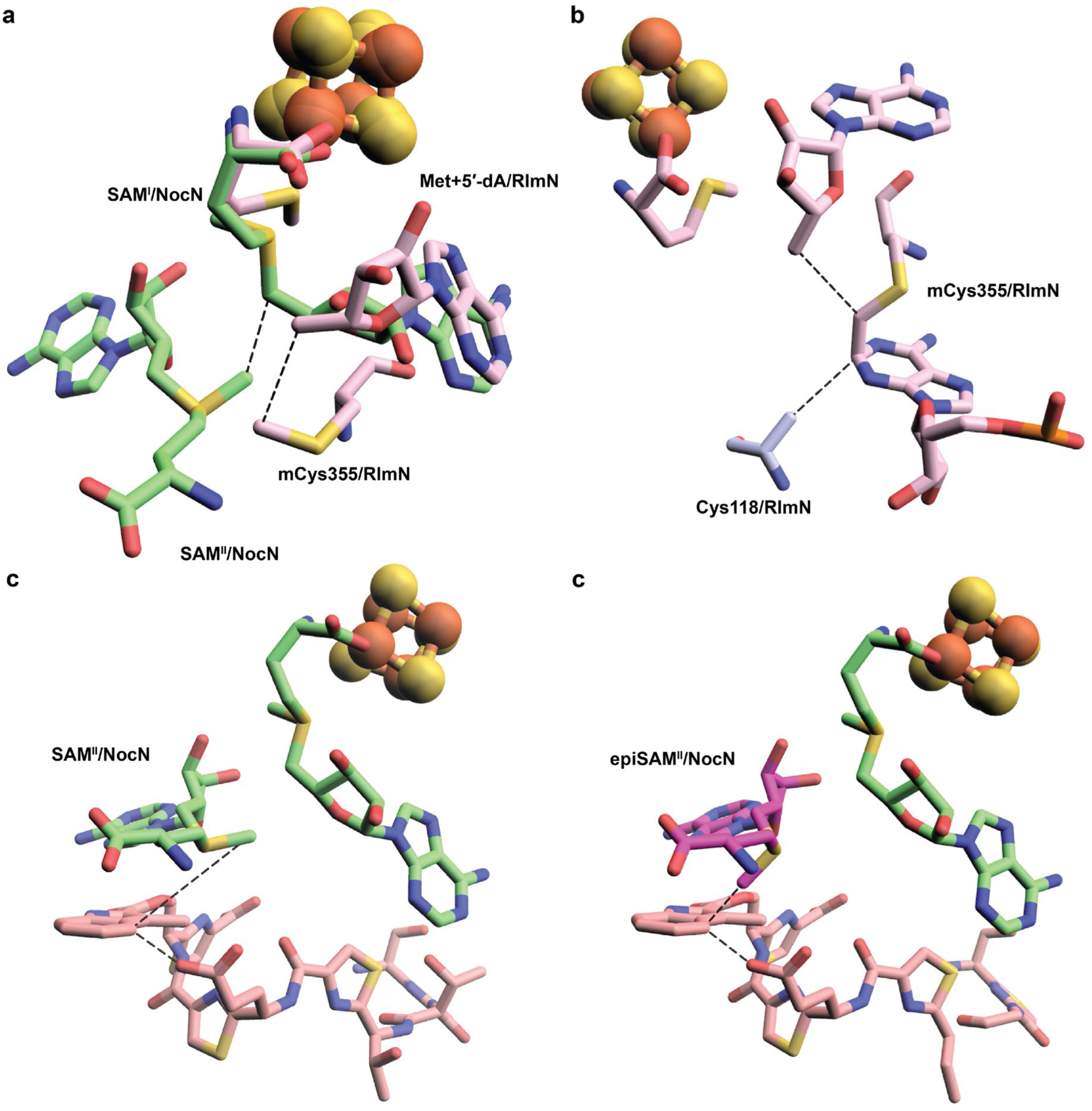
**a.** Structures of RlmN (PDB ID:5HR7) and NocN pair-fitted by [Fe_4_S_4_] cluster. Met, 5′-dA, and mCys355 from RlmN are shown pink; SAM^I^ and SAM^II^ from NocN are shown green; **b**. Closeup of 5′dA, A2503, and Ala355 within the active site of RlmN (PDB ID: 5HR7), showing a “sandwiched” arrangement of how mCys355 radical attacks the SN2-hybridized C8 of A2503 from its upper face and the following deprotonation by Cys355 from the lower face of A2503. In the structure of NocN with SRC bound in the active site, both SAM^II^ without (**c**) or with (**d**) an epimerized sulfonium atom may mimic the “sandwiched” structure observed in RlmN. To generate the figure, the carbon atom between C4 of MIA and the carboxyl side chain of ThzGlu is manually deleted in Pymol.

Established principles of electrophilic radical addition, such as those underlying the Minisci reaction, predict that the methylene radical approaches the electron-rich indole ring in a parallel orientation.^42^ This geometry is evident in the RlmN structure (**Figure 6b**). Assuming the substrate binds to NocN in a manner similar to the SRC species in our current NosN structure, both SAM^II^ (**Figure 6c**) and epimerized SAM^II^ (**Figure 6d**) can reproduce the substrate organization observed in RlmN. In the configuration shown in **Figure 6c**, MIA must shift laterally to sit between SAM^II^ and the carboxylate of ThzGlu, which likely serves as the general base (analogous to Cys118 in RlmN). In the alternative model (**Figure 6d**), MIA would rotate slightly to align the π orbital of its indole ring with the p orbital of the methylene radical, maximizing orbital overlap and facilitating bond formation. Together, our findings define the structural and mechanistic basis of dual-SAM catalysis in Class C RSMs and establish NocN as a paradigm for this enzyme class.

## ASSOCIATED CONTENT

### Supporting Information

Additional supplementary figures, including Figure S1-S22; crystallographic Table S1 and DFT calculation Table S2; experimental procedures for enzyme purification, crystallization, data processing, DFT calculation; synthesis, purification, and NMR characterization of substrats.

## ACKNOWLEDGMENT

This research used resources of the Advanced Photon Source, a US Department of Energy (DOE) Office of Science User Facility operated for the DOE Office of Science by Argonne National Laboratory under contract no. DE-AC02-06CH11357. Use of GM/CA@APS has been funded in whole or in part with federal funds from the National Cancer Institute (ACB-12002) and the National Institute of General Medical Sciences (AGM-12006). The Eiger 16M detector at GM/CA-XSD was funded by NIH grant S10 OD012289. Use of the LS-CAT Sector 21 was supported by the Michigan Economic Development Corporation and the Michigan Technology Tri-Corridor (grant 085P1000817). This research also used the resources of the Berkeley Center for Structural Biology supported in part by the Howard Hughes Medical Institute. The Advanced Light Source is a Department of Energy Office of Science User Facility under contract no. DE-AC02-05CH11231. The ALS-ENABLE beamlines are supported in part by the NIH, National Institute of General Medical Sciences, grant P30 GM124169.

## ABBREVIATIONS

5’-dA: 5’-deoxyadenosine
5’-dA•: 5’-deoxyadenosyl radical
DFT: density functional theory
DT: sodium dithionite
*d3*-SAM: *S*-adenosyl-[*methyl*-^2^H3]methionine
EPR: electron paramagnetic resonance
HF: hyperfine
MIA: 3-methyl-2-indolic acid
NOC: nocathiacin
NOS: nosiheptide
PFL-AE: pyruvate formate-lyase activating enzyme
RiPP: ribosomally synthesized and post-translationally modified peptide
RMSD: root-mean-square deviation
RRE: RiPP recognition element
RS: radical *S*-adenosylmethionine
RSM: radical *S*-adenosylmethionine methylase
SAH: *S*-adenosylhomocysteine
SAM: *S*-adenosylmethionine
SOMO: singly occupied molecular orbital
SRC: side-ring closed product
ThzGlu: thiazolyl glutamate
TIM: triosephosphate isomerase.

## TOC Graphic

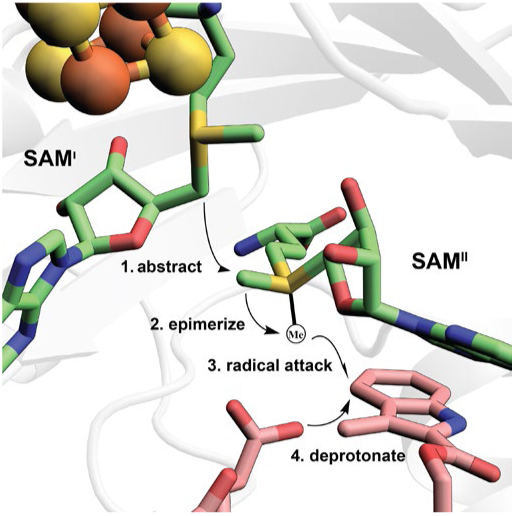

