## supporting information for "Structural and Spectroscopic Basis for Catalysis by a Class C Radical *S*-adenosylmethionine Methylase Involved in Nosiheptide/Nocathiacin Biosynthesis"

### TABLE of CONTENTS

**Page 2.** Table of contents

**Page 3.** Figure S1. Chemical structures of nosiheptide and nocathiacin I

**Page 4.** Figure S2. NocN reaction on compound **1** using SAM and *$d_3$*-SAM

**Page 5.** Figure S3. NosN reaction on compound **3** using *$d_7$*-SAM

**Page 6.** Figure S4. Comparison of the overall structures of NocN and HemN

**Page 7.** Figure S5. Comparison of the RS domains of NocN, HemN, and PFL-AE

**Page 8.** Figure S6. Interaction networks between two SAM molecules and NocN or HemN

**Page 9.** Figure S7. π-cation-π stacking by the side chain of R230

**Page 10.** Figure S8. Sequence alignment of nine HemN-like RS enzymes

**Page 11.** Figure S9. Conformations of the methionine moieties of $SAM^{II}$ in NocN/HemN

**Pages 12.** Figure S10. Binding poses of two AzaSAM molecules

**Pages 13.** Figure S11. Chemical structures of SRC and full-length SRC

**Pages 14.** Figure S12. SRC2 in the C-terminal domain of NocN

**Pages 15.** Figure S13. Hypothetical conformational change upon hydrogen abstraction.

**Pages 16.** Figure S14. DFT calculation of sulfonium epimerization

**Pages 17.** Figure S15. Turnover values of NosN mutants

**Pages 18.** Figure S16. Chemical structures of SAM adducts

**Pages 19.** Figure S17. Y202 and Y251 of NosN are not the base that deprotonates the radical intermediate

**Pages 20.** Figure S18. The carboxyl group of ThzGlu is the base that deprotonates the radical intermediate

**Pages 21.** Figure S19. NosN reaction using compound **6** using SAM, *$d_3$*SAM, and $^{13}C$-SAM

**Pages 22.** Figure S20. Time course EPR signals of compound **1**

**Pages 23.** Figure S21. Time course EPR signals of compound **5**

**Pages 24.** Figure S22. Spin density distribution of Compound **1** radical and Compound **8** radical

**Pages 25-94.** Experimental procedure

**Pages 95-96.** References

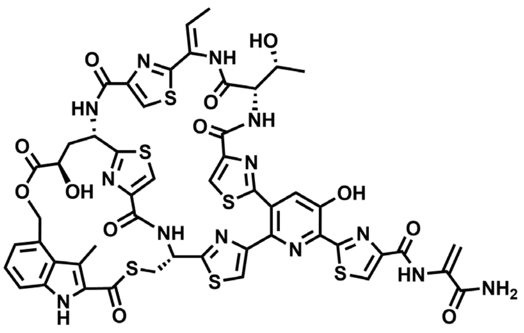

**nosiheptide**

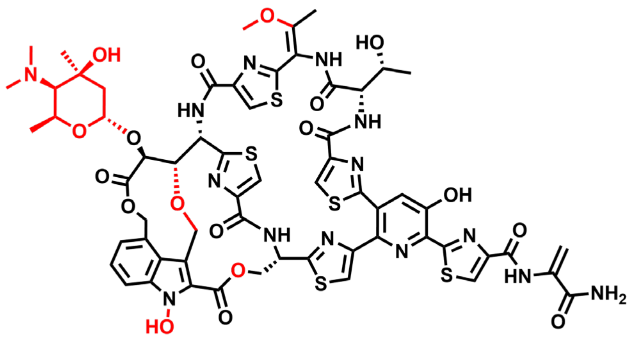

**nocathiacin I**

**Figure S1**. Chemical structures of nosiheptide and nocathiacin I. The scaffolds of both thiopeptide natural products are shown in black. Nocathiacin I has further decorations, including hydroxylations, methylations, oxidations, and glycosylation, which are highlighted in red.

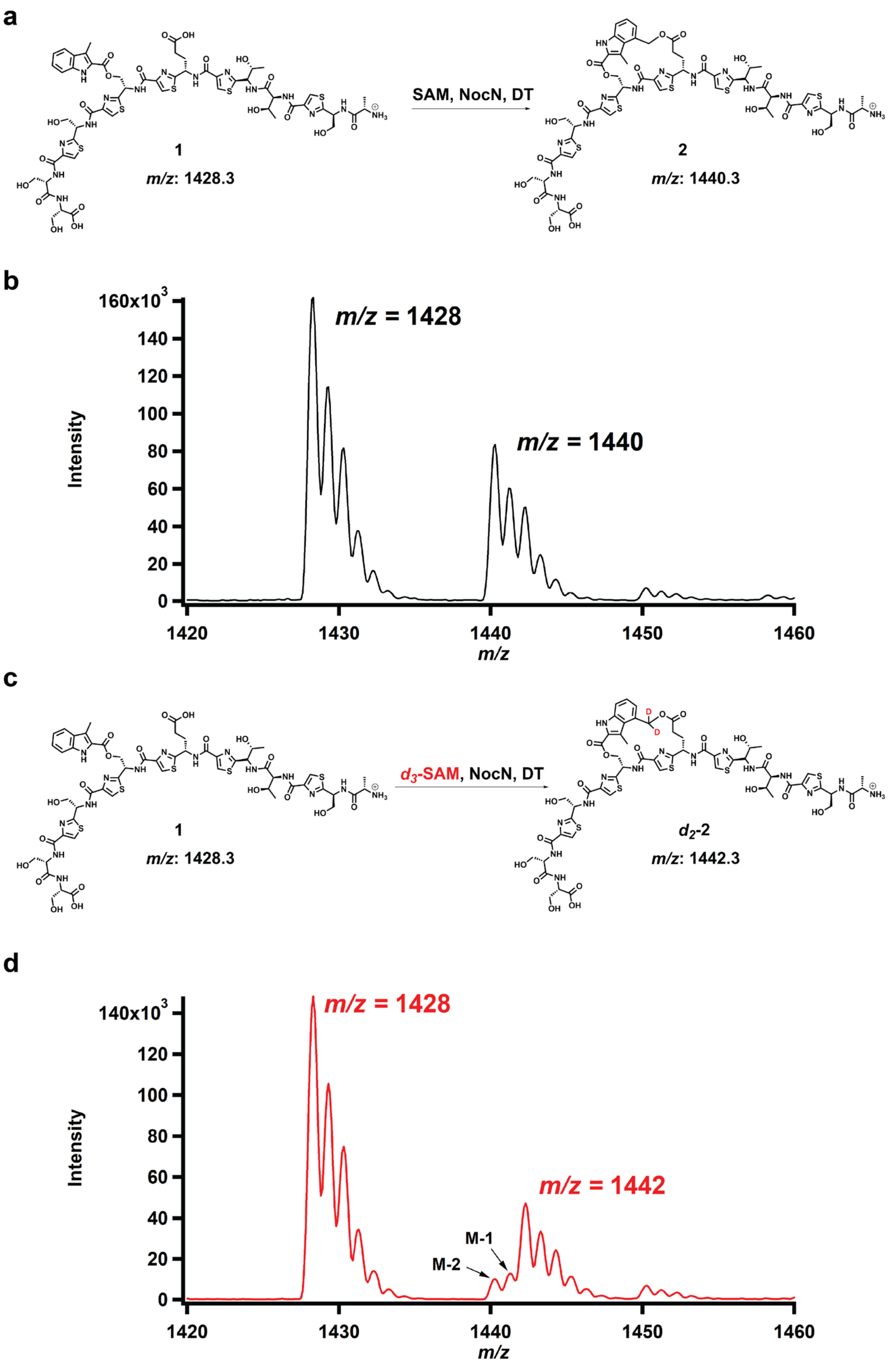


**Figure S2**. NocN reaction on compound **1** using SAM (**a**) and *$d_3$*-SAM (**c**) and *m/z* of the corresponding ring-closed product of NocN reactions using SAM (**b**) and *$d_3$*-SAM (**d**). DT stands for sodium dithionite which is the reductant used for the reaction.

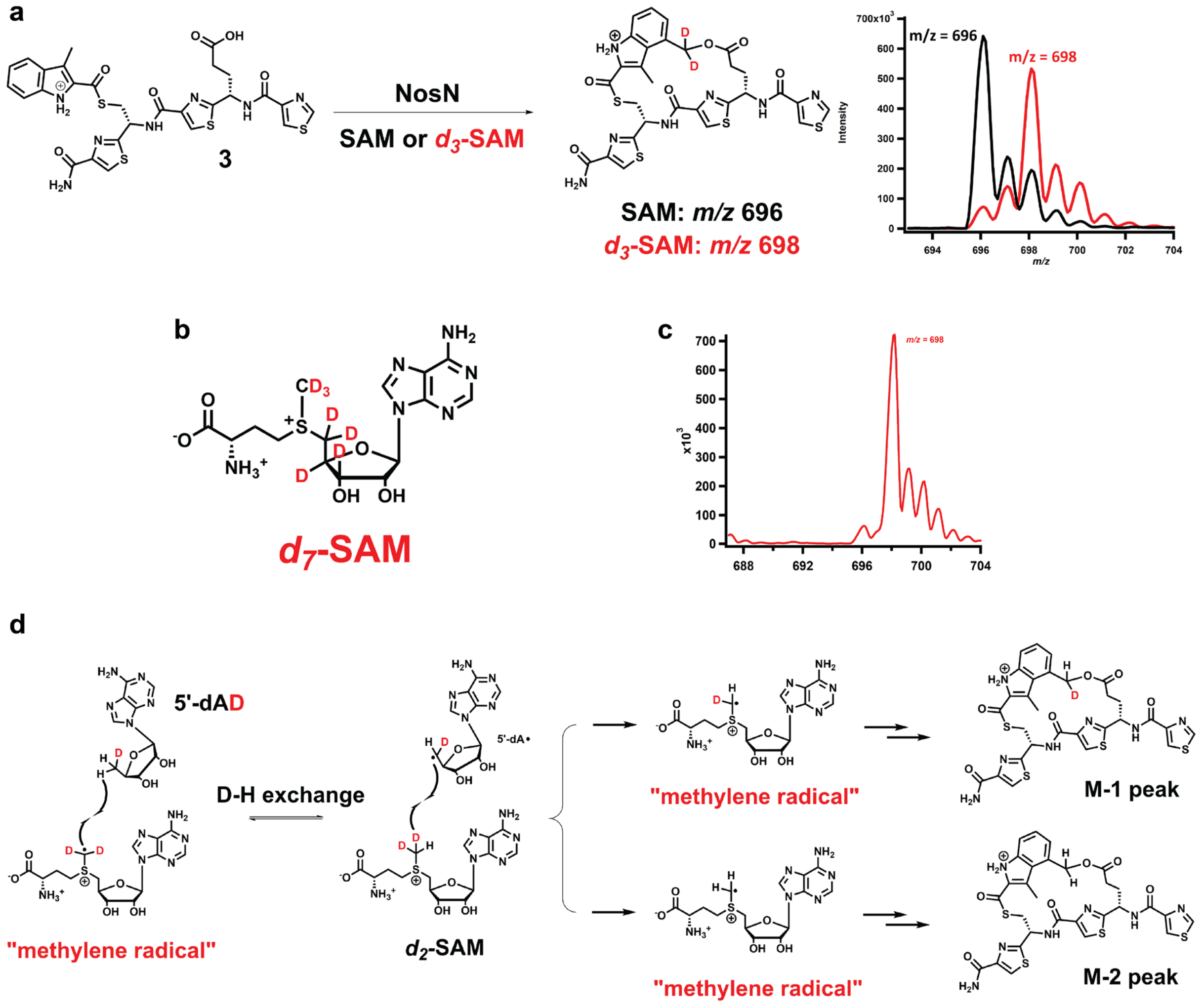


**Figure S3**. **a.** NosN reaction on peptide **3** using SAM or *d*$_3$-SAM. *d*$_3$-SAM gave an M-1 isotope peak that is more intense than the M-2 peak; **b.** chemical structure of *d*$_7$-SAM; **c.** In the NosN reaction on peptide **3** using *d*$_7$-SAM, the M-1 and M-2 peaks disappeared; **d.** A proposed mechanism of deuterium exchange.

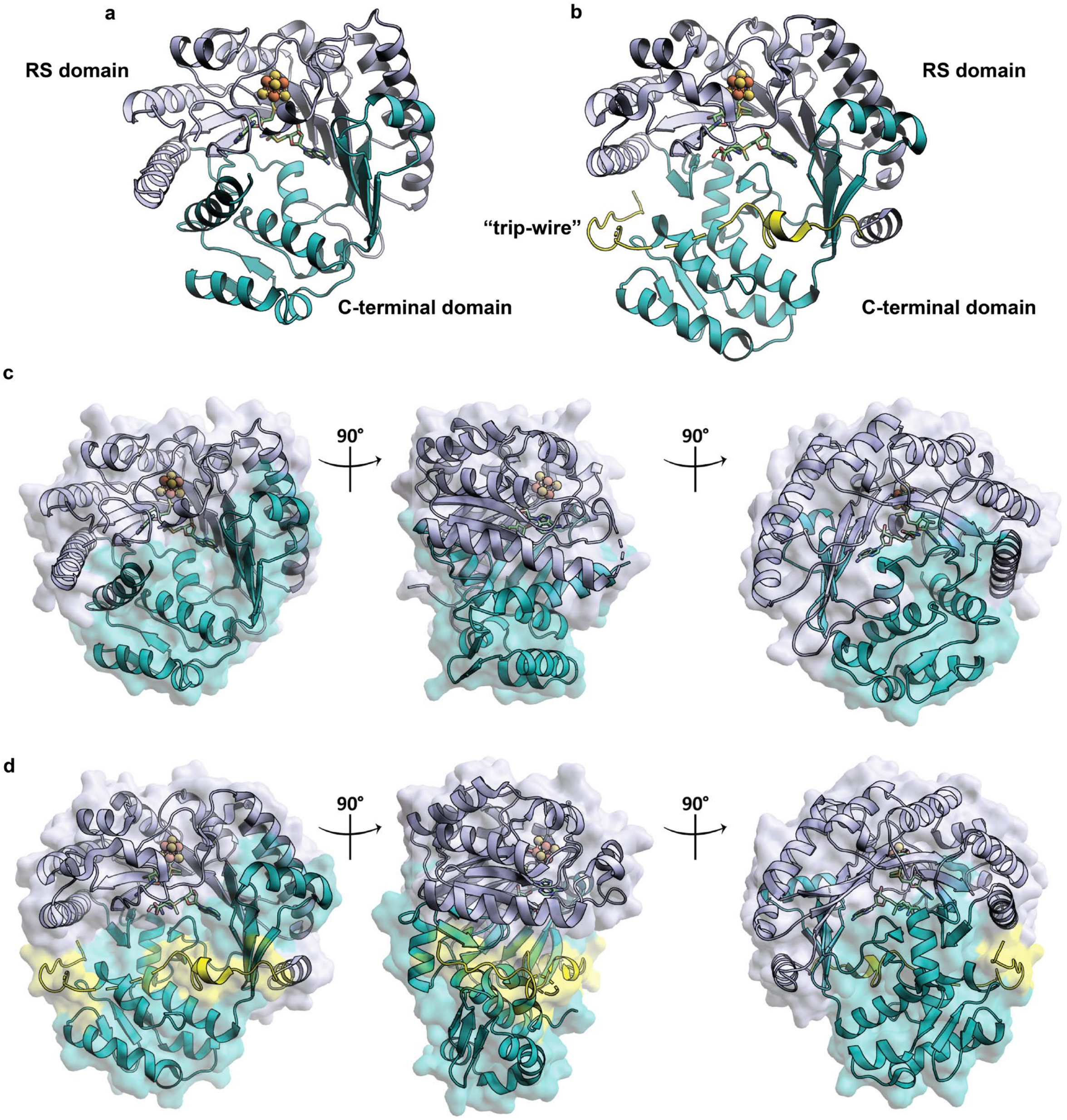


**Figure S4**. Ribbon diagrams of NocN (**a**) and HemN (PDB ID: 1OLT) (**b**). The radical SAM domain is colored light blue and the C-terminal domain is colored in teal. The N-terminal "trip-wire" domain of HemN is colored in yellow; **c.** Surface view of NocN; **d.** Surface view of HemN.

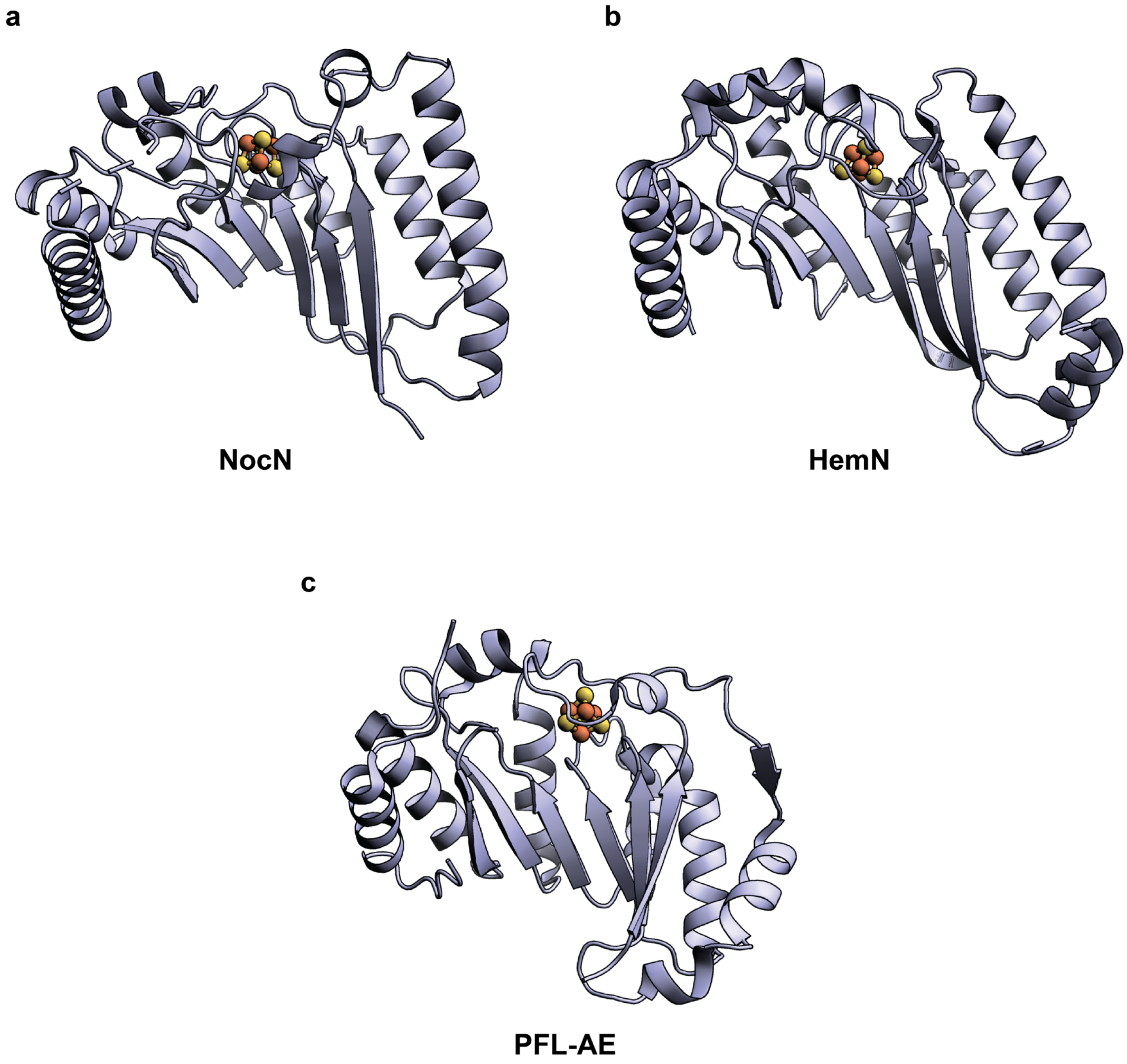


**Figure S5.** Ribbon diagrams of radical SAM domains of NocN (**a**), HemN (PDB ID: 1OLT) (**b**) and PFL-AE (PDB ID: 3C8F) (**c**). The RS domains of NocN and HemN show very similar structures with a loose curvature of the barrel to accommodate two simultaneously bound SAM molecules.

**a**

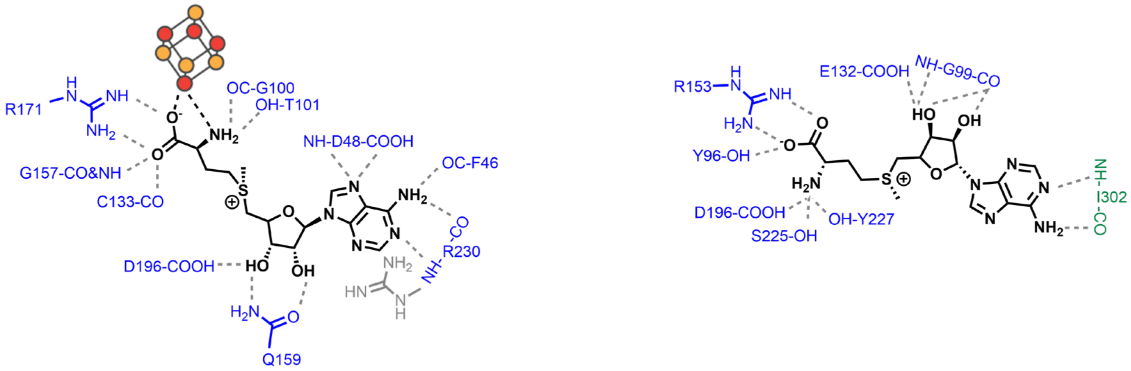


**NocN**

**b**

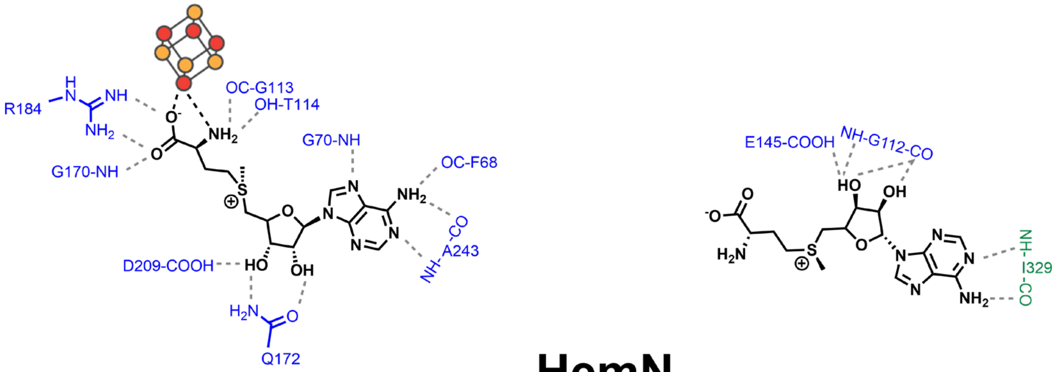


**HemN**

**Figure S6. a.** Interaction network between SAM$^{I}$/SAM$^{II}$ and NocN; **b.** Interaction network between SAM$^{I}$/SAM$^{II}$ and HemN (PDB ID: 1OLT). The residues from the RS domain of both enzymes are shown in blue. The residues from the C-terminal domain of both enzymes are shown in green.

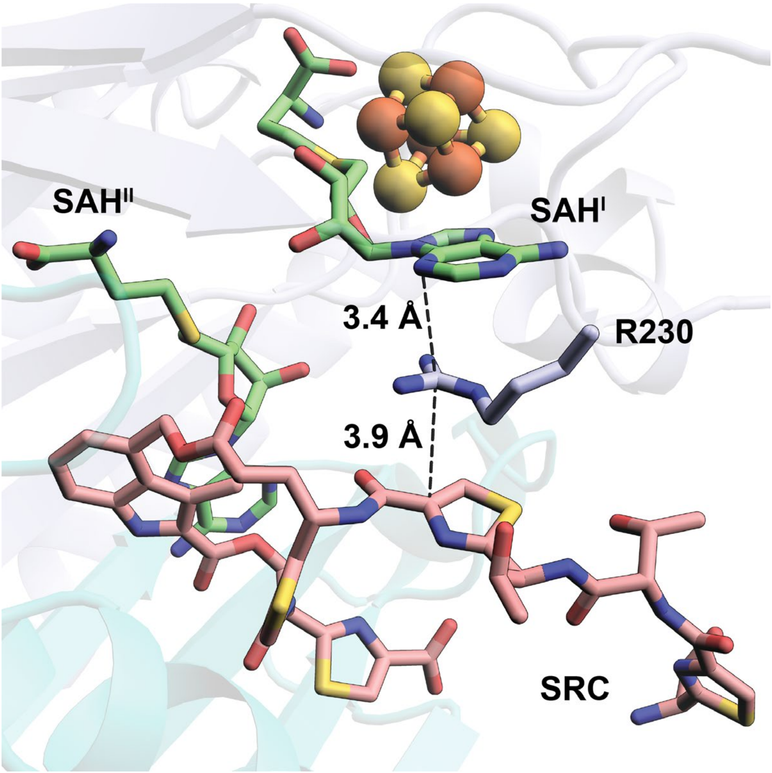


**Figure S7.** The guanidium side chain of R230 forms a π-cation-π stacking interaction with the adenine ring of SAH$^{I}$ and a thiazole ring of the side-ring closed product (SRC) in the crystal structure of NocN with SAH and SRC bound.

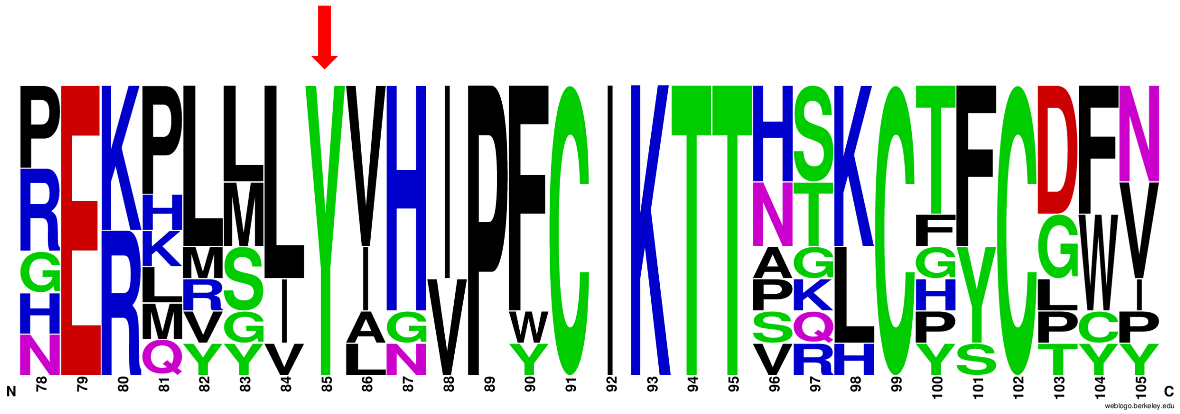


**Figure S8.** Sequence alignment of nine characterized HemN-like RS enzymes indicates that the tyrosine residue with a red arrow above (Tyr34 for NocN, or Tyr56 for HemN) is completely conserved. This sequence alignment is visualized in Weblogo format. Selected class C radical SAM methylases include: NocN (uniprot ID: E5DUI5), NosN (uniprot ID: C6FX53), C10P (uniprot ID: B1KQP7), ChuW (uniprot ID: A0A384LP51), HemN (uniprot ID: P32131), HemW (uniprot ID: P52062), Jaw5 (uniprot ID: A0A060PWX2), MqnK (uniprot ID: Q7MAP2), TbtI (uniprot ID: D6Y4Z7).

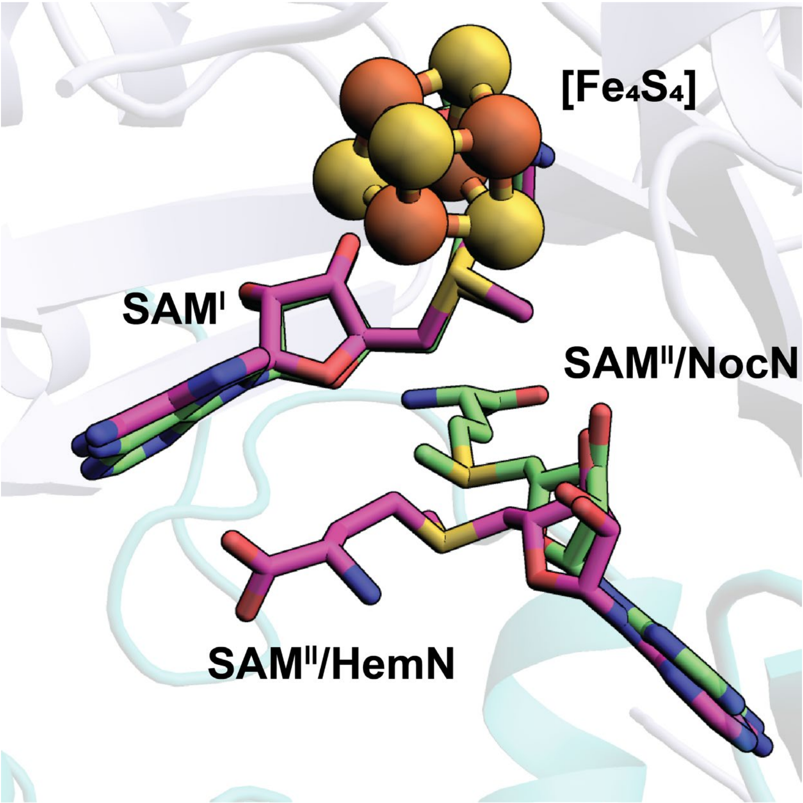


**Figure S9.** Overlay of NocN and HemN (PDB ID: 1OLT) shows different binding poses of the methionine moieties of SAM$^{II}$ from NocN and SAM$^{II}$ from HemN. It should be noted that SAM$^{II}$ in the HemN structure adopts an R,S configuration, whereas naturally occurring SAM has the S,S configuration.

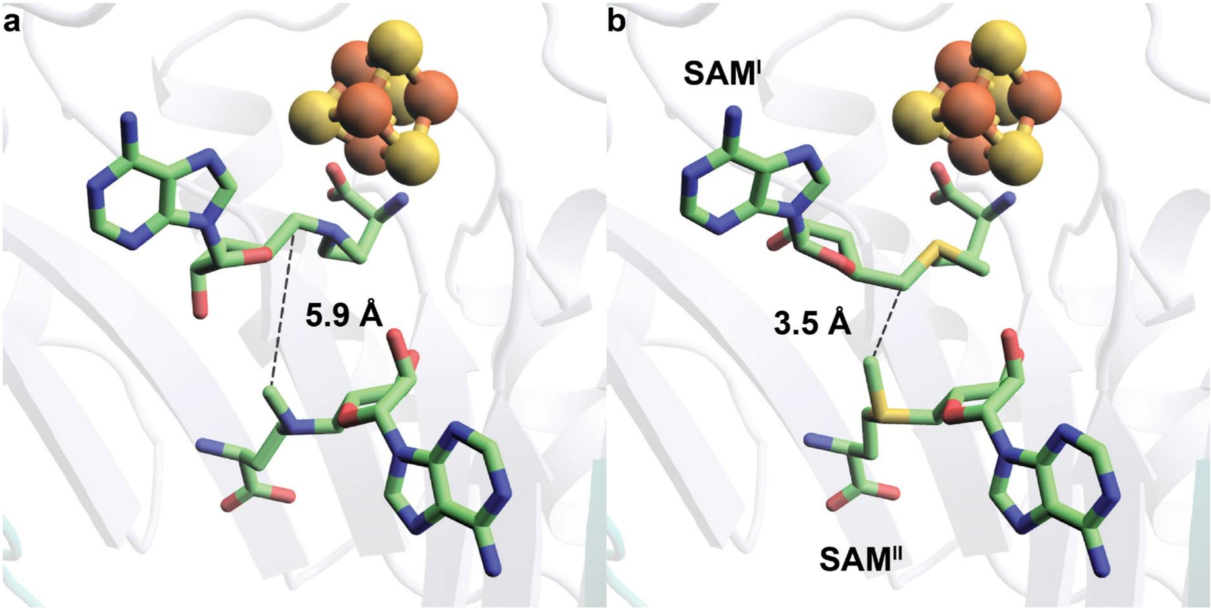


**Figure S10.** Comparison of the binding poses of two AzaSAM molecules with the two SAM molecules.

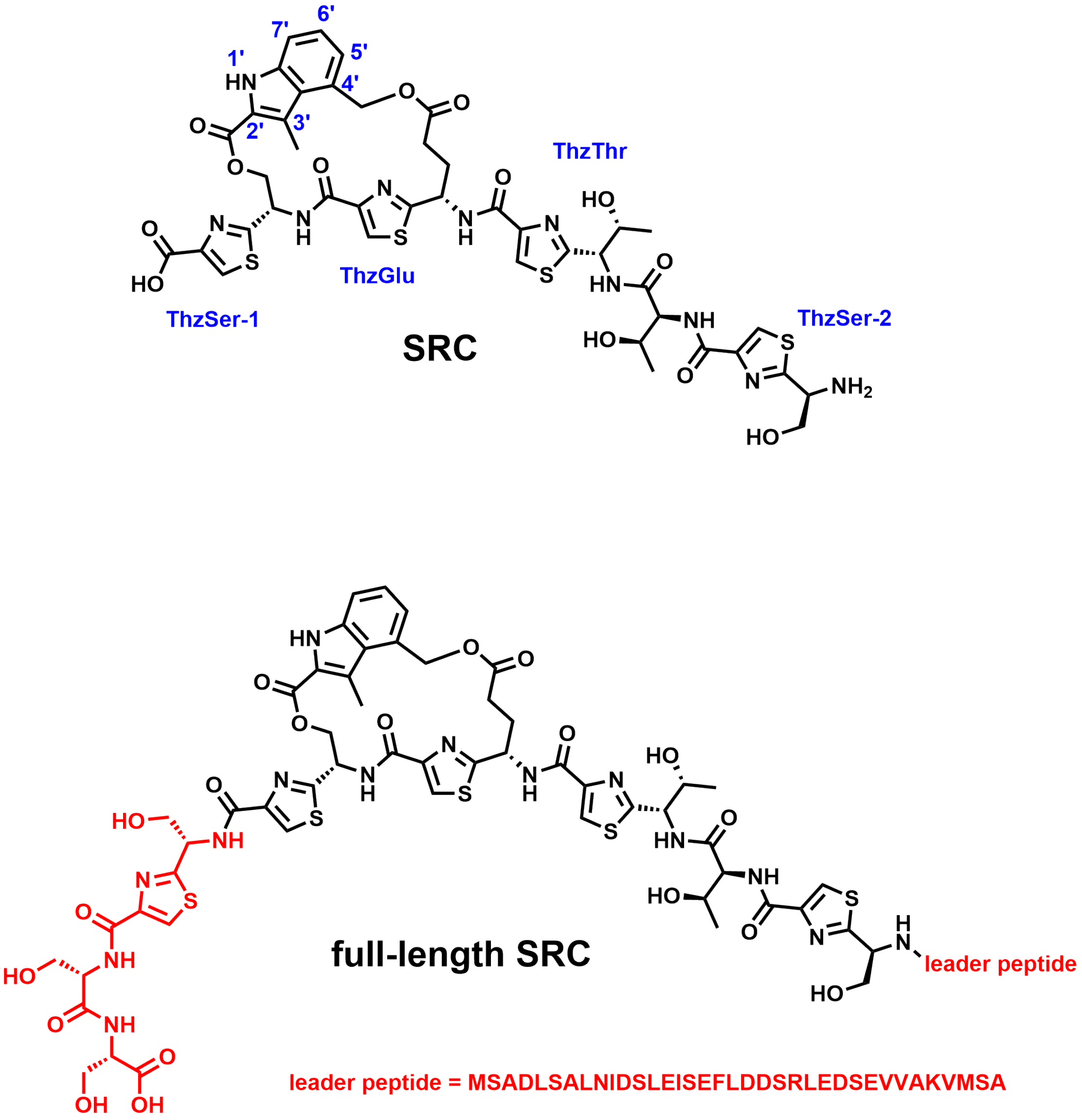


**Figure S11**. Chemical structures of SRC used for crystallization and full-length SRC. The residues colored in red in the structure of full-length SRC were removed to give the structure of SRC.

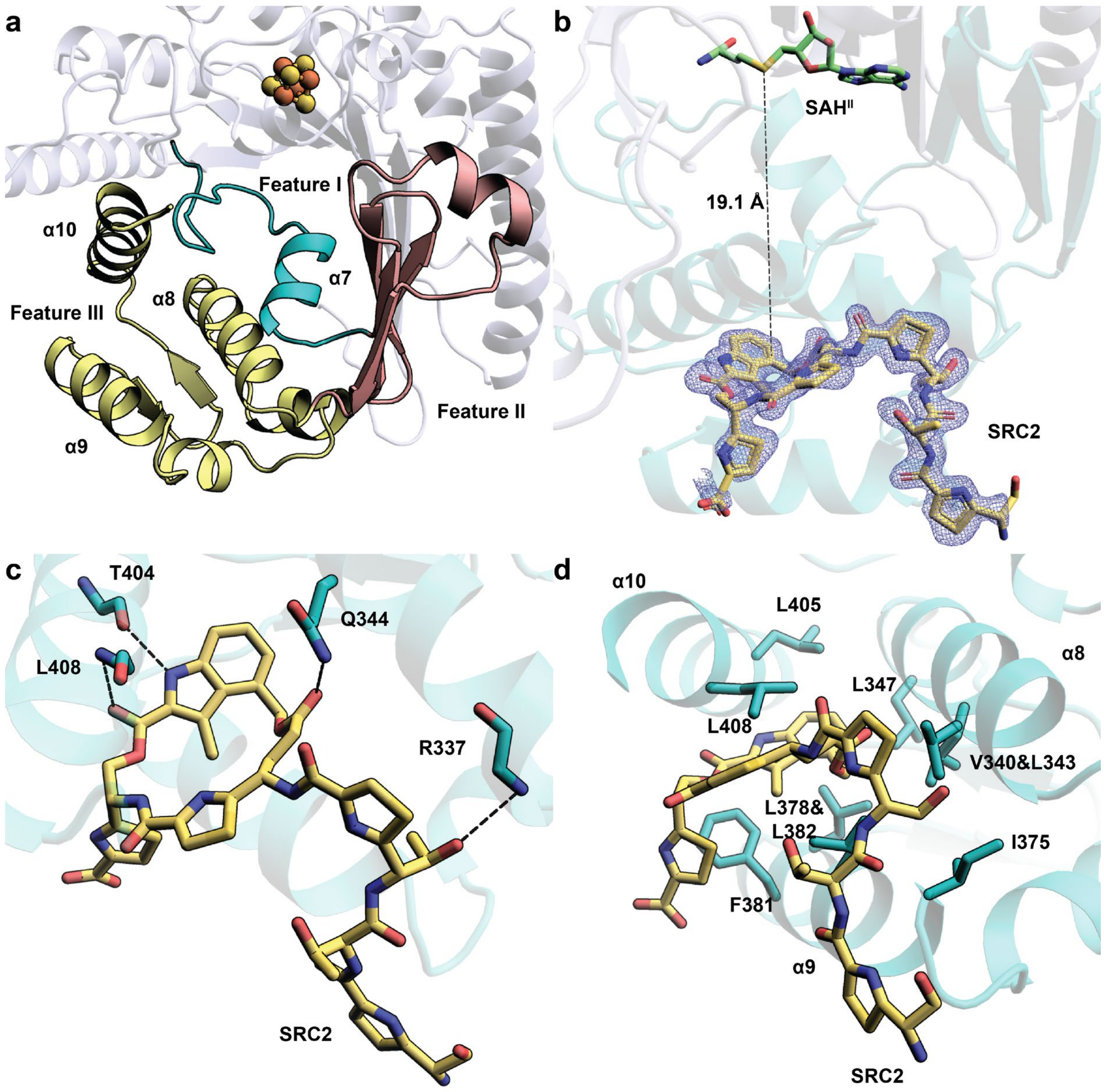


**Figure S12. a.** Three features of the C-terminal domain of NocN. Feature I is colored teal. Feature II is colored pink. Feature III is colored yellow; **b.** Close-up of SAH$^{II}$ in the active site and SRC2 in the C-terminal domain of NocN structure. Fo-Fc omit electron density maps are shown for SRC2 (blue mesh, contoured at 3.0 σ). The distances between the sulfur atom of SAH$^{II}$ and the 4′-C of MIA are shown as a black dashed line; **c.** Direct H-bond interaction between NocN and SRC2; **d.** Hydrophobic residues from α8, α9, and α10 around SRC2.

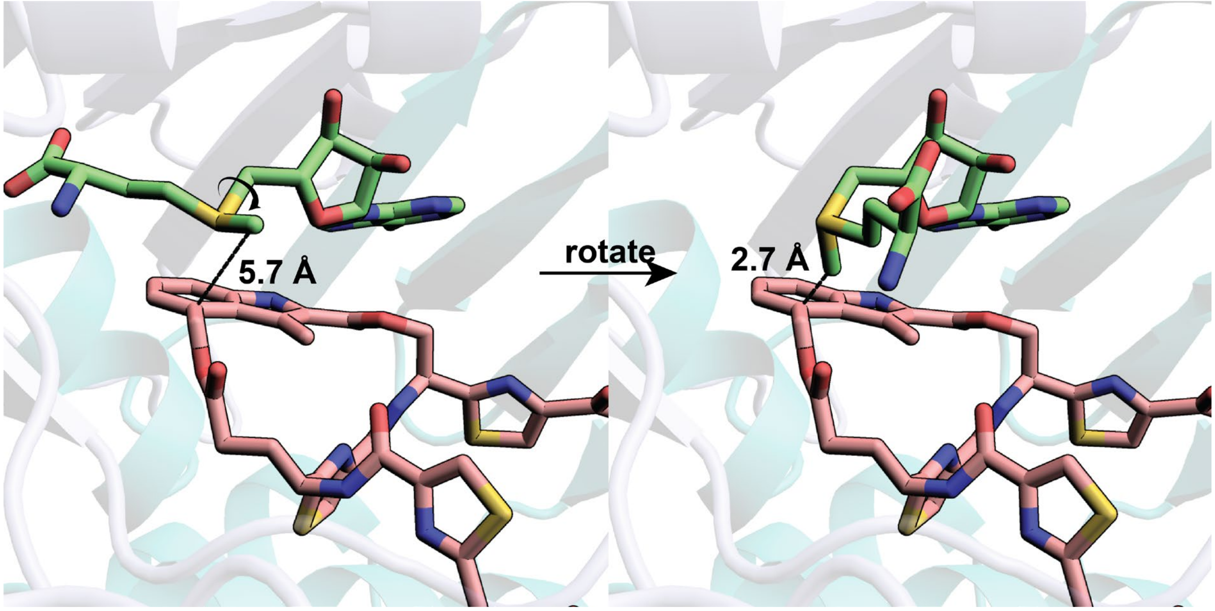


**Figure S13.** Upon hydrogen abstraction on the methyl group of $SAM^{II}$, rotation of the methionine moiety along 5′-C-S bond makes the methylene radical in a position for the attachment to 4′-C of MIA.

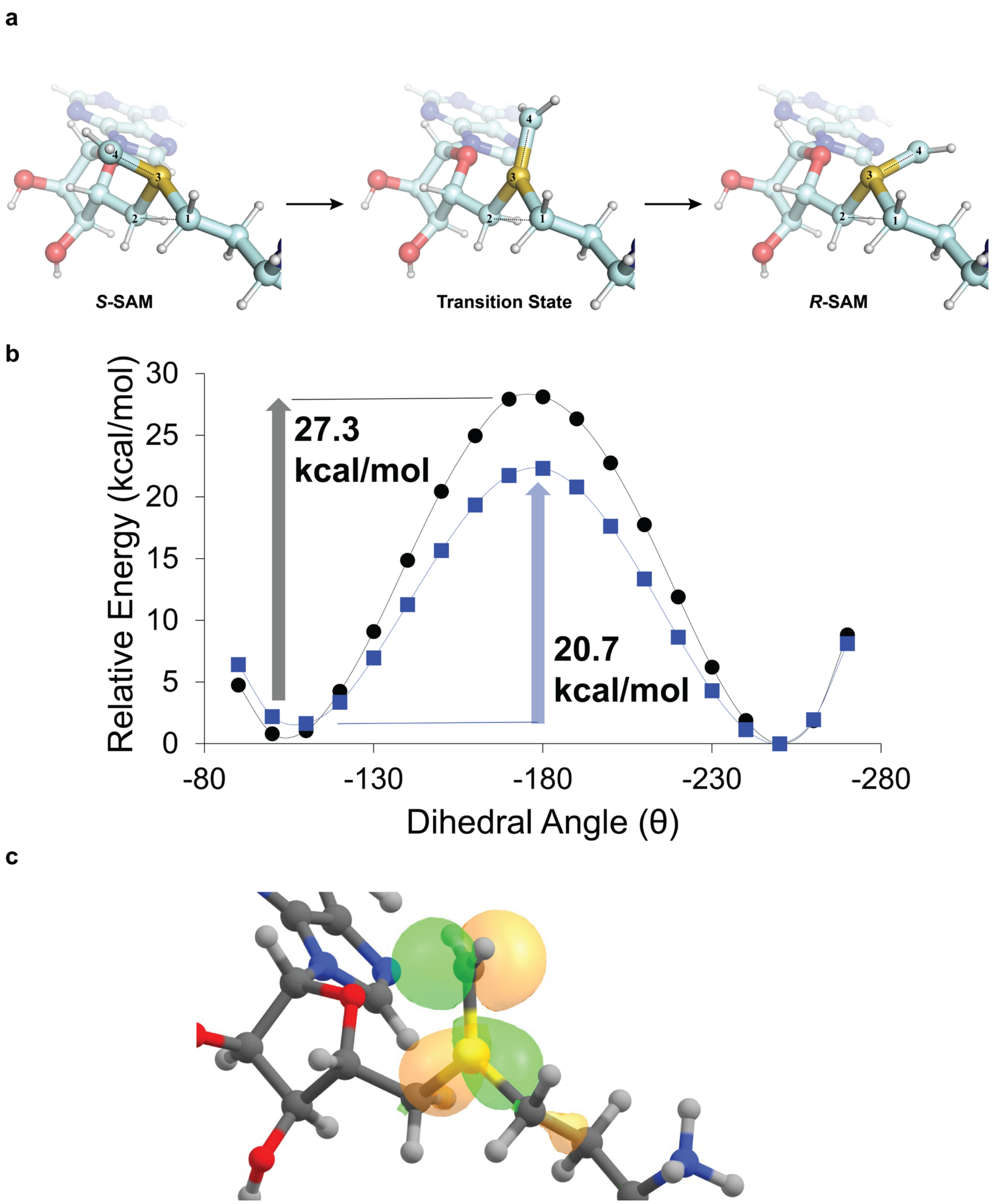


**Figure S14. a.** Ball and stick diagram showing epimerization of *S*-SAM methylene radical into *R*-SAM radical *via* a transition state in which the sulfonium atom has a planar structure; **b.** Energy chart of epimerization of SAM (black circles) or SAM methylene radical (blue squares); **c.** Singly occupied molecular orbital (SOMO) of the SAM methylene radical transition state (-180°) with + and – lobes as yellow and green, respectively.

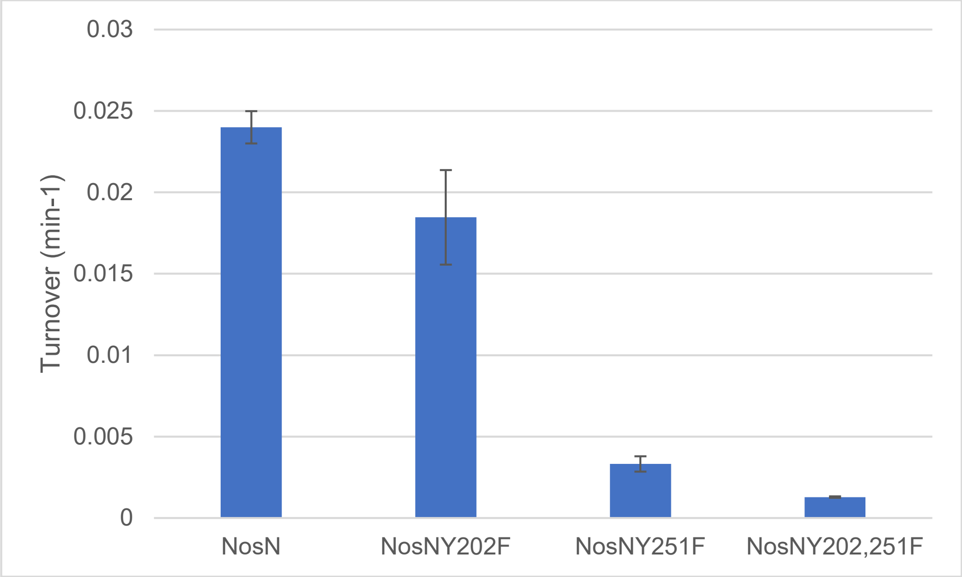


**Figure S15.** Turnover values of NosN (0.024 ± 0.00099 $min^{-1}$), Y202F (0.018 ± 0.0029 $min^{-1}$), Y251F (0.0033 ± 0.00047 $min^{-1}$), and Y202/251F (0.0013 ± 0.000045 $min^{-1}$). Turnover values were calculated from initial rates determined from the linear region (2–10 min) of the time-course data in **Figure 4b** and normalized to an enzyme concentration of 50 μM.

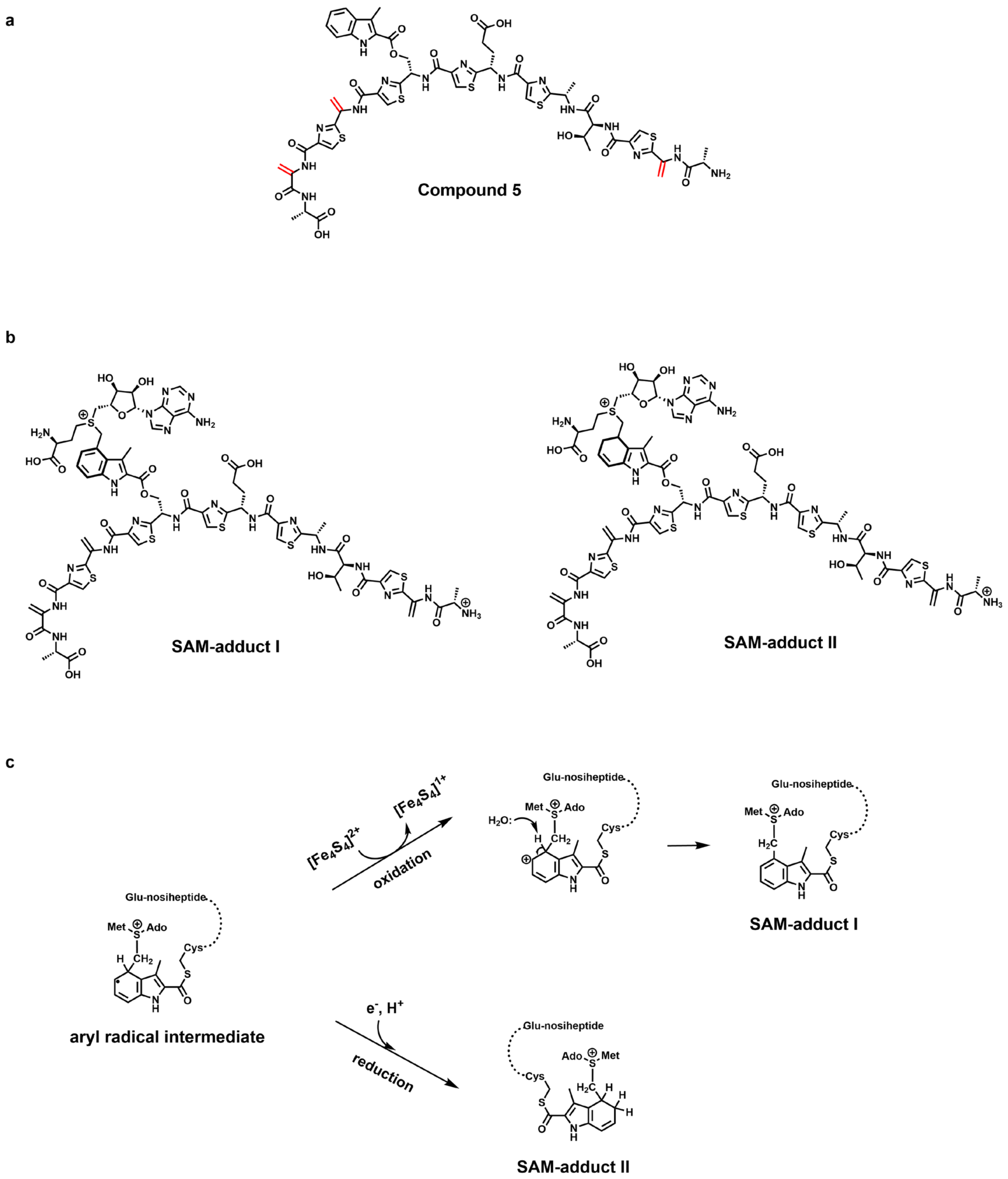


**Figure S16. a.** Chemical structure of compound **5**. The dehydroalanines are highlighted in red; **b.** Chemical structures of the two SAM adducts detected in the NosN reaction using compound **5** as a substrate; **c.** The mechanism of the formation of two SAM adducts.

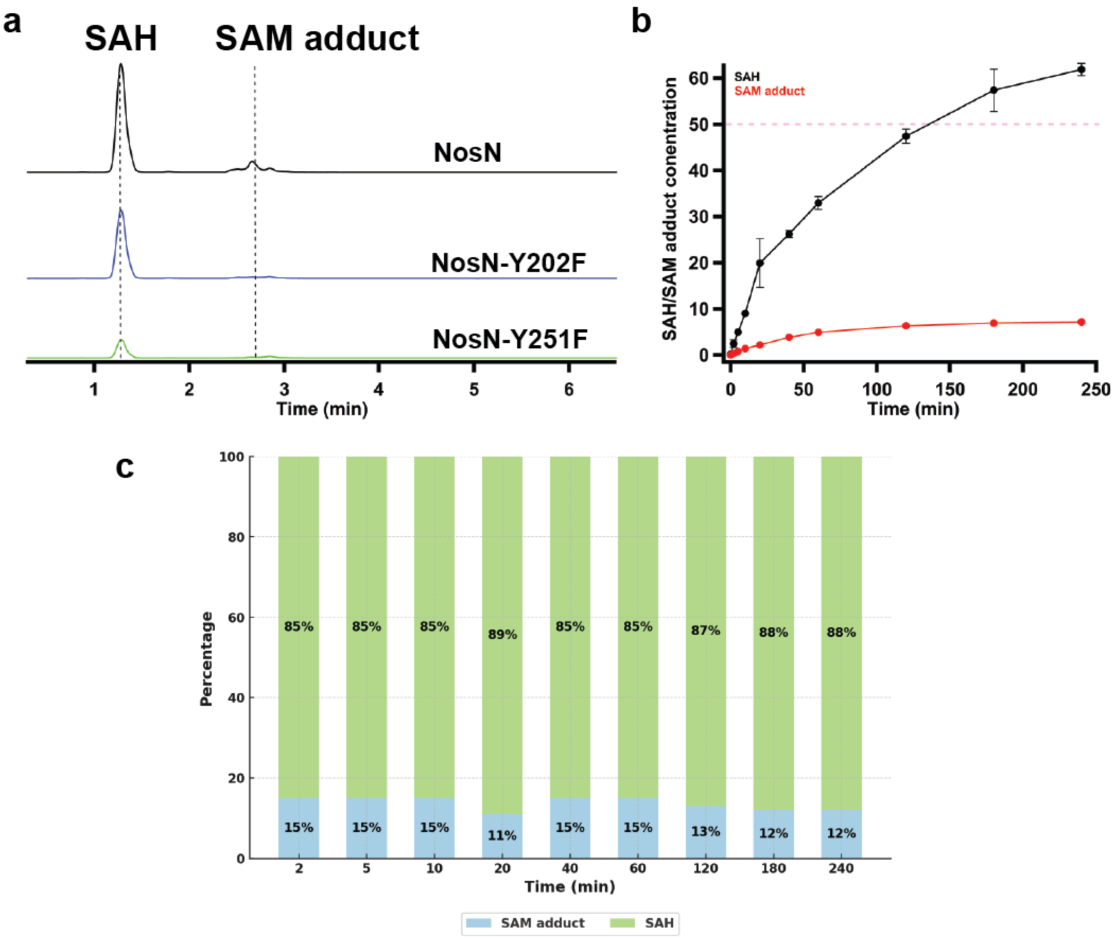


**Figure S17. a.** Multiple reactant monitoring (MRM) chromatogram (m/z 385 → 87.9) of SAH from reactions of NosN (black trace), NosN-Y202F (blue trace), and NosN-Y251F (green trace) using compound **3**. For each reaction, NosN or its variants (50 µM) was incubated with compound **3** (500 µM) in the presence of HEPES pH 7.5 (50 mM), SAM (1 mM), and tryptophan (150 µM) as an internal standard for 5 minutes. The reactions were initiated with the addition of sodium dithionite (2 mM). At 0, 2, 5, 10, 20, 40, 60, 120, 180, and 240 min, the reactions (18 µL) were quenched by the addition of sulfuric acid (9 µL, 100 mM). After quenching, methanol (27 µL) was added to precipitate the enzyme. The reaction mixtures were pelleted by centrifugation, and the supernatants were analyzed by LC-MS. The MRM chromatograms shown are from reactions quenched at 60 minutes. The peak at 1.2 min corresponds to authentic SAH. The peak at 2.7 min gives the signal of SAH, but the retention time indicates that this species is not authentic SAH. It is the SAH that is generated from mass spectroscopy-induced C-S cleavage that we reported in a previous study;[24] **b.** Quantification of SAH and SAM adduct in the NosN reaction with compound **3**; **c.** Ratio of SAM adduct-SAH in the NosN reaction using compound **3** as substrate.

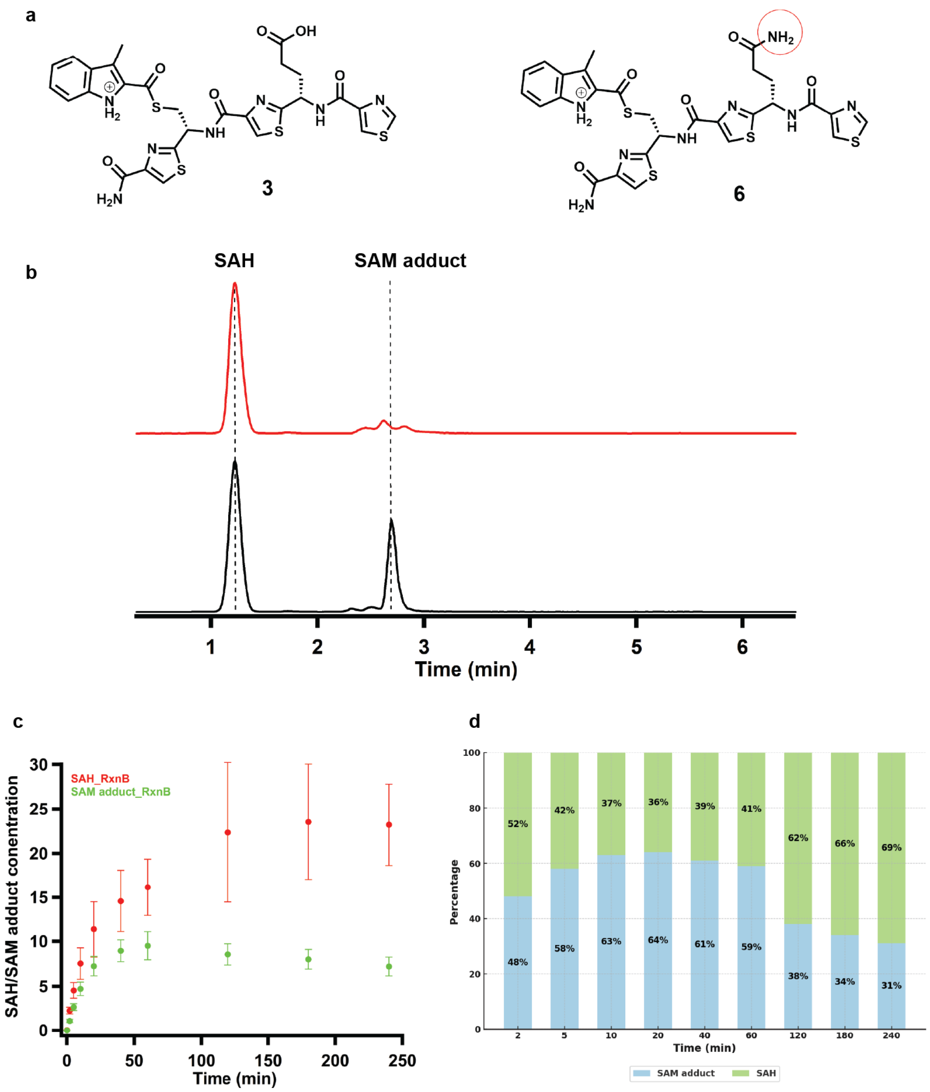


**Figure S18. a.** Chemical structure of compound **3** and compound **6**. Compound **6** has an amide instead free carboxylic acid as the side chain of ThzGlu; **b.** MRM chromatogram of SAH (m/z 385 → 87.9) from NosN reaction using compound **3** (red trace) and using compound **6** (black trace). The MRM chromatograms shown are from reactions quenched at 60 minutes. The peak at 1.2 min corresponds to authentic SAH. The peak at 2.7 min gives the signal of SAH, but the retention time indicates that this species is not authentic SAH. It is the SAH that is generated from mass spectroscopy-induced C-S cleavage that we reported in a previous study;[24] **c.** Quantification of SAH and SAM adduct in the NosN reaction with compound **6**. NosN (50 µM) was incubated with compound **6** (500 µM) in the presence of HEPES pH 7.5 (50 mM), SAM (1 mM), and tryptophan (150 µM) as an internal standard for 5 minutes. The reactions were initiated with the addition of sodium dithionite (2 mM). At 0, 2, 5, 10, 20, 40, 60, 120, 180, and 240 min, the reactions (18 µL) were quenched by the addition of sulfuric acid (9 µL, 100 mM). After quenching, methanol (27 µL) was added to precipitate the enzyme. The reaction mixtures were pelleted by centrifugation, and the supernatants were analyzed by LC-MS; **d.** Ratio of SAM adduct-SAH in the NosN reaction using compound **6** as substrate.

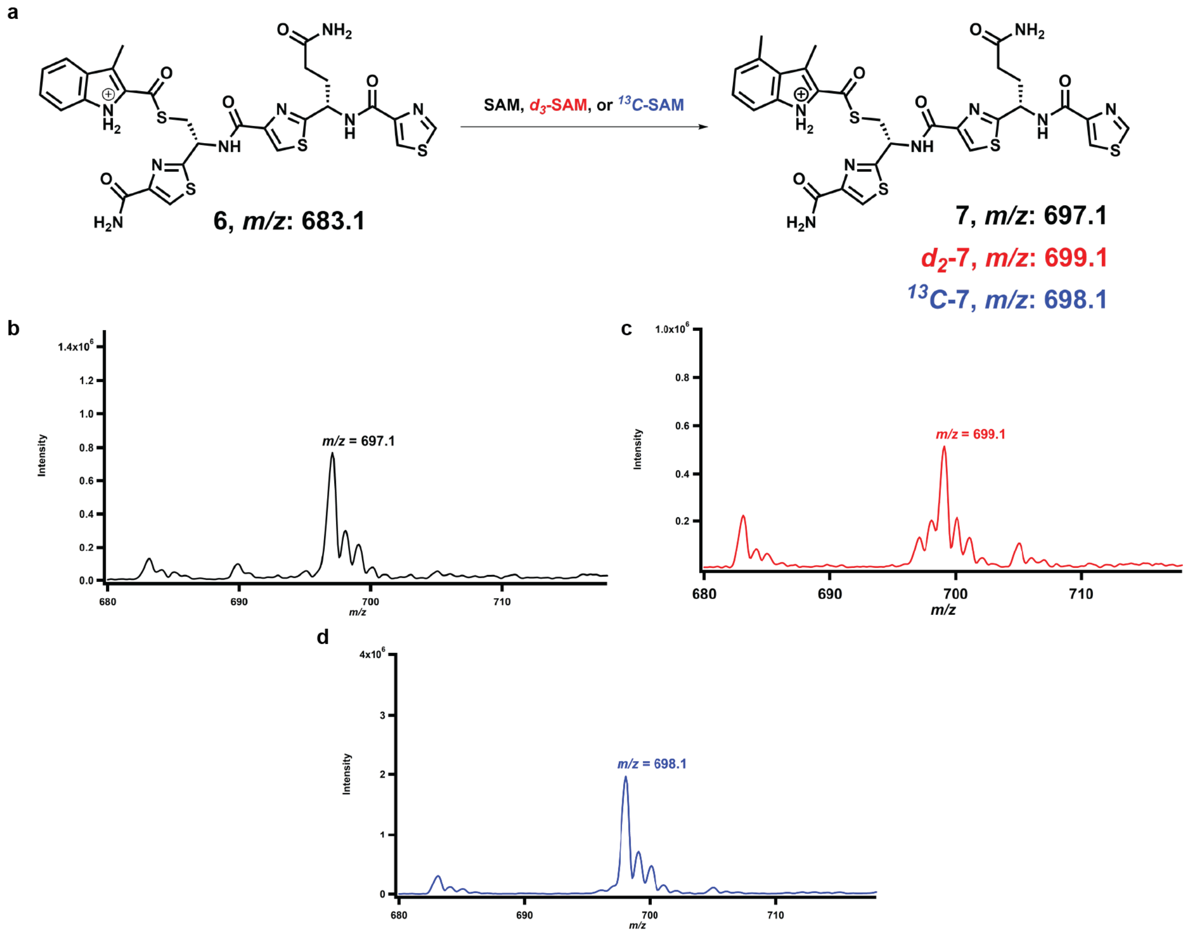


**Figure S19. a.** Chemical structure and *m/z* of compound **6** and its methylated product; **b.** *m/z* of the methylated product of compound **6** using SAM; **c.** *m/z* of the methylated product of compound **6** using $d_3$SAM; **d.** *m/z* of the methylated product of compound **6** using [*methyl*-$^{13}C$]-SAM ($^{13}C$-SAM). NosN (100 µM) was incubated with compound **6** (300 µM) in the presence of HEPES pH 7.5 (50 mM) and SAM/$d_3$SAM/$^{13}C$-SAM (1 mM) for 5 minutes. The reactions were initiated with the addition of sodium dithionite (2 mM) and incubated for 3 hours. The reactions (50 µL) were quenched by the addition of sulfuric acid (25 µL, 100 mM). After quenched, methanol (75 µL) was added to precipitate the enzyme. The reaction mixtures were pelleted by centrifugation, and the supernatants were analyzed by LC-MS.

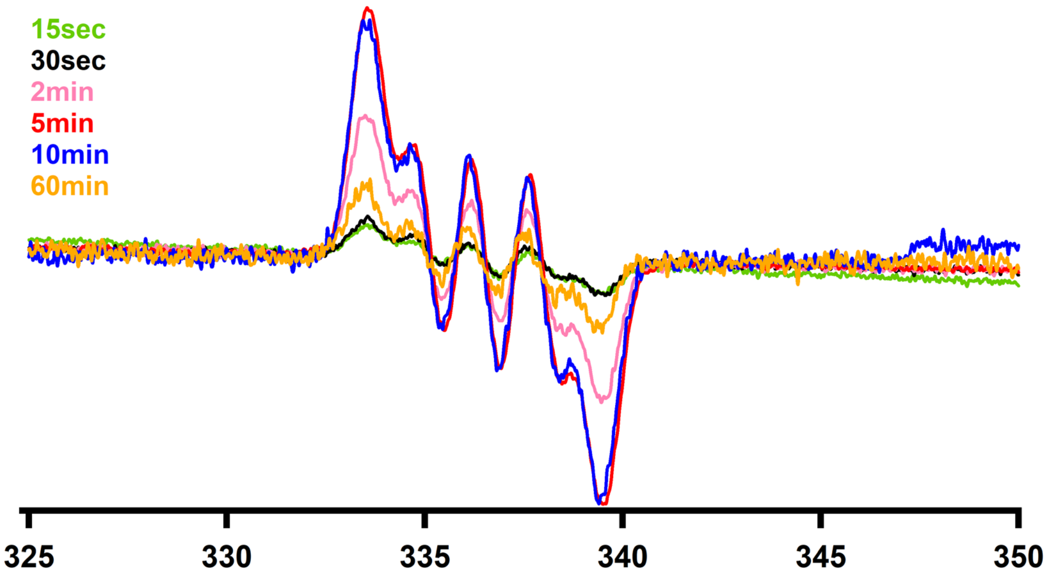


**Figure S20**. Time course EPR signals of compound **1**. The samples contain 100 mM Hepes pH 7.5, 10% glycerol, 550 μM NosN, 3 mM SAM, 1.5 mM compound **1**, and 4 mM sodium dithionite. The spectra were collected at 80K with an attenuation of 20 dB for four scans.

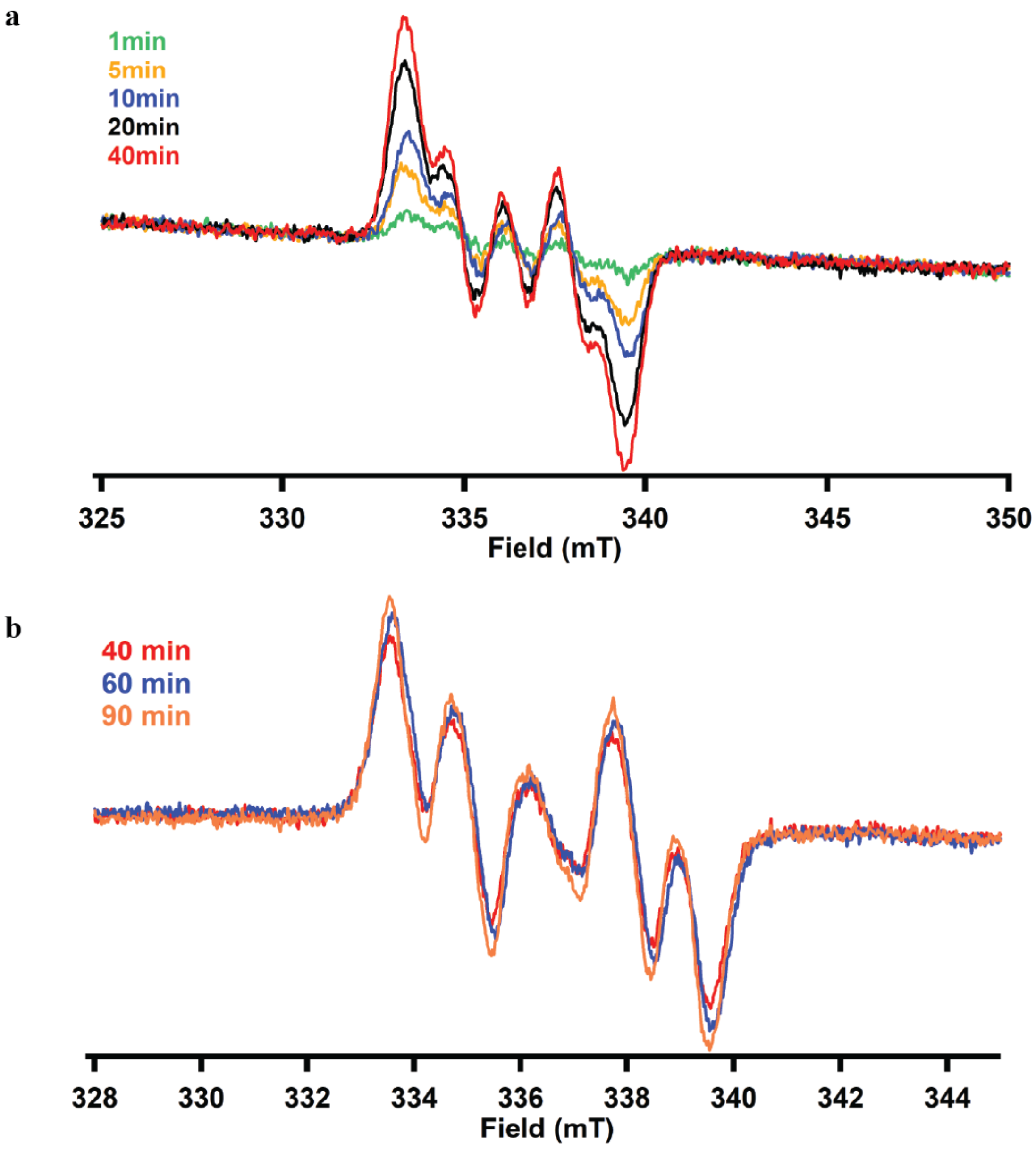


**Figure S21**. **A.** Time course EPR signals of compound **5** up to 40min. The samples contain 100 mM Hepes pH 7.5, 10% glycerol, 550 μM NosN, 3 mM SAM, 1.5 mM compound **5**, and 4 mM sodium dithionite. The spectra were collected at 80K with an attenuation of 20 dB for four scans; **B.** Time course EPR signals of compound **5** at 40min, 60min, and 90min. The samples contain 400 mM Hepes pH 7.5, 10% glycerol, 250 μM NosN, 2 mM SAM, 1 mM compound **5**, and 2 mM sodium dithionite. The spectra were collected at 60K with attenuation of 40 dB for four scans.

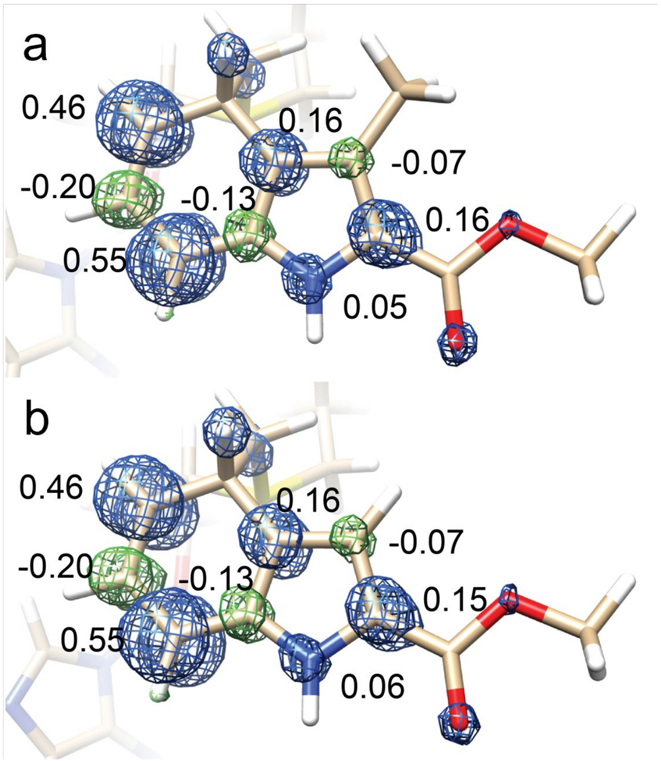


**Figure S22**. Spin density distribution in Compound **1** (**a**) and Compound **8** (**b**). Positive spin densities are depicted at 0.004 level; positive and negative densities are depicted in blue and green mesh respectively. Numbers represent relevant Mulliken spin populations on the ring structures.

### Experimental Procedures

**Materials.** All commercial materials were used as received unless otherwise noted. *N*-(2-Hydroxyethyl)-piperazine-*N*'-(2-ethanesulfonic acid) (HEPES) was were purchased from Fisher Scientific. Imidazole was purchased from J. T. Baker Chemical Co. Potassium chloride and glycerol were purchased from EMD Chemicals. 2-Mercaptoethanol, and *S*-adenosylhomocysteine (SAH) were purchased from MilliporeSigma. Kanamycin, ampicillin, Isopropyl β-D-1-thiogalactopyranoside (IPTG), and tris(2-carboxyethyl)phosphine (TCEP) were purchased from Gold Biotechnology. Ni-NTA resin was purchased from Qiagen. Plasmid DNA isolation kits were purchased from Macherey-Nagel (Dueren, Germany). Organic solvents, including tetrahydrofuran, dichloromethane, acetonitrile, and DMF were obtained from a JC Meyer solvent dispensing system and used without further purification. SiliaFlash 60 silica gel (230-400 mess) for flash chromatography was obtained from Silicycle Inc. SAM was synthesized and purified as described previously.[1] Reagents for peptide coupling and solid phase peptide synthesis were purchased from Chem-impex. All other chemicals were from Ambeed, Sigma-Aldrich, or Chem-impex. 3-methyl-2-indolic acid was purchased from eNovation Chemical LLC.

**General Methods.** UV-visible spectra were recorded on a Cary 50 spectrometer from Varian (Agilent Technologies, Santa Clara, CA) using the WinUV software package to control the instrument. High resolution mass spectrometry (HRMS) was conducted on a Thermo Scientific Vanquish UPLC in line with a Q Exactive HF-X hybrid quadrupole-Orbitrap mass spectrometer. Data were collected and processed using Thermo Scientific Xcalibur 4.2.47. NMR spectra of all compounds were collected on a Bruker AV-3-HD-500 instrument or a Bruker AVANCE NanoBay NEO-400 (NANO-2) instrument. Purifications of peptides from solid phase peptide synthesis were performed on an Agilent 1260 Infinity II Preparative HPLC.

### 1. Overexpression and purification of NocN

Expression and purification of NosN were performed following the previously reported procedure.[2] The gene for *nocN* from *Nocardia* sp. ATCC 202099 was codon-optimized for expression in *E. coli* (sequence below) and synthesized by GeneArt (ThermoFisher, Carlsbad, CA). The gene was subcloned into pSUMO vector using *Nde*I and *Xho*I restriction sites. The resulting plasmid was co-transformed with *E. coli* BL21 (DE3) competent cells containing plasmid pDB1282. Starting from a single colony, a 200 mL starter culture containing 50 mg/L kanamycin and 100 mg/L ampicillin was shaken at 37 °C and 250 rpm for 12 h. 10 mL of the starter culture was used to inoculate 4 L of M9 medium containing 50 mg/L kanamycin and 100 mg/L ampicillin and incubated at 37 °C and 180 rpm. Expression of the genes encoded on plasmid pDB1282 was induced at an $OD_{600}$ of 0.3 with 0.2 % arabinose (8 g). At $OD_{600}$ of 0.6, the flasks were placed in an ice-water bath for 0.5 h. Once cooled, IPTG was added to a final concentration of 0.5 mM, and iron chloride was added to a final concentration of 25 µM. The cultures were incubated overnight for 18 h at 18 ºC with shaking at 180 rpm. The cells were harvested (~30g, from 4 × 4L growth), flash-frozen in liquid nitrogen, and stored at -80 ºC until use.

For purification, 30 g of frozen cell paste was resuspended in 150 mL of lysis buffer (50 mM HEPES, pH 7.5, 300 mM KCl, 10% (v/v) glycerol, and 10 mM BME) containing lysozyme (150 mg) and DNaseI (15 mg). Resuspended cells were incubated on ice and subjected to six sonic bursts (40% output) on a QSonica instrument in a Coy anaerobic chamber for 45 s each with 8 min intermittent pauses. The lysate was then centrifuged for 1 h at 45,000g at 4 ºC. The resulting supernatant was loaded onto Ni-NTA resin equilibrated in the lysis buffer. The resin was washed twice with 100 mL of the lysis buffer prior to elution of SUMO-NocN with elution buffer (50 mM HEPES, pH 7.5, 300 mM KCl, 250 mM imidazole, 10% (v/v) glycerol, and 10 mM BME). The pooled eluate was concentrated by ultracentrifugation using an Amicon Ultra centrifugal filter unit with a 10 kDa molecular weight cutoff membrane. The buffer of the eluate was exchanged into a cleavage buffer (50 mM HEPES, pH 7.5, 300 mM KCl, 250 mM imidazole, 10% (v/v) glycerol, and 10 mM BME) using PD-10 column (GE Healthcare). Ulp1 (0.5 mL, 8 mg/mL, 4 mg) was added to the resulting exchanged SUMO-NocN solution and incubated on ice overnight to remove the SUMO tag. The cleavage reaction mixture was applied on a Ni-NTA column equilibrated with cleavage buffer. Flow-through was collected, pooled, and concentrated using an Amicon Ultra centrifugal filter unit with a 10 kDa molecular weight cutoff membrane. The resulting NocN protein was further purified by size-exclusion chromatography on a HiPrep 26/60 S200 column with an isocratic method using S200 buffer (50 mM HEPES, pH 7.5, 400 mM KCl, 10% glycerol, and 10 mM TCEP) as the mobile phase. Fractions indicative of monomeric NocN were pooled, concentrated, frozen by liquid nitrogen, and stored in liquid nitrogen.

Sequence of the codon-optimized NocN gene as supplied by GeneArt:

5′-

CATATGAGCACCGCAGTTAGCCTGAGCAGCCTGGTTGATGTTGTTCCGGCACCGGCA
CCGAATCTGCCGAGCAGCACCGATGAACAGCACATGCTGATGCTGTATGTTCATGTT
CCGTTTTGTCATAGCAAATGCACCTTTTGTGATTGGGTTCAAGCAATTCCGACCAAA
GATCTGCTGCGTAAACCGGAAGATAGCGTTCGTAAAAACTATATTCGTGCACTGGTG
ACCGAAATTGAAACCCGTGGTGCACAGCTGCGTGCAGCAGGTCAGGTTCCGTATGTT
GTTTATTGGGGTGGTGGCACCGCAAGCAGTCTGGATAATGCAGAAGCCGAAGCAAT
TTGGGGTGCACTGGATAGCGCATTTGATCTGAGCACCGTTGCGGAAGCAACCATTGA
ATGTAGTCCGGATACCGTTGATAAAGCCAAACTGGAATTTTTCCGTGGTCTGGGTTT

TAATCGTGTTAGCAGCGGTGTTCAGAGCTTTGATGATGCACGTCTGCGTCGTCTGGG

TCGTCGTCATACCGCAGGCGAAGCAGATCGTATTGTTCATCATGCACGTGAAGCAGG

TTTTGATGAAGTGAGCATTGATATTATGAGCGGCTTTCCGGATCAAGAACTGGATGA

ACTGCGTGCAACCGTGGAAAAAGCCGTTAGCCTGCCGCTGACACATCTGAGCCTGT

ATAGCTTTCGTCCGACACCGGGTACATTTATGCGTCGTAAACTGGCAGGCACCGAAA

AACGTGCATATCTGCGCAAACAGCAGGCACTGTTTACCGAAGCACGTCGTATGATTA

TTGATGCCGGTCTGCCGGAATATGCAAGCGGTTATTTTGGTCGTGTGAGCCCGTTTG

CAGCAATGTATTTTCAGTTACGTGCAGATACAGCCGGTTTTGGTAGCGGTGCAATTA

GCCTGGTGGATCGTCAGTTTCTGAGCCATGCAAAAGGTAAACTGCATGCCTATATTC

AGGATCCGCTGGCCTATGATATTGACGTTCCGGCAGGCCAGGATCGTGTTCTGGTTA

GCTTTCTGCAGGCAGGTCTGGCAATGTTTGATGGTGTTCTGCGTGAAGAATGGCGTG

TTTCAACCGGTACAGATCTGGATGAAGTTCTGACCCGTCCGAGCATTGCTCCGCTGG

CAGATTTTCTGCGTGGTCGTGGTCTGATTGAAGATGAACGTGGTATTCGTCTGGATC

CGCGTCTGGCTGGTCTGACCCTGATTGAACTGGCATTTGAAATGGCAATGAGCCAGC

CGGAAAGCGCATAACTCGAG-3′

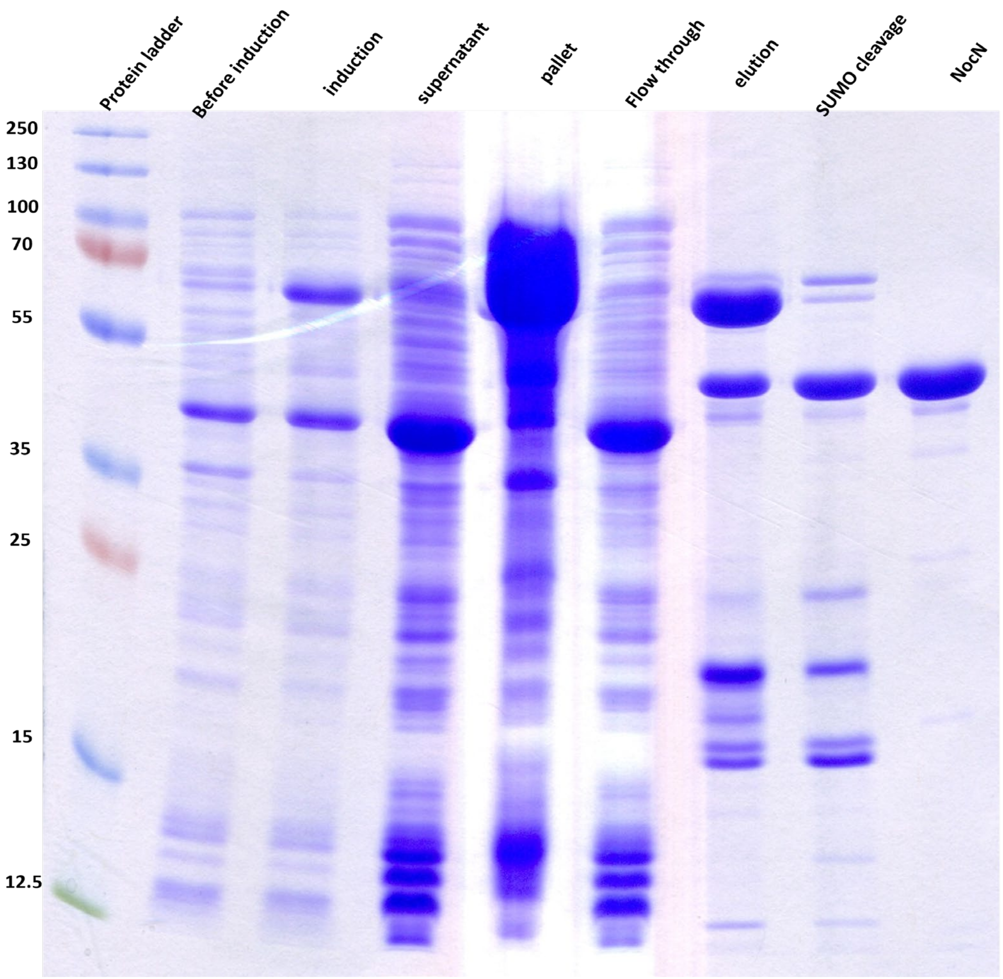


SDS-PAGE gel of NocN overexpression and purification.

### 2. Crystallization and structure determination of NocN

All manipulations were carried out in a Coy (Grass Lake, MI) anaerobic chamber at room temperature. For AzaSAM crystals, the protein solutions were prepared as follows: 6 mg/mL NocN, 50 mM HEPES pH 7.5, and 4 mM AzaSAM. All crystals were large brown plates that were obtained within 24 h using the hanging-drop vapor-diffusion method. Aza-SAM crystals formed in 200 mM $Ca(OAc)_2$, 20% w/v PEG 3350 as the precipitating reagent by mixing 2 μl of protein with 2 μl precipitating solution in the hanging drop and equilibrating against a precipitating solution (0.5 mL, 200 mM $Ca(OAc)_2$ and 20% w/v PEG3350). Crystals were briefly dipped in cryoprotectant (100 mM $Ca(OAc)_2$, 35% w/v PEG 3350), mounted on nylon loops, and flash-cooled in liquid nitrogen.

For SAM crystals, the protein solutions were prepared as follows: 6 mg/mL NocN, 50 mM HEPES pH 7.5, and 3 mM SAM. The crystals formed in 100 mM Hepes pH 7.5, 100 mM $Li_2SO_4$, 100 mM $MgCl_2$, 20% w/v PEG 3350, and 10% w/v 1,6-hexanediol as the precipitating reagent by mixing 2 μl of protein with 2 μl precipitating solution in the hanging drop and equilibrating against precipitating reagent (0.5 mL, 100 mM Hepes pH 7.5, 100 mM $Li_2SO_4$, 100 mM $MgCl_2$, 20% w/v PEG 3350, and 10% w/v 1,6-hexanediol). Crystals were briefly dipped in cryoprotectant (perfluoropolyether), mounted on nylon loops, and flash-cooled in liquid nitrogen.

For SRC crystal, the protein solutions were prepared as follows: 9 mg/mL NocN, 50 mM HEPES pH 7.5, 2 mM SAH, 3.3 mM SRC. Notably, the SRC product stock solution was 10 mM in isopropanol-water (30%-70% v/v), due to the low aqueous solubility. Thus, in the protein solution, there is 10%v/v isopropanol. The crystals formed in 100 mM Tris pH 8.0, 200 mM $CaCl_2$, and 20% w/v PEG 6000 as the precipitating reagent by mixing 2 μl of protein with 2 μl precipitating solution in the hanging drop and equilibrating against precipitating reagent (0.5 mL, 100 mM Tris pH 8.0, 200 mM $CaCl_2$, and 20% w/v PEG 6000). After equilibrating for 72 hours, the mixture was seeded to assist the formation of well-diffracting crystals. Crystals were briefly dipped in cryoprotectant (50 mM Tris pH 8.0, 100 mM $CaCl_2$, and 35% w/v PEG 6000), mounted on nylon loops, and flash-cooled in liquid nitrogen.

X-ray diffraction datasets were collected at the General Medical Sciences and Cancer Institutes Collaborative Access Team (GM/CA-CAT) and the Life Sciences Collaborative Access Team (LS-CAT) at the Advanced Photon Source, Argonne National Laboratory, and at the Berkeley Center for Structural Biology (BCSB) beamlines at the Advanced Light Source at Lawrence Berkeley National Laboratory. All datasets were processed using the HKL2000 package, and structures were determined by single anomalous dispersion (SAD) phasing using autosol/HySS or by molecular replacement using the program PHASER.[3-6] Model building and refinement were performed with Coot and phenix.refine.[3,7] Data collection and refinement statistics are shown in **Table S1**. Figures were prepared using PyMOL.[8,9]

Diffraction data for SAD phasing were collected at the iron K-edge absorption peak ($\lambda$=1.7219 Å) with 360° of data measured using a 1.0° oscillation range to 2.80 Å resolution (**Table S1**). Additionally, a 1.80-Å resolution native dataset, with AzaSAM bound, was collected at $\lambda$ =1.0332 Å. Heavy atom sites were identified using HySS implemented with Phenix Autosol.[3] The initial figure-of-merit (FOM) was 0.367, and the Bayes CC was 14.5.[3] Phenix Autobuild was used to generate an initial model of 308 residues out of 420 in chain A and 384 residues out of 420 in chain B, with $R_{work}/R_{free}$ of 0.24/0.32. This model was subjected to one cycle of rigid body refinement before being used to obtain phase information for the 1.80-Å resolution native dataset by molecular replacement using Phenix Phase-MR.[3] Iterative manual model building and refinement were performed in Coot and Phenix.[3,7] $R_{free}$ flags were determined by Phenix using the default settings (10% or up to 2000 reflections) A high-resolution (HR, 1.40 Å) AzaSAM structure was collected at λ=0.9999 Å. Using Phase-MR, the 1.40-Å model was fit into the HR map data as detailed above. Geometric restraints for AzaSAM were obtained from the Grade Web Server (Global Phasing). The final HR AzaSAM bound structure consists of residues 19-418 (of 420 residues), one [4Fe-4S] cluster, and two molecules of AzaSAM in chain A. The final structure also contains 517 water molecules. Analysis of the Ramachandran statistics showed that 98.5% of residues are in favored regions, with the remaining 1.0% and 0% in allowed and disallowed regions, respectively. Data collection and refinement statistics are shown in **Table S1**.

Structures containing the other SAM analogs were processed the same as above, using the same $R_{free}$ test set, and phasing with Phase-MR using the model from the 1.40-Å dataset. The structures were then manually adjusted with Coot, fitting in substrates if appropriate and building in loops. The following structures were obtained: SAM bound and SRC bound. The SAM bound structure consists of residues 20-238 and 245-417 (of 420 residues), one [4Fe-4S] cluster, one SAM molecule, one SAH molecule, three molecules of 1,6-hexanediol, and one molecule of imidazole in chain A; residues 20-235 and 237-417, one [4Fe-4S] cluster, one SAM molecule, one SAH molecule, and one molecule of 1,6-hexanediol in chain B. The structure also consists of 692 water molecules. Analysis of the Ramachandran statistics showed that 98.7% of residues are in favored regions, with the remaining 1.3% and 0% in allowed and disallowed regions, respectively. The SRC bound structure consists of residues 20-416 (of 420 residues), one [4Fe-4S] cluster, two SAH molecules, two molecules of SRC, one calcium ion, and one nickle ion in chain A; residues 20-233 and 243-414, one [4Fe-4S] cluster, two SAH molecules, two molecules of SRC, and one calcium ion in chain B. The structure also consists of 765 waters. Analysis of the Ramachandran statistics showed that 98.3% of residues are in favored regions, with the remaining 1.7% and 0% in allowed and disallowed regions, respectively. Geometric restraints for SRC were obtained by eLBOW using the eLBOW AM1 QM method. Restraints for [4Fe-4S] clusters were based on *M. thermoacetica* carbon monoxide dehydrogenase/acetyl-CoA synthase (PDB ID: 3I01).[10]

**Table S1**. Crystallographic data table for NocN structures obtained.

| | NocN/azaSAM(peak) | NocN+AzaSAM (HR) | NocN+SAM | NocN+SAH+SRC |
|---|---|---|---|---|
| **Data collection** | | | | |
| Wavelength | 1.72186 | 0.99999 | 0.97856 | 0.97856 |
| Resolution range | 50 – 2.80 (2.85 – 2.80) | 50 - 1.40 (1.42 - 1.40) | 50 - 1.84 (1.87 - 1.84) | 50 - 1.78 (1.81 - 1.78) |
| Space group | $P\,2\,2_1\,2_1$ | $P\,1\,2_1\,1$ | $P\,2\,2_1\,2_1$ | $P\,2\,2_1\,2_1$ |
| Cell dimentions | | | | |
| *a, b, c* (Å) | 59.9, 86.3, 144.8 | 46.6, 70.5, 54.7 | 60.9, 86.2, 145.0 | 59.9, 88.0, 146.9 |
| α, β, γ (°) | 90, 90, 90 | 90, 98, 90 | 90, 90, 90 | 90, 90, 90 |
| Total reflections | | 119228 (5503) | 107114 (5984) | 117445 (6977) |
| Unique reflections | 19281 | 66618 (5275) | 66951 (5909) | 74656 (6898) |
| $R_{sym}$ or $R_{merge}$ * | 0.07 (0.20) | 0.036 (0.143) | 0.100 (0.878) | 0.067 (0.690) |
| $R_{pim}$ | 0.02 (0.06) | 0.020 (0.114) | 0.037 (0.322) | 0.025 (0.256) |
| I / σ * | 38.7 (14.0) | 28.6 (5.9) | 21.1 (2.7) | 28.6 (2.9) |
| $CC_{1/2}$ * | 0.99 | 0.996 (0.952) | 0.836 (0.867) | 0.999 (0.857) |
| Completeness (%) * | 100.0 (100.0) | 96.7 (73.1) | 100.0 (100.0) | 100.0 (100.0) |
| Redundancy * | 13.8 | 3.9 | 8.2 | 8.2 |
| **Refinement** | | | | |
| Resolution (Å) | | 1.40 | 1.84 | 1.78 |
| No. reflections | | 66605 | 66952 | 74544 |
| $R_{work}$ / $R_{free}$ | | 0.13 / 0.16 | 0.17 / 0.20 | 0.16 / 0.19 |
| No. atoms | | 3828 | 7047 | 7375 |
| Macromolecules | | 3222 | 6322 | 6314 |
| AzaSAM | | 81 | N/A | N/A |
| Fe/S cluster | | 8 | 16 | 16 |
| 1,6-hexanediol | | N/A | 32 | N/A |
| SAM | | N/A | 108 | N/A |
| SAH | | N/A | N/A | 130 |
| imidazole | | N/A | 5 | N/A |
| calcium | | N/A | N/A | 2 |
| Nickel | | N/A | N/A | 1 |
| SRclosed | | N/A | N/A | 276 |
| water | | 517 | 672 | 765 |
| *B*-factors (Å$^2$) | | | | |
| Macromolecules | | 16.77 | 22.83 | 25.07 |
| AzaSAM | | 11.41 | N/A | N/A |
| Fe/S cluster | | 10.11 | 15.63 | 17.40 |
| 1,6-hexanediol | | N/A | 26.63 | N/A |
| SAM | | N/A | 18.31 | N/A |
| SAH | | N/A | N/A | 18.64 |
| imidazole | | N/A | 56.94 | N/A |
| calcium | | N/A | N/A | 36.38 |
| Nickel | | N/A | N/A | 13.97 |
| SRclosed | | N/A | N/A | 48.96 |
| water | | 30.15 | 31.90 | 33.08 |
| RMS deviations | | | | |
| Bond lengths (Å) | | 0.011 | 0.008 | 0.007 |
| Bond angles (°) | | 1.36 | 0.86 | 1.08 |
| Clashs score | | 0.60 | 1.66 | 2.26 |
| Rotamer outliers (%) | | 0.29 | 0.31 | 0.00 |
| Ramachandran | | | | |
| Most favored (%) | | 98.5 | 98.7 | 98.3 |
| Allowed (%) | | 1.5 | 1.3 | 1.7 |
| Outliers (%) | | 0 | 0 | 0 |
| Number of TLS groups | | 11 | 12 | 7 |
| PDB accession code | | 9P2U | 9P3B | 9P3C |

*represents the (highest resolution shell) **all datasets result from a single protein crystal

### 3. Experimental procedures for EPR

Stocks of NosN, SAM, substrate peptides, and sodium dithionite powder were brought into an anaerobic chamber together with a Dewar containing liquid nitrogen for freezing the samples. A stock solution of sodium dithionite (1 M in 50 mM Hepes pH 7.5) was freshly prepared in an anaerobic chamber. The reaction mixtures containing Hepes pH 7.5 (100 mM), glycerol (10%), NosN (550 uM), SAM (3 mM), and substrate peptides (1.5 mM) were incubated at room temperature for 5 minutes, and then the reaction was initiated by the addition of sodium dithionite solution to the final concentration of 4 mM. The reaction mixture was incubated at room temperature for the time indicated and then frozen in liquid nitrogen. EPR data collection and simulation were performed according to the reported procedure.[11]

### 4. DFT calculations for substrate radical

All calculations were performed in Orca 5.0.1[12,13] utilizing TPSSh functional[14,15] and def2-TZVP basis set[16] on all atoms with the default general auxiliary basis set def2/J used within the Resolution of Identity (RI) approximation.[17] All calculations utilized the conductor-like polarizable continuum model (CPCM, dielectric constant ϵ=10) to account for the protein dielectric environment. Starting geometries were derived from crystallographic data.

Table 2. Hyperfine coupling constants (in MHz) for Compound **1**, **5**, and **8**

| Nucleus | Experiment** | | | | DFT-calculated*** | | | |
|---|---|---|---|---|---|---|---|---|
| | $A_x$ | $A_y$ | $A_z$ | $A_{iso}$* | $A_x$ | $A_y$ | $A_z$ | $A_{iso}$* |
| | | | | **Compound 1** | | | | |
| $^1$H (C4) | 74.5 | 78.5 | 80.9 | 78.0 | 79.1 | 80.6 | 85.2 | 81.6 |
| $^1$H (C5) | 13.4 | 23.0 | 38.7 | 25.0 | -11.1 | -28.0 | -44.8 | -28.0 |
| $^1$H (C6) | 5.6 | 9.7 | 14.2 | 9.8 | 4.8 | 7.1 | 14.0 | 8.6 |
| $^1$H (C7) | 20.4 | 40.9 | 44.5 | 35.3 | -14.8 | -33.6 | -54.0 | -34.1 |
| | | | | **Compound 5** | | | | |
| $^1$H (C4) | 78.1 | 83.0 | 85.0 | 82.0 | - | - | - | - |
| $^1$H (C5) | 13.7 | 23.2 | 35.3 | 24.1 | - | - | - | - |
| $^1$H (C6) | 4.8 | 9.6 | 14.8 | 9.7 | - | - | - | - |
| $^1$H (C7) | 23.6 | 41.3 | 46.3 | 37.1 | - | - | - | - |
| | | | | **Compound 8** | | | | |
| $^1$H (C4) | 80.3 | 91.7 | 94.8 | 89.0 | 95.3 | 96.9 | 101.3 | 97.8 |
| $^1$H (C5) | 18.8 | 19.8 | 39.7 | 26.1 | -11.1 | -28.2 | -45.1 | 28.2 |
| $^1$H (C6) | 2.0 | 3.6 | 9.3 | 5.0 | 4.8 | 7.2 | 14.1 | 8.7 |
| $^1$H (C7) | 20.4 | 42.5 | 46.1 | 36.3 | -15.0 | -34.0 | -54.5 | 34.5 |
| $^{13}$C(SAM) | 59.0 | 42.0 | 69.0 | 56.7 | 65.5 | 65.6 | 76.7 | 69.2 |

* $A_{iso}=(A_x+A_y+A_z)/3$; ** Absolute sign of HF coupling interactions is undertermined; *** DFT calculations of compound **1** and compound **5** do not include the polypeptide chains, thus the numbers obtained for these substrates are identical.

### 5. DFT Calculations for SAM Epimerization

All epimerization DFT calculations were performed using Orca (ver. 5.0.4).[12,13] Optimizations were done using unrestricted Kohn-Sham with the B3LYP functional and def2-TZVP basis set.[16,18] The calculations also utilized the RIJCOSX approximation, Grimme's D3 dispersion correction, and a CPCM solvent model using water.[19-22] Starting coordinates of SAM were obtained from $SAM^{II}$ in the crystal structure of NocN (PDB: 9P3B). Hydrogens were added to the structure using PyMol and optimized with Orca using the "OptH" keyword. Having prepared the SAM structure, the methylene radical structure was formed by removing a hydrogen atom from the methyl group and reoptimizing the hydrogens.

For both structures, a relaxed surface scan was performed using a dihedral angle between the γ-carbon of methionine (1), 5′-carbon of adenosyl (2), sulfur (3), and the methyl carbon (4) as shown in **Figure S13a**. The starting angle was -90° in the *S* conformation and scanned by 10° steps to -270° into the *R* conformation. Initial calculations showed as the SAM approached the planar sulfonium transition state, the terminal ends of the molecule swung together forming an intramolecular interaction between the methionine carboxylate and adenine ring. Though this interaction would be possible in solution, **Figure 4d** reveals multiple hydrogen bond interactions in the protein active site binding these terminal moieties in place. Therefore, to better mimic the overall conformation in the active site, the calculations were repeated with constrained coordinates for the methionine carboxylate oxygens and amine nitrogen, as well as adenine nitrogen atom at position 1 and the amine group. The resulting energy diagram is displayed in Figure **S13b**. Unrestricted natural orbitals were generated with Orca for the transition state of the methylene radical epimerization using the keyword "UNO" in the Orca input file. The singly occupied molecular orbital (SOMO) was visualized using Chemcraft (ver. 1.8) with a contour value of 0.05, as shown in **Figure S13c**.

### 6. Synthesis of SRC for crystallography

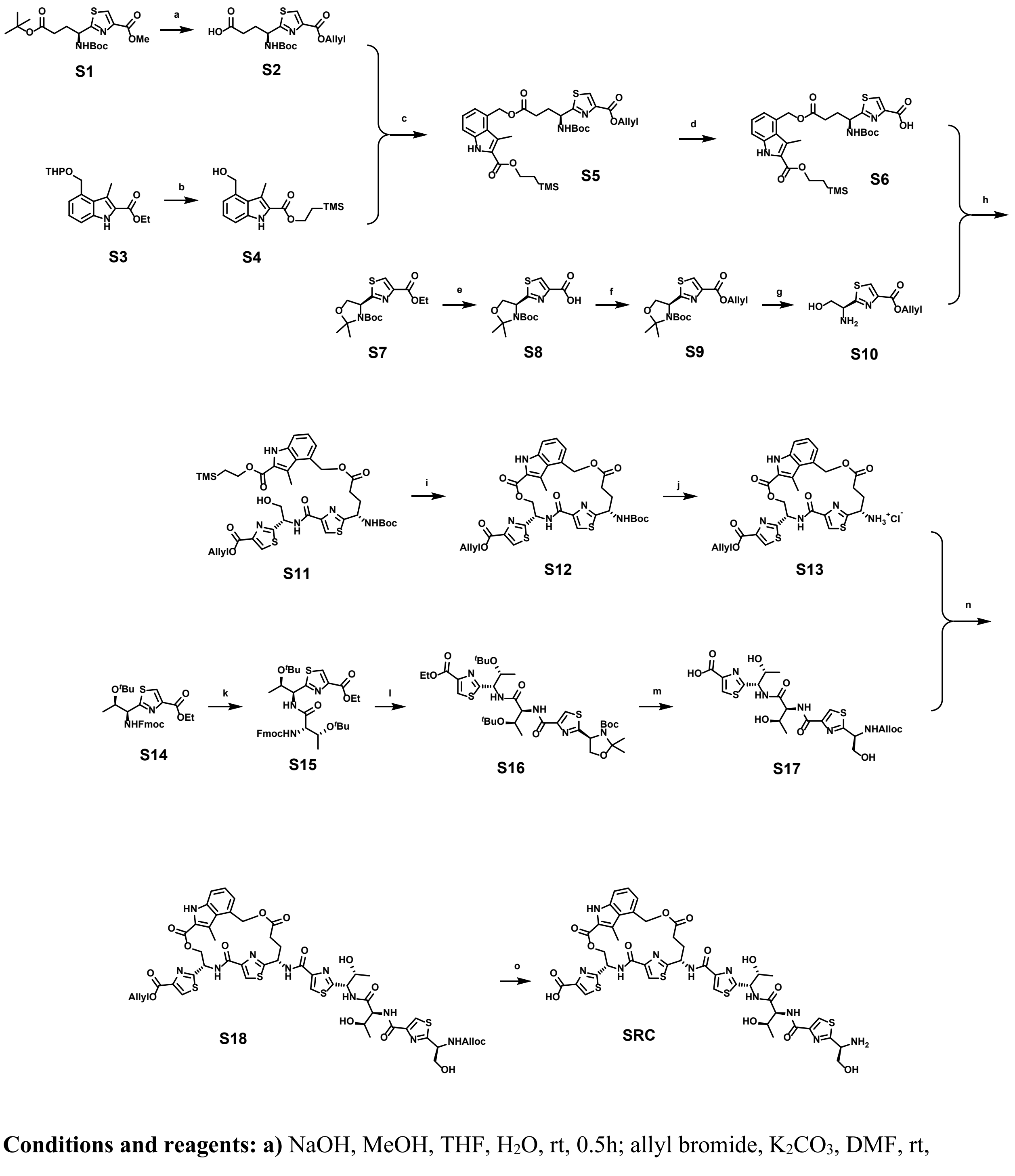


**Conditions and reagents: a)** NaOH, MeOH, THF, $H_2O$, rt, 0.5h; allyl bromide, $K_2CO_3$, DMF, rt, overnight; TFA, DCM, rt, 1h; $Boc_2O$, $NaHCO_3$, THF, $H_2O$, rt, overnight, 70% over four steps; **b)** NaOH, MeOH, THF, $H_2O$, 75 °C, 3h; 2-(trimethylsilyl)ethanol, EDC, DMAP, DCM, rt, overnight; *p*-TsOH, DCM, MeOH, rt, 2h, 70% over three steps; **c)** EDC, DMAP, DCM, rt, overnight, 75%; **d)** $Pd(PPh_3)_4$, 1,3-dimethylbarbituric acid , THF, $H_2O$, rt, 1h; **e)** NaOH, MeOH, THF, $H_2O$, rt, 0.5h, 95%; **f)** allyl bromide,

$K_2CO_3$, DMF, rt, overnight, 85%; **g)** TFA, DCM, $H_2O$, rt, 1h; **h)** HATU, TEA, DMF, 0 ºC, 1h, 82%; **i)** TBAF, THF, rt, 1h; $PPh_3$, DEAD, THF, rt, 80% over two steps; **j)** 4M HCl in dioxane, rt, 2h; **k)** TBAF, THF, rt; Fmoc-Thr(O$^t$Bu)-OH, HATU, TEA, DMF, 0 ºC, 75% over two steps; **l)** TBAF, THF, rt; **S8**, HATU, TEA, DMF, 0 ºC, 70% over two steps; **m)** NaOH, MeOH, THF, $H_2O$, rt, 2h; TFA, $H_2O$, rt, 1h; AllocCl, $NaHCO_3$, $H_2O$, acetonitrile, rt, 1h; **n)** HATU, TEA, DMF, rt, 1h, 55%; **o)** $Pd(PPh_3)_4$, THF, $PhSiH_3$, rt, 1h, 20%.

**Step a:**

Compound **S1** was synthesized according to the reported procedure.[23]

To the stirred solution of **S1** (4.0 g, 10.0 mmol, 1.0 equiv) in THF (50 mL)-MeOH (50 mL)-water (50 mL) was added sodium hydroxide (1.2 g, 30.0 mmol, 3.0 equiv). The reaction mixture was stirred at room temperature for 0.5 h. After completion, the reaction mixture was diluted with water (100 mL) and washed with diethyl ether (100 mL). The aqueous layer was acidified by hydrochloric acid (1 M) to pH 3.0 and extracted with ethyl acetate (100 mL). The organic layer was washed with water and brine. The organic layer was dried over anhydrous sodium sulfate and concentrated *in vacuo*, and the resulting residue was used for the next step without further purification.

To an ice-water cooled solution of the residue from the previous step in DMF (50 mL) was added $K_2CO_3$ (2.8 g, 20.0 mmol, 2.0 equiv) and allyl bromide (1.8 g, 15.0 mmol, 1.5 equiv). The resulting reaction was warmed up to room temperature and stirred overnight. After completion, the reaction mixture was diluted with ethyl acetate (200 mL) and washed thoroughly with hydrochloric acid (0.1 M), saturated aqueous sodium bicarbonate, and brine. The organic layer was dried over anhydrous sodium sulfate and concentrated *in vacuo*, and the resulting residue was used for the next step without further purification.

To an ice-water cooled solution of the residue from the previous step in DCM (50 mL) was added TFA (30 mL). The resulting reaction was warmed up to room temperature and stirred for 1 h. After completion, the reaction mixture was concentrated *in vacuo*, and the resulting residue was used for the next step without further purification.

To an ice-water cooled solution of the residue from the previous step in THF (50 mL) was added saturated aqueous sodium bicarbonate (50 mL) and $Boc_2O$ (3.3 g, 15.0 mmol, 1.5 equiv). After completion, the reaction mixture was diluted with water (100 mL) and washed with diethyl ether (100 mL). The aqueous layer was acidified by hydrochloric acid (1 M) to pH 3.0 and extracted with ethyl acetate (150 mL). The organic layer was washed with water and brine. The organic layer was dried over anhydrous sodium sulfate and concentrated *in vacuo*, giving 2.6 g of compound **S2** as a yellowish gum in a yield of 70%. $^1$H NMR (500 MHz, DMSO) δ 12.18 (s, 1H), 8.47 (s, 1H), 7.84 (d, *J* = 8.0 Hz, 1H), 6.12 – 5.95 (m, 1H), 5.40 (dd, *J* = 17.2, 1.6 Hz, 1H), 5.27 (dd, *J* = 10.5, 1.5 Hz, 1H), 4.86 – 4.75 (m, 3H), 2.43 – 2.28 (m, 2H), 2.27 – 2.12 (m, 1H), 1.99 – 1.84 (m, 1H), 1.41 (s, 9H); $^{13}$C NMR (126 MHz, DMSO) δ 176.10, 174.22, 160.79, 155.86, 145.95, 132.97, 129.67, 118.66, 79.14, 65.51, 52.63, 30.48, 29.66, 28.62; **HRMS**: calculated for $C_{16}H_{21}N_2O_6S^-$ [M-H$^+$]: 369.11258; found: 369.11262.

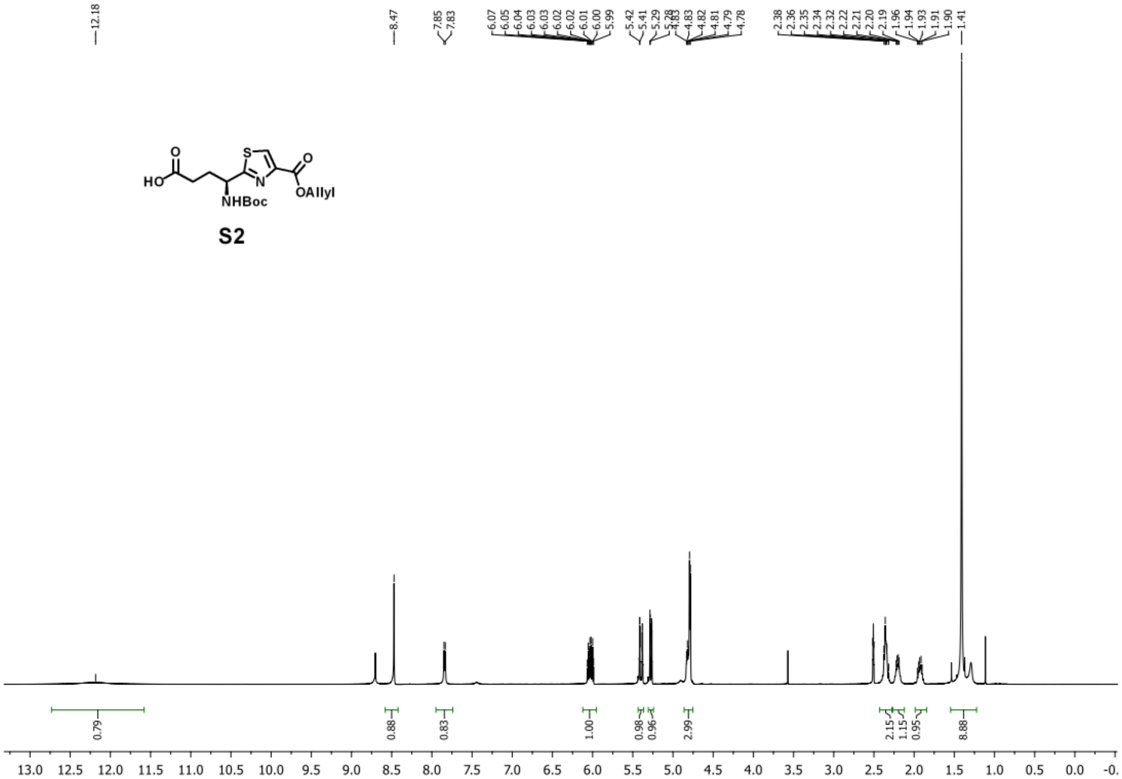

S2

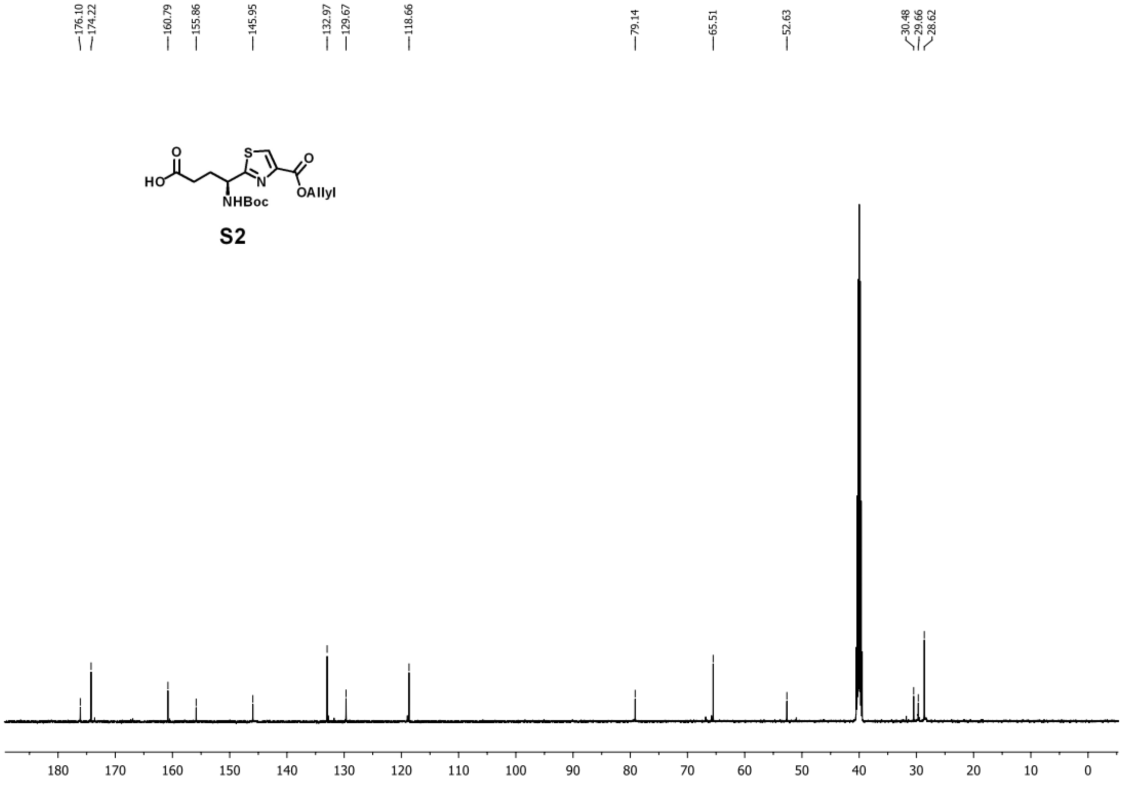

S2

**Step b:**

Compound **S3** was synthesized according to the reported procedure.[2]

To the stirred solution of **S3** (2.5 g, 7.9 mmol, 1.0 equiv) in THF (50 mL)-MeOH (50 mL)-water (50 mL) was added sodium hydroxide (1.3 g, 31.6 mmol, 4.0 equiv). The reaction mixture was refluxed at 75 °C for 3 h. After completion, the reaction mixture was diluted with water (100 mL) and washed with diethyl ether (100 mL). The aqueous layer was acidified by hydrochloric acid (1 M) to pH 3.0 and extracted with ethyl acetate (100 mL). The organic layer was washed with water and brine. The organic layer was dried over anhydrous sodium sulfate and concentrated *in vacuo*, and the resulting residue was used for the next step without further purification.

To an ice-water cooled solution of the residue from the previous step in DCM (100 mL) was added 2-(trimethylsilyl)ethanol (2.3 g, 20.0 mmol, 2.5 equiv), EDC (3.0 g, 15.8 mmol, 2.0 equiv), and DMAP (200 mg, 1.6 mmol, 0.2 equiv). The resulting reaction was warmed up to room temperature and stirred overnight. After completion, the reaction mixture was diluted with DCM (100 mL) and washed thoroughly with hydrochloric acid (0.1 M), saturated aqueous sodium bicarbonate, and brine. The organic layer was dried over anhydrous sodium sulfate and concentrated *in vacuo*, and the resulting residue was used for the next step without further purification.

To a stirred solution of the residue from the previous step in DCM (50 mL) and methanol (50 mL) was added p-TsOH (680 mg, 4.0 mmol, 0.5 equiv). The resulting reaction was stirred at room temperature for 2 h. After completion, the reaction mixture was diluted with DCM (100 mL) and washed thoroughly with saturated aqueous sodium bicarbonate and brine. The organic layer was dried over anhydrous sodium sulfate and concentrated *in vacuo*, and the resulting residue was purified by silica gel flash chromatography (hexanes : ethyl acetate = 20:1-5:1), giving 1.7 g of compound **S4** as a white solid in a yield of 70% over three steps. $^{1}$H NMR (500 MHz, $CDCl_3$) δ 8.95 (s, 1H), 7.32 (dd, *J* = 8.2, 0.7 Hz, 1H), 7.29 – 7.24 (m, 1H), 7.11 (d, *J* = 6.9 Hz, 1H), 5.08 (s, 2H), 4.54 – 4.40 (m, 2H), 2.86 (s, 3H), 2.07 (s, 1H), 1.27 – 1.13 (m, 2H), 0.12 (s, 9H); $^{13}$C NMR (126 MHz, $CDCl_3$) δ 162.93, 136.49, 135.47, 125.95, 125.21, 123.82, 120.28, 120.23, 112.01, 63.70, 63.16, 17.70, 11.91, -1.44; **HRMS**: calculated for $C_{16}H_{24}NO_3Si^+$ [$M+H^+$]: 306.15200; found: 306.15240.

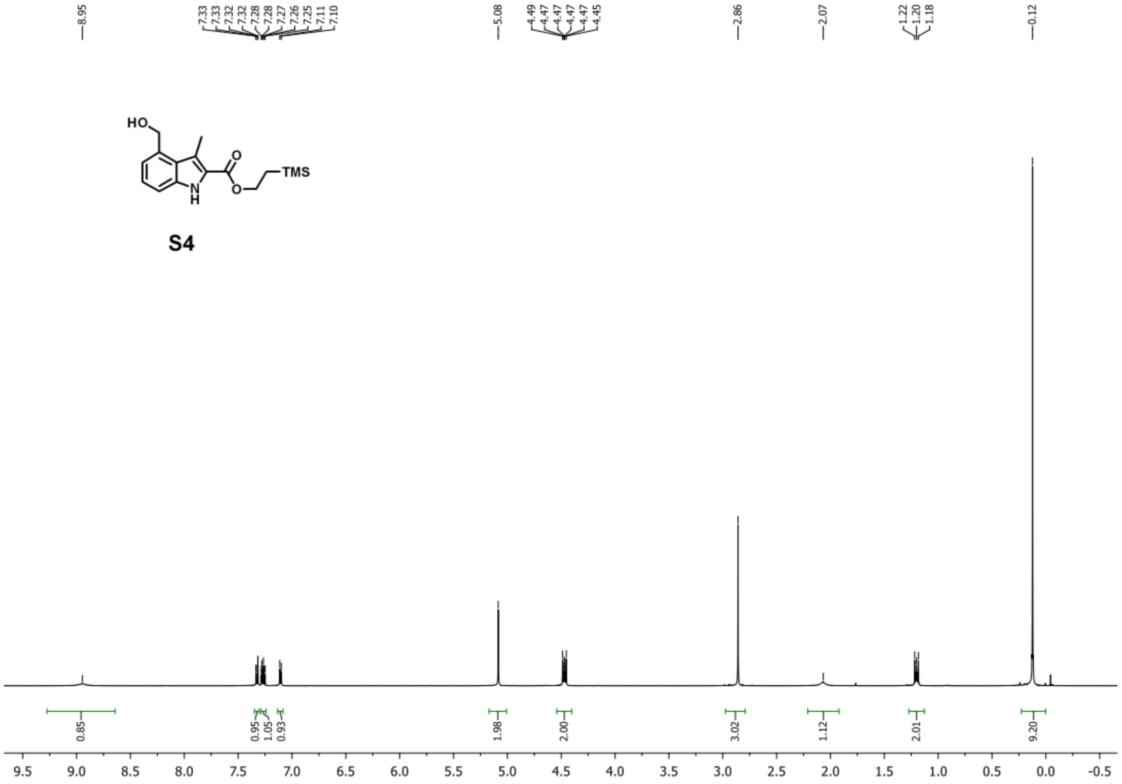

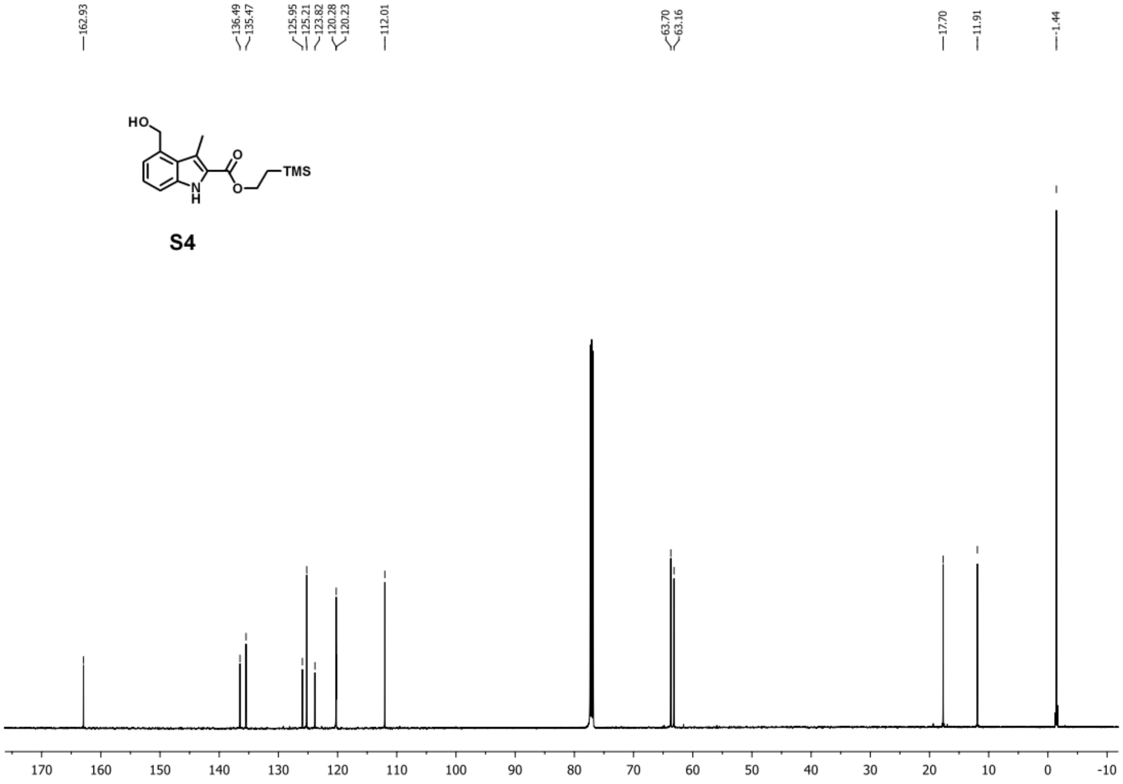


**Step c:**

To an ice-water cooled solution of S2 (1.5 g, 4.0 mmol, 1.0 equiv) and S4 (1.2 g, 4.0 mmol, 1.0 equiv) in DCM (100 mL) was added EDC (1.2 g, 6.0 mmol, 1.5 equiv) and DMAP (98 mg, 0.8 mmol, 0.2 equiv). The resulting reaction was warmed up to room temperature and stirred overnight. After completion, the reaction mixture was diluted with DCM (100 mL) and washed thoroughly with hydrochloric acid (0.1 M), saturated aqueous sodium bicarbonate, and brine. The organic layer was dried over anhydrous sodium sulfate and concentrated *in vacuo*, and the resulting residue was purified by silica gel flash chromatography (hexanes : ethyl acetate = 10:1-4:1), giving 2.0 g of compound **S5** as white foam in a yield of 75%. $^{1}$H NMR (500 MHz, $CDCl_3$) δ 9.24 – 8.95 (m, 1H), 8.11 (d, $J$ = 2.9 Hz, 1H), 7.41 – 7.34 (m, 1H), 7.29 – 7.22 (m, 1H), 7.13 – 7.07 (m, 1H), 6.11 – 5.94 (m, 1H), 5.64 – 5.55 (m, 1H), 5.54 – 5.47 (m, 2H), 5.45 – 5.36 (m,

1H), 5.34 – 5.26 (m, 1H), 5.09 (s, 1H), 4.89 – 4.79 (m, 2H), 4.53 – 4.40 (m, 2H), 2.77 (s, 3H), 2.60 – 2.51 (m, 2H), 2.50 – 2.39 (m, 1H), 2.29 – 2.17 (m, 1H), 1.44 (s, 9H), 1.22 – 1.15 (m, 2H), 0.13 – 0.08 (m, 9H); $^{13}$C NMR (126 MHz, $CDCl_3$) δ 173.26, 172.77, 162.81, 160.86, 155.22, 146.92, 136.44, 131.83, 129.74, 127.63, 126.39, 125.03, 124.15, 122.29, 119.67, 118.96, 112.80, 80.32, 66.00, 65.23, 63.21, 52.52, 30.72, 30.41, 28.31, 17.70, 11.82, -1.45; **HRMS**: calculated for $C_{32}H_{44}N_3O_8SSi^+$ [M+H$^+$]: 658.26129; found: 658.26156.

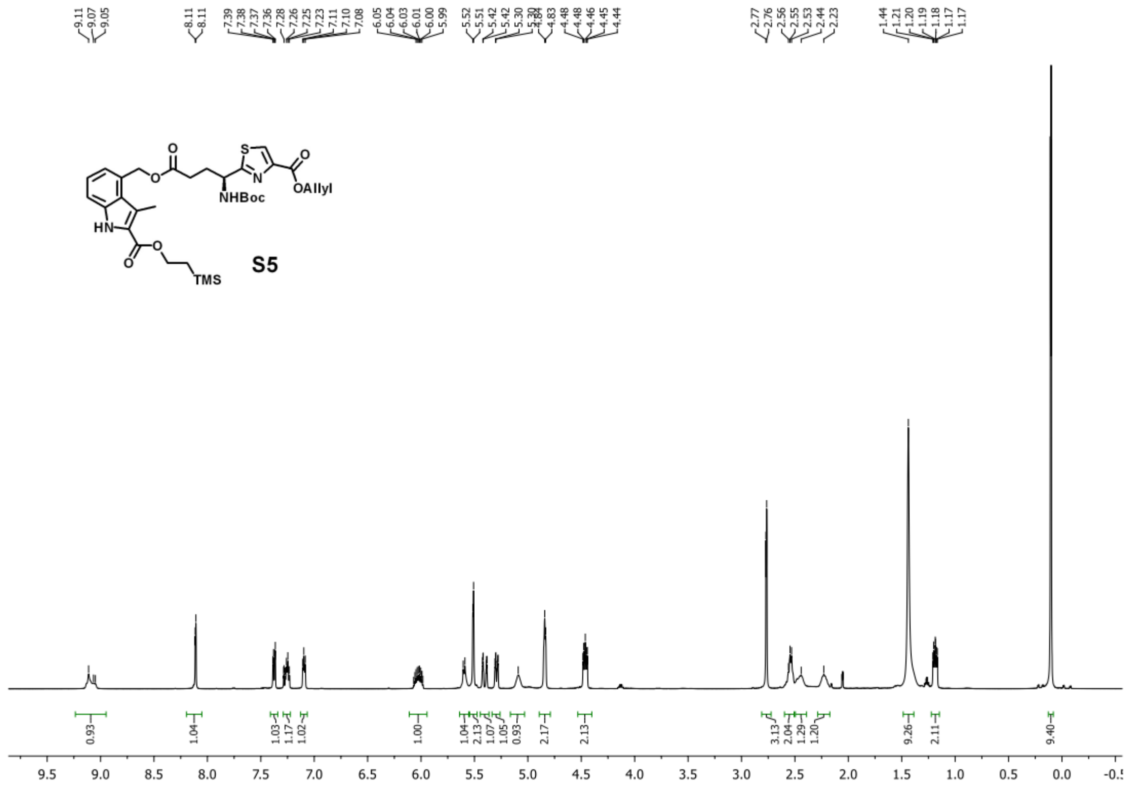

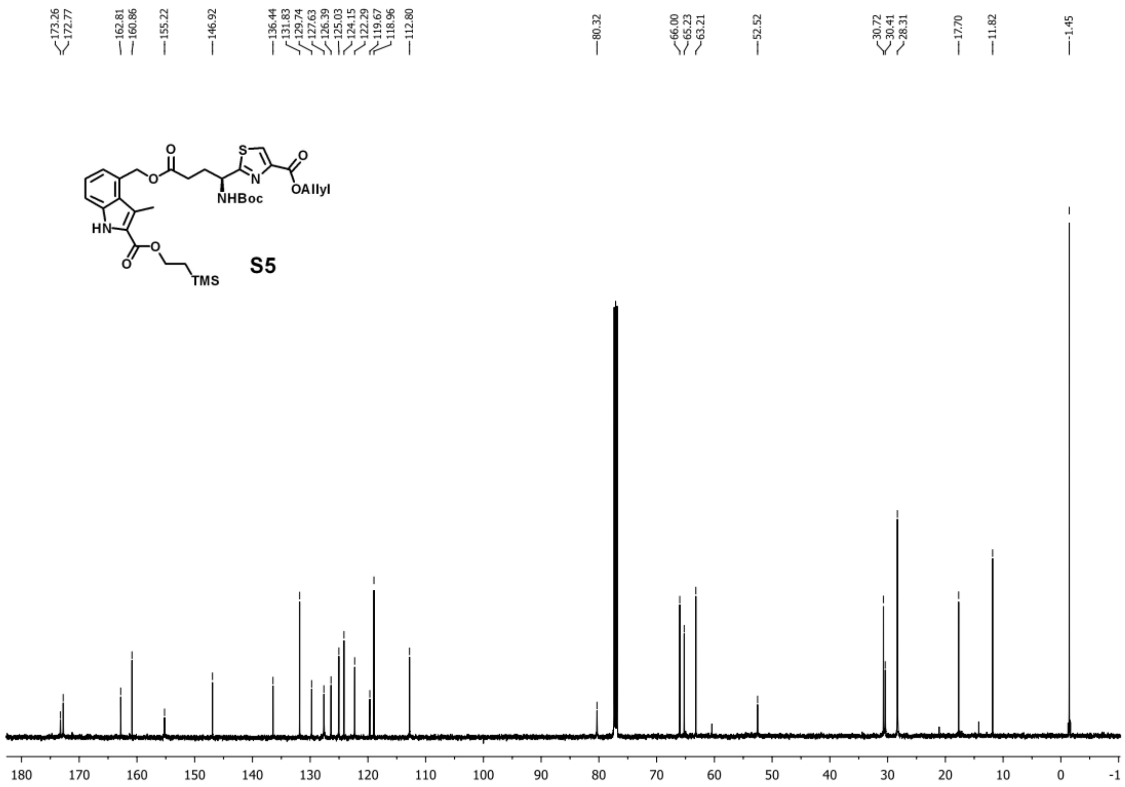


**Step d:**

To a stirred solution of **S5** (500 mg, 0.76 mmol, 1.0 equiv) in THF (20 mL) and $H_2O$ (2 mL) was added $Pd(PPh_3)_4$ (87 mg, 0.076 mmol, 0.1 equiv) and 1,3-dimethylbarbituric acid (118 mg, 0.76 mmol, 1.0 equiv). The resulting reaction was stirred at room temperature for 1 h. After completion, the reaction mixture was diluted with ethyl acetate (100 mL) and washed thoroughly with water and brine. The organic layer was dried over anhydrous sodium sulfate and concentrated *in vacuo*, and the resulting compound **S6** was used for the next step without further purification.

**Step e:**

Compound **S7** was synthesized according to the reported procedure.[2,24]

To the stirred solution of **S7** (2.8 g, 7.9 mmol, 1.0 equiv) in THF (50 mL)-MeOH (50 mL)-water (50 mL) was added sodium hydroxide (1.3 g, 31.6 mmol, 4.0 equiv). The reaction mixture was stirred at room temperature for 0.5 h. After completion, the reaction mixture was diluted with water (100 mL) and washed with diethyl ether (100 mL). The aqueous layer was acidified by hydrochloric acid (1 M) to pH 3.0 and extracted with ethyl acetate (100 mL). The organic layer was washed with water and brine. The organic layer was dried over anhydrous sodium sulfate and concentrated *in vacuo*, giving 2.5 g of compound **S8** as a white solid in a yield of 95%. $^{1}H$ NMR (500 MHz, DMSO at 345K) δ 12.73 (s, 1H), 8.33 (s, 1H), 5.37 – 5.08 (m, 1H), 4.33 (dd, *J* = 9.2, 6.4 Hz, 1H), 4.08 (dd, *J* = 9.2, 1.3 Hz, 1H), 1.67 (s, 3H), 1.54 (s, 3H), 1.36 (s, 9H); $^{13}C$ NMR (126 MHz, DMSO at 345K) δ 173.81, 162.34, 147.54, 128.57, 94.74, 80.69, 68.88, 59.30, 28.38, 27.03, 23.88; **HRMS**: calculated for $C_{14}H_{19}N_2O_5S^-$ [M-H$^+$]: 327.10202; found: 327.10301.

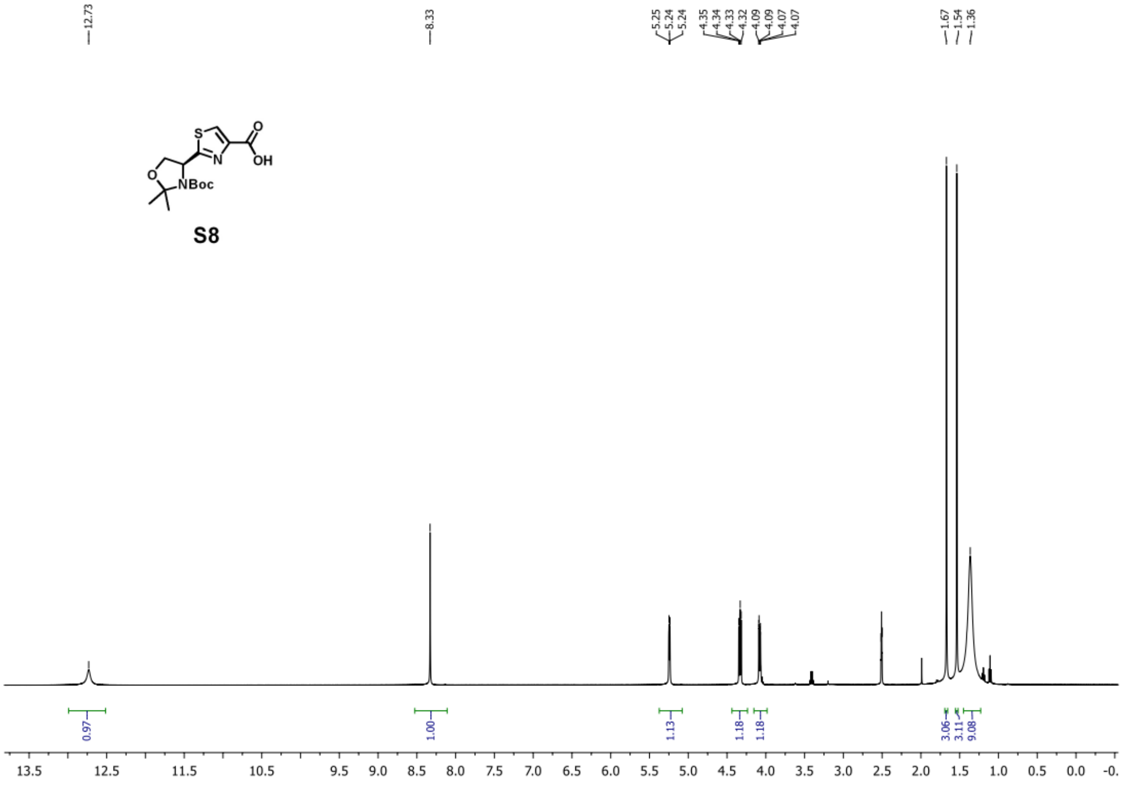

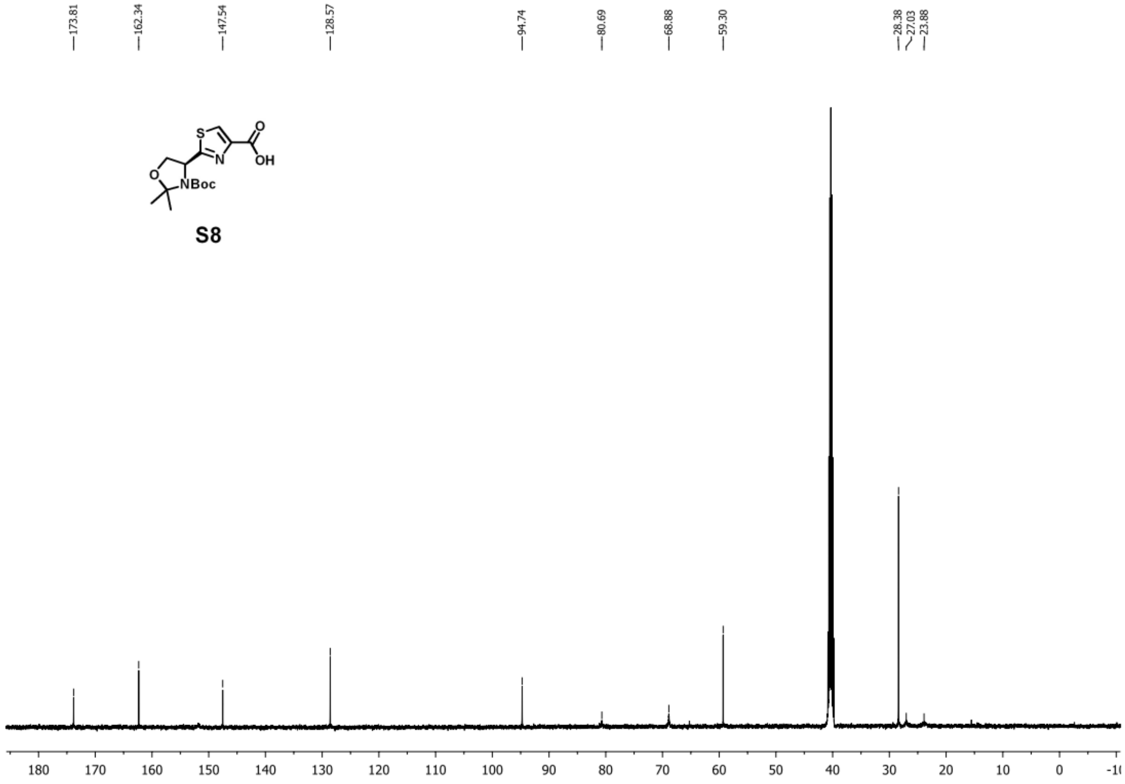


**Step f:**

To an ice-water cooled solution of S8 (1.0 g, 3.0 mmol, 1.0 equiv) in DMF (20 mL) was added $K_2CO_3$ (1.2 g, 9.0 mmol, 3.0 equiv) and allyl bromide (0.52 mL, 6.0 mmol, 2.0 equiv). The resulting reaction was warmed up to room temperature and stirred overnight. After completion, the reaction mixture was diluted with ethyl acetate (200 mL) and washed thoroughly with hydrochloric acid (0.1 M), saturated aqueous sodium bicarbonate, and brine. The organic layer was dried over anhydrous sodium sulfate and concentrated *in vacuo*, giving 0.94 g of compound **S9** as a white solid in a yield of 85%. $^1$H NMR (500 MHz, DMSO) δ 8.45 (s, 1H), 6.13 – 5.92 (m, 1H), 5.41 (dq, $J$ = 17.2, 1.6 Hz, 1H), 5.33 – 5.23 (m, 2H), 4.85 – 4.76 (m, 2H), 4.34 (dd, $J$ = 9.3, 6.4 Hz, 1H), 4.08 (dd, $J$ = 9.3, 1.5 Hz, 1H), 1.68 (s, 3H), 1.54 (s, 3H), 1.37 (s, 9H); $^{13}$C NMR (126 MHz, DMSO) δ 174.34, 160.78, 146.06, 132.99, 129.46, 118.53, 94.77, 80.72, 68.88,

65.47, 59.31, 28.37, 27.01, 23.80; **HRMS**: calculated for $C_{17}H_{25}N_2O_5S^+$ [M+H$^+$]: 369.14787; found: 369.14850.

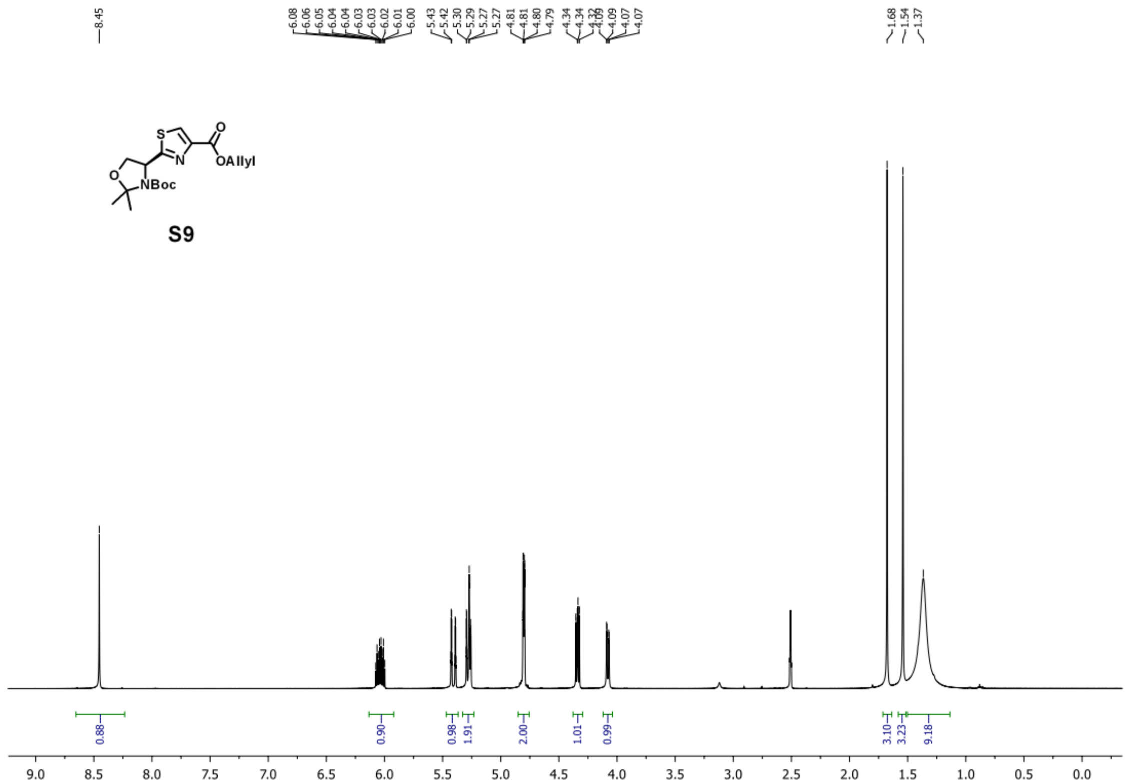

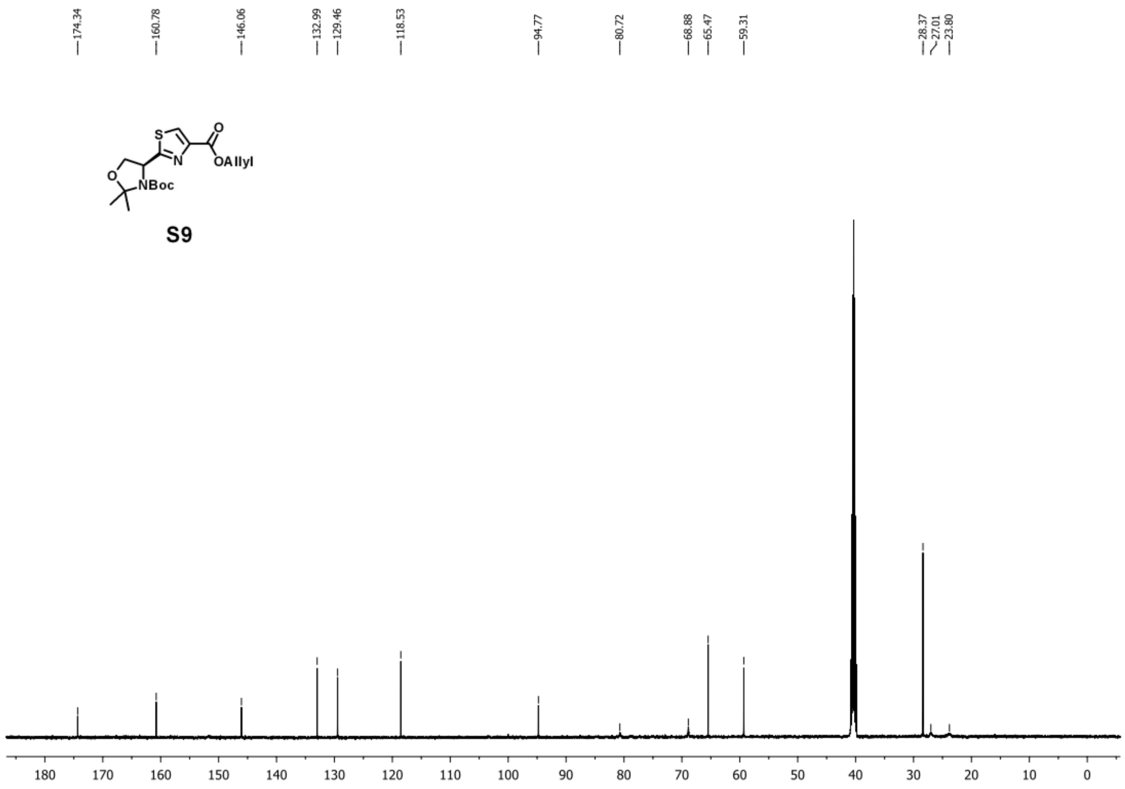


**Step g:**

To an ice-water cooled solution of **S9** (420 mg, 1.1 mmol, 1.0 equiv) in DCM (10 mL) was added TFA (10 mL) and $H_2O$ (1 mL). The resulting reaction was warmed up to room temperature and stirred for 1 h. After completion, the reaction mixture was concentrated *in vacuo*, and the resulting residue was used for the next step without further purification.

**Step h:**

To an ice-water cooled solution of **S6** (0.76 mmol, 1.0 equiv) and **S10** (1.1 mmol, 1.5 equiv) in DMF (20 mL) was added HATU (433 mg, 1.1 mmol, 1.5 equiv) and TEA (0.53 mL, 3.8 mmol, 5.0 equiv). The resulting reaction was stirred at 0 °C for 1 h. After completion, the reaction mixture was diluted with ethyl acetate (100 mL) and washed thoroughly with hydrochloric acid

(0.1 M), saturated aqueous sodium bicarbonate, and brine. The organic layer was dried over anhydrous sodium sulfate and concentrated *in vacuo*, and the resulting residue was purified by silica gel flash chromatography (hexanes : acetone = 10:1-3:1), giving 516 mg of compound **S11** as white foam in a yield of 82%. $^{1}$H NMR (500 MHz, DMSO at 345K) δ 11.58 (s, 1H), 8.72 (d, *J* = 8.0 Hz, 1H), 8.46 (s, 1H), 8.27 (s, 1H), 7.89 (d, *J* = 7.9 Hz, 1H), 7.43 (d, *J* = 8.3 Hz, 1H), 7.20 (dd, *J* = 8.2, 7.2 Hz, 1H), 7.03 (d, *J* = 7.0 Hz, 1H), 6.11 – 5.94 (m, 1H), 5.50 – 5.32 (m, 4H), 5.31 – 5.24 (m, 1H), 4.91 – 4.84 (m, 1H), 4.79 (d, *J* = 5.0 Hz, 2H), 4.45 – 4.36 (m, 2H), 4.01 – 3.89 (m, 2H), 2.89 (s, 2H), 2.74 (s, 2H), 2.68 (s, 3H), 2.61 – 2.52 (m, 2H), 2.38 – 2.27 (m, 1H), 2.07 – 1.95 (m, 1H), 1.40 (s, 9H), 1.17 – 1.10 (m, 2H), 0.07 (s, 9H); $^{13}$C NMR (126 MHz, DMSO at 345K) δ 176.10, 172.50, 172.30, 162.77, 162.49, 160.97, 160.79, 155.85, 149.31, 145.89, 137.25, 132.96, 132.00, 131.92, 129.99, 129.87, 129.28, 129.18, 125.97, 125.20, 124.75, 124.23, 121.85, 118.69, 118.66, 113.74, 79.19, 65.53, 65.08, 63.23, 62.65, 54.10, 52.64, 36.25, 31.24, 30.57, 29.75, 28.60, 17.73, 11.77, -1.02; **HRMS**: calculated for $C_{38}H_{50}N_5O_{10}S_2Si^+$ [M+H$^+$]: 828.27629; found: 828.27717.

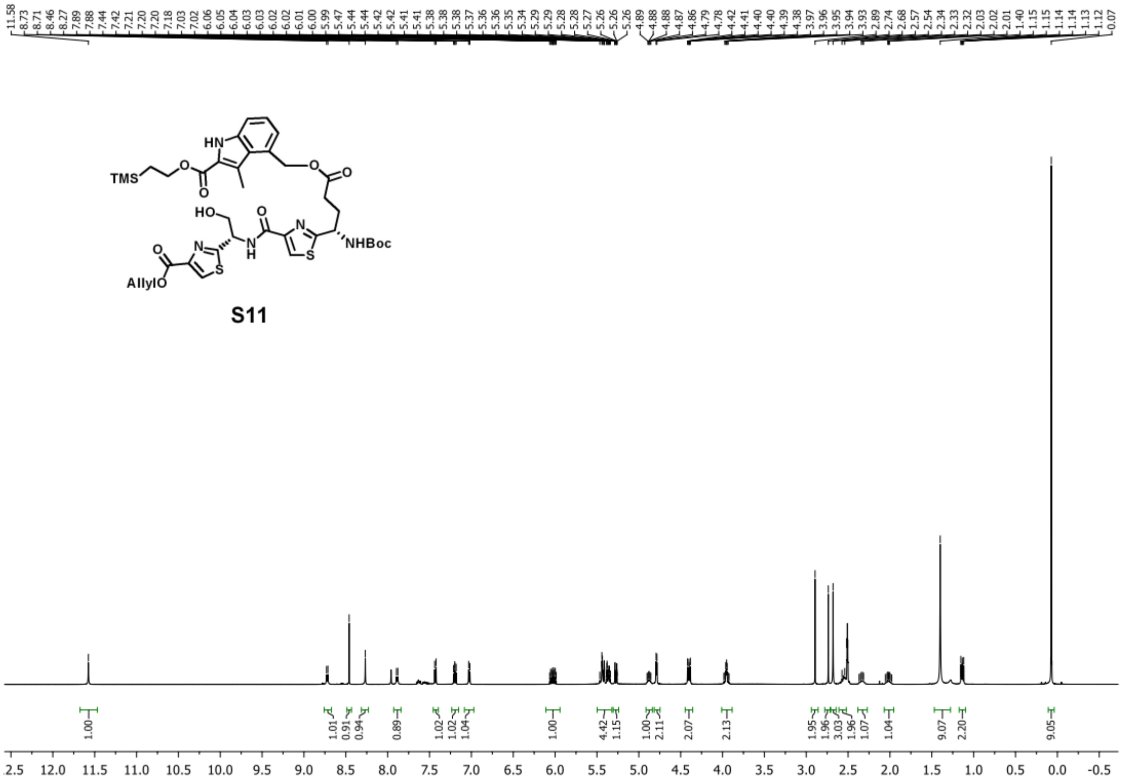
11.58 8.73 8.71 8.46 8.27 7.89 7.88 7.44 7.42 7.21 7.20 7.20 7.18 7.03 7.02 6.06 6.05 6.04 6.03 6.03 6.02 6.01 6.00 5.99 5.47 5.44 5.44 5.42 5.41 5.38 5.38 5.38 5.37 5.36 5.36 5.35 5.34 5.29 5.29 5.28 5.28 5.27 5.26 5.26 5.26 4.89 4.88 4.88 4.87 4.86 4.79 4.78 4.42 4.41 4.40 4.40 4.39 4.38 3.97 3.96 3.95 3.94 3.93 2.89 2.74 2.68 2.57 2.54 2.34 2.33 2.32 2.03 2.02 2.01 1.40 1.15 1.14 1.13 1.12 0.07
TMS
HN
HO
NHBoc
AllylO
S11
1.00 1.01 0.91 0.94 0.89 1.02 1.02 1.04 1.00 4.42 1.15 1.00 2.11 2.07 2.13 1.95 1.96 3.03 1.96 1.07 1.04 9.07 2.20 9.05
2.5 12.0 11.5 11.0 10.5 10.0 9.5 9.0 8.5 8.0 7.5 7.0 6.5 6.0 5.5 5.0 4.5 4.0 3.5 3.0 2.5 2.0 1.5 1.0 0.5 0.0 -0.5

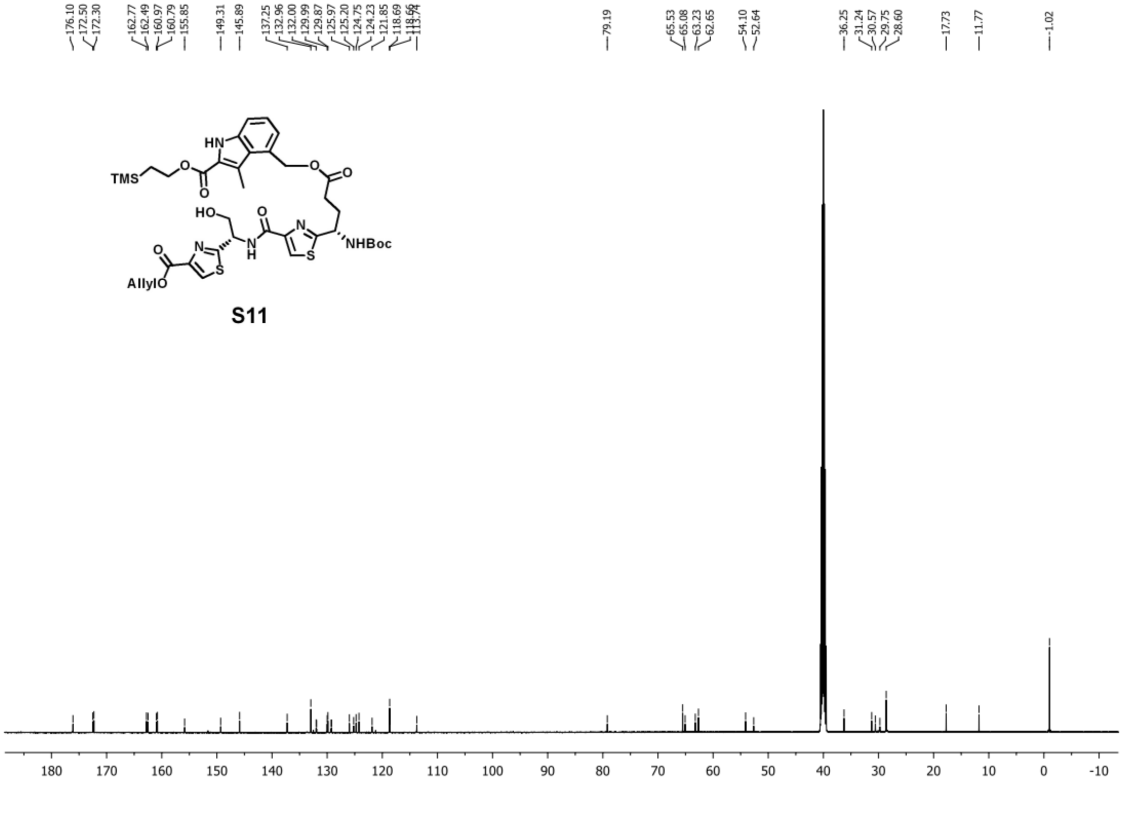
176.10 172.50 172.30 162.77 162.49 160.97 160.79 155.85 149.31 145.89 137.25 132.96 132.00 129.99 129.87 125.97 125.20 124.75 124.23 121.85 118.69 113.74 79.19 65.53 65.08 63.23 62.65 54.10 52.64 36.25 31.24 30.57 29.75 28.60 17.73 11.77 -1.02
TMS
HN
HO
NHBoc
AllylO
S11
180 170 160 150 140 130 120 110 100 90 80 70 60 50 40 30 20 10 0 -10

**Step i:**

To a stirred solution of **S11** (500 mg, 0.6 mmol, 1.0 equiv) in THF (20 mL) was added TBAF (1.2 mL, 1.2 mmol, 2.0 equiv, 1M solution in THF). The resulting reaction was stirred for 1 h. After completion, the reaction mixture was diluted with ethyl acetate (100 mL) and washed thoroughly with hydrochloric acid (0.1 M), water, and brine. The organic layer was dried over anhydrous sodium sulfate and concentrated *in vacuo*, and the resulting residue was used for the next step without further purification.

To a stirred solution of the residue from the previous step in THF (150 mL) was added $PPh_3$ (629 mg, 2.4 mmol, 4.0 equiv). DEAD (1.0 g, 2.4 mmol, 4.0 equiv, 40% wt in toluene) was added to the reaction dropwise. The reaction was complete upon the addition of DEAD. The reaction was concentrated *in vacuo*, and the resulting residue was purified by silica gel flash chromatography (DCM : acetone = 10:1-7:1), giving 340 mg of compound **S12** as white foam in a yield of 80% over two steps. $^1$H NMR (500 MHz, DMSO at 345K) δ 11.63 (s, 1H), 8.64 (d, *J* = 8.5 Hz, 1H), 8.47 (s, 1H), 8.29 (s, 1H), 7.46 (d, *J* = 8.2 Hz, 1H), 7.19 (dd, *J* = 8.2, 7.1 Hz, 1H), 7.05 (d, *J* = 6.9 Hz, 1H), 6.10 – 5.95 (m, 1H), 5.91 – 5.79 (m, 1H), 5.46 – 5.34 (m, 3H), 5.26 (dd, *J* = 10.5, 1.3 Hz, 1H), 5.08 (dd, *J* = 11.1, 3.4 Hz, 1H), 4.87 – 4.71 (m, 4H), 2.75 (s, 3H), 2.64 – 2.54 (m, 1H), 2.47 – 2.35 (m, 2H), 2.22 – 2.11 (m, 1H), 1.42 (s, 9H); $^{13}$C NMR (126 MHz, DMSO at 345K) δ 176.10, 171.98, 171.26, 162.40, 160.69, 160.64, 148.84, 146.54, 138.10, 132.92, 129.97, 129.86, 126.20, 125.57, 124.74, 124.48, 123.50, 118.81, 118.64, 114.37, 79.40, 66.44, 66.20, 65.57, 50.72, 31.77, 28.64, 12.85; **HRMS**: calculated for $C_{33}H_{36}N_5O_9S_2^+$ [M+H$^+$]: 710.19490; found: 710.19610.

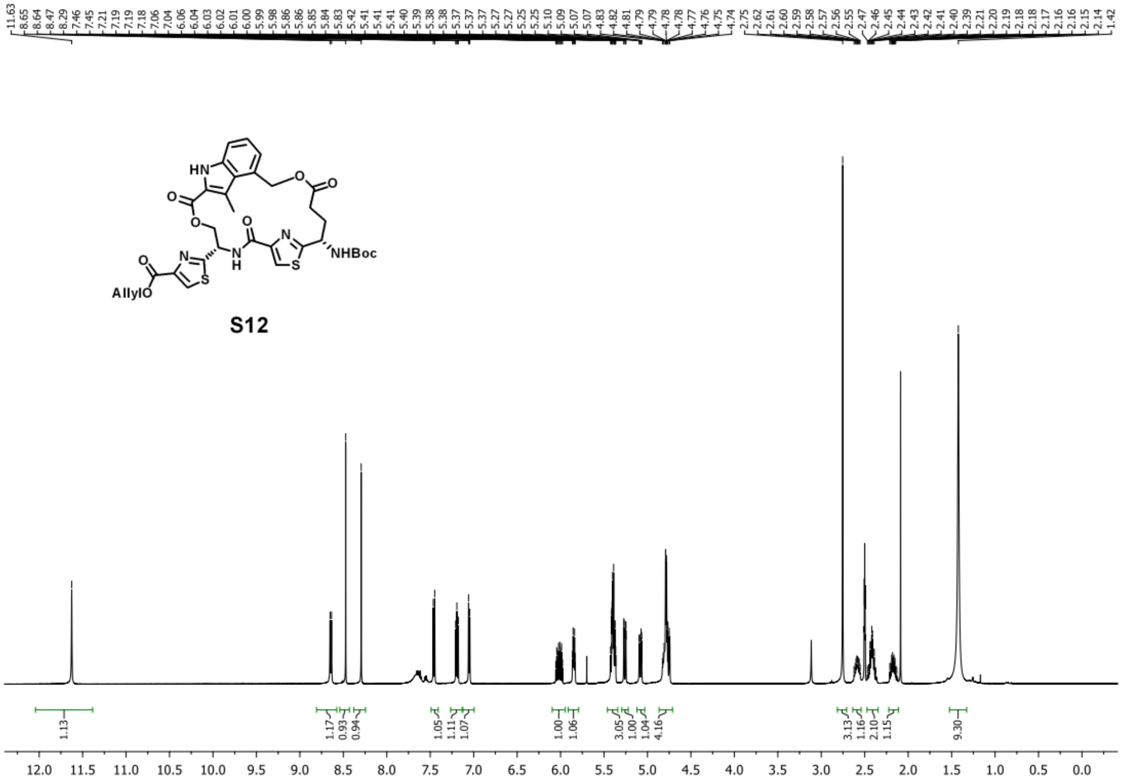
NHBoc
AllylO
S12

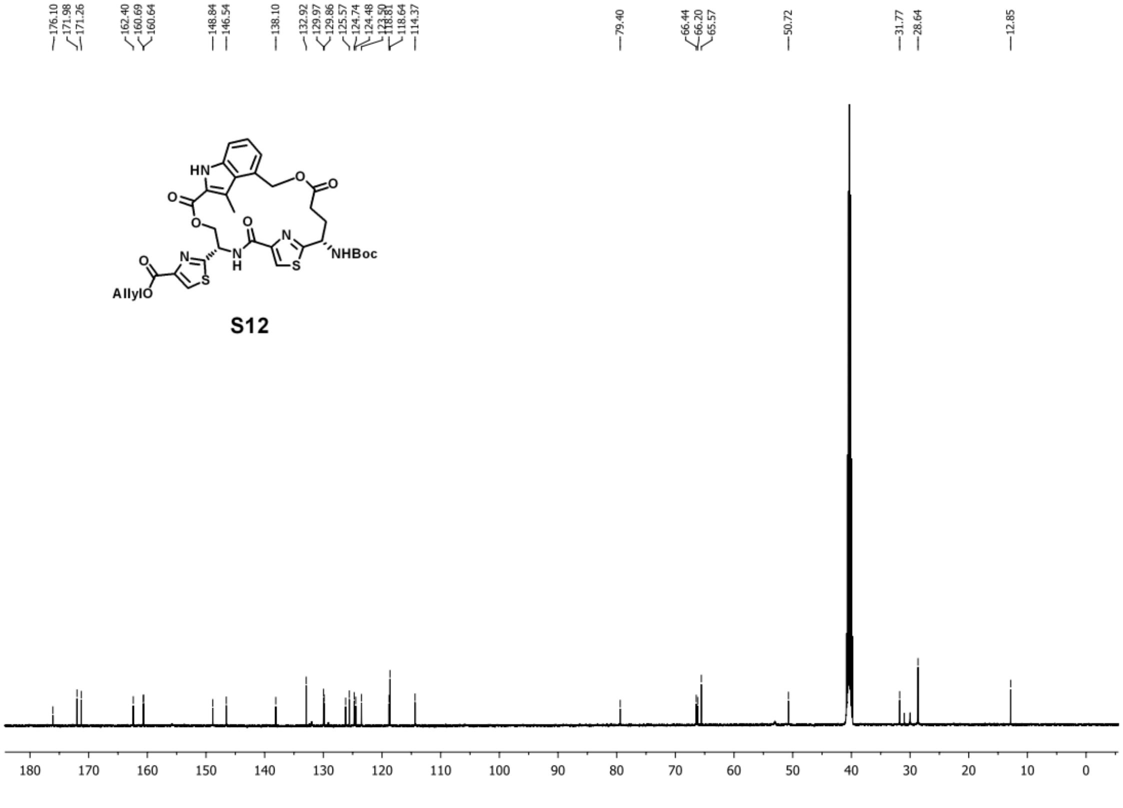
NHBoc
AllylO
S12

**Step j:**

Compound **S12** (98 mg, 138 µmol, 1.0 equiv) was added to 4M HCl in dioxane (10 mL). The resulting reaction was stirred at room temperature for 2 h. After completion, the reaction mixture was concentrated *in vacuo*, and the resulting white powder was used for the next step without further purification.

**Step k:**

Compound **S14** was synthesized according to the reported procedure.[2,24]

To a stirred solution of **S14** (2.0 g, 4.0 mmol, 1.0 equiv) in DMF (20 mL) was added TBAF (8.0 mL, 8.0 mmol, 2.0 equiv, 1M solution in THF). The resulting reaction was stirred at room temperature for 2 h. After completion, the reaction mixture was diluted with DCM (200 mL) and washed thoroughly with aqueous sodium hydroxide (0.1 M), saturated aqueous sodium bicarbonate, and brine. The organic layer was dried over anhydrous sodium sulfate and concentrated *in vacuo*, and the resulting residue was used for the next step without further purification.

To an ice-water cooled solution of the residue from the previous step in DMF (30 mL) was added Fmoc-Thr(O$^t$Bu)-OH (2.4 g, 6.0 mmol, 1.5 equiv), HATU (2.3 g, 6.0 mmol, 1.5 equiv), and TEA (1.4 mL, 10.0 mmol, 2.5 equiv). The resulting reaction was stirred at 0 °C for 1 h. After completion, the reaction mixture was diluted with ethyl acetate (200 mL) and washed thoroughly with hydrochloric acid (0.1 M), saturated aqueous sodium bicarbonate, and brine. The organic layer was dried over anhydrous sodium sulfate and concentrated *in vacuo*, and the resulting residue was purified by silica gel flash chromatography (hexanes : ethyl acetate = 10:1-5:1), giving 2.0 g of compound **S15** as white foam in a yield of 75% over two steps. $^1$H NMR (500

MHz, DMSO) δ 8.43 (s, 1H), 8.13 (d, *J* = 8.4 Hz, 1H), 7.89 (d, *J* = 7.5 Hz, 2H), 7.83 – 7.72 (m, 2H), 7.46 – 7.28 (m, 5H), 5.14 (dd, *J* = 8.5, 2.4 Hz, 1H), 4.38 – 4.22 (m, 6H), 4.20 – 4.11 (m, 1H), 4.00 – 3.92 (m, 1H), 1.30 (t, *J* = 7.1 Hz, 3H), 1.22 (s, 9H), 1.16 – 1.11 (m, 6H), 0.90 (s, 9H); $^{13}$C NMR (126 MHz, DMSO) δ 173.02, 170.19, 161.22, 156.46, 146.09, 144.37, 144.17, 141.20, 129.31, 128.11, 127.49, 125.86, 120.57, 74.95, 74.30, 69.15, 67.93, 66.37, 61.16, 59.23, 57.41, 47.16, 34.66, 31.43, 25.25, 22.54, 21.06, 20.48, 18.55, 14.66, 14.43; **HRMS**: calculated for $C_{36}H_{48}N_3O_7S^+$ [M+H$^+$]: 666.32075; found: 666.32169.

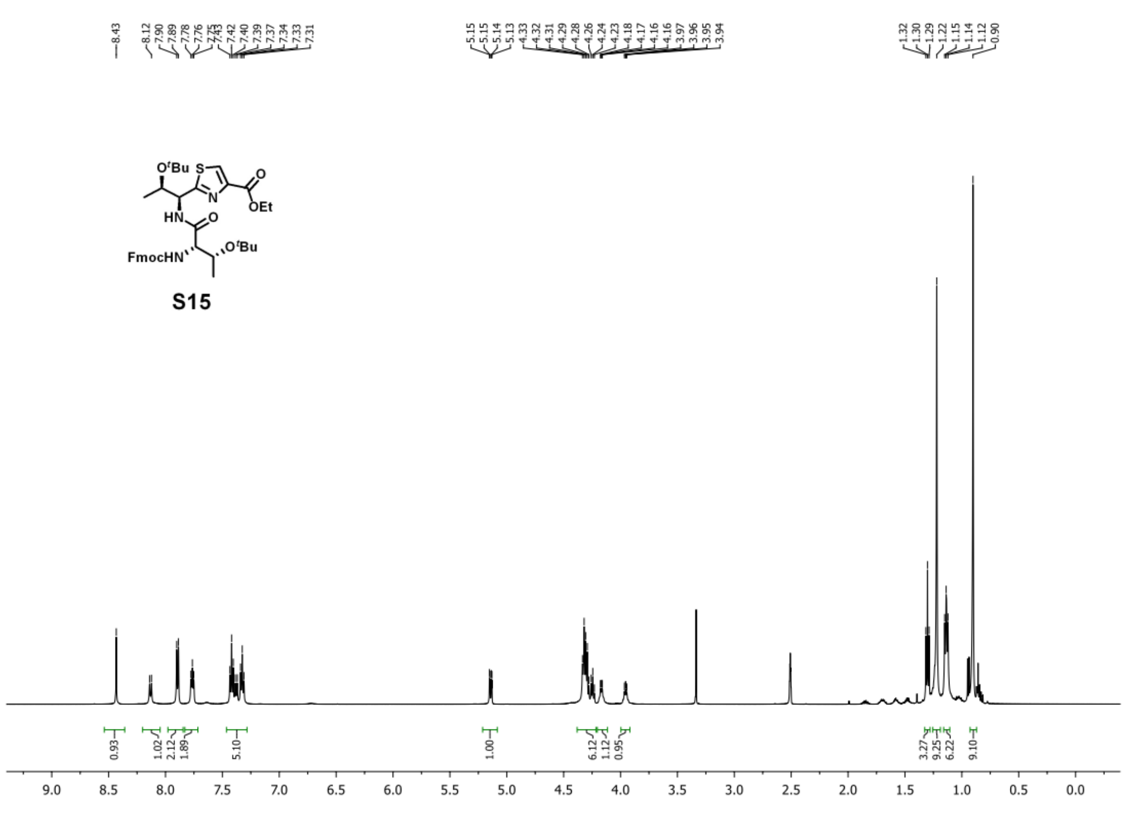

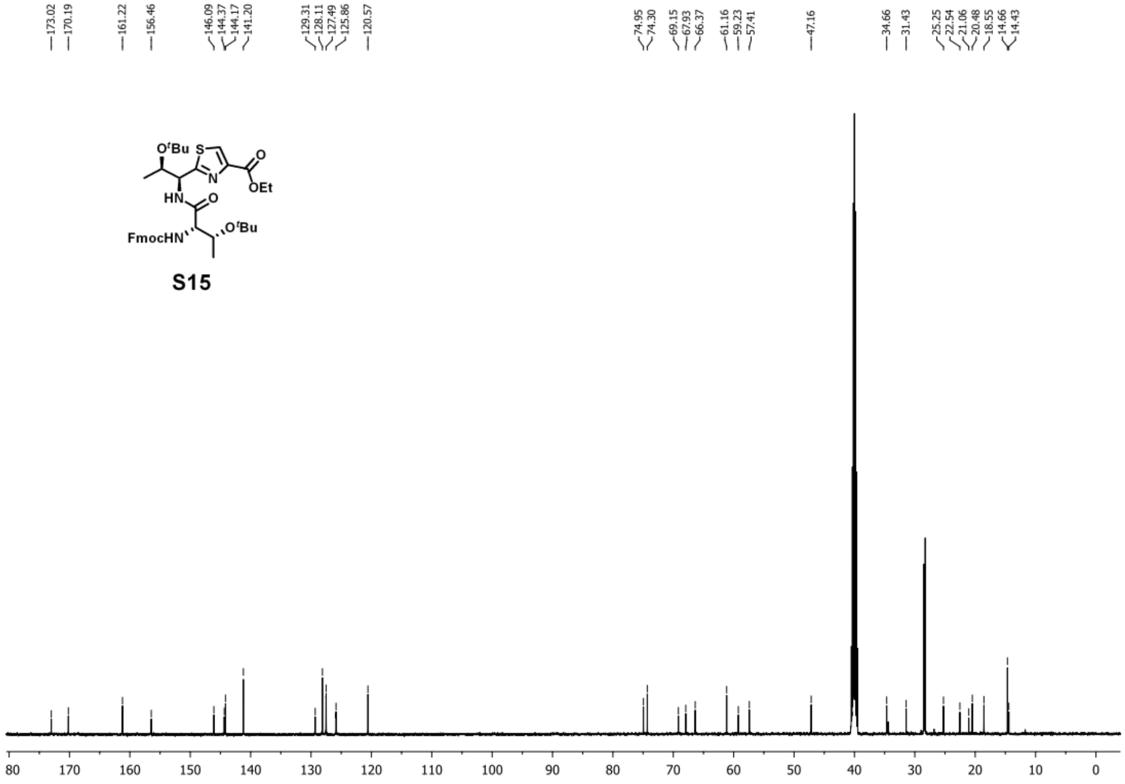


**Step l:**

To a stirred solution of **S15** (1.3 g, 2.0 mmol, 1.0 equiv) in DMF (20 mL) was added TBAF (4.0 mL, 4.0 mmol, 2.0 equiv, 1M solution in THF). The resulting reaction was stirred at room temperature for 2 h. After completion, the reaction mixture was diluted with DCM (150 mL) and washed thoroughly with aqueous sodium hydroxide (0.1 M), saturated aqueous sodium bicarbonate, and brine. The organic layer was dried over anhydrous sodium sulfate and concentrated *in vacuo*, and the resulting residue was used for the next step without further purification.

To an ice-water cooled solution of the residue from the previous step in DMF (20 mL) was added **S8** (852 mg, 2.6 mmol, 1.3 equiv), HATU (1.1 g, 3.0 mmol, 1.5 equiv), and TEA (1.4 mL, 10.0 mmol, 5.0 equiv). The resulting reaction was stirred at 0 °C for 1 h. After completion, the reaction mixture was diluted with ethyl acetate (200 mL) and washed thoroughly with hydrochloric acid (0.1 M), saturated aqueous sodium bicarbonate, and brine. The organic layer was dried over anhydrous sodium sulfate and concentrated *in vacuo*, and the resulting residue was purified by silica gel flash chromatography (hexanes : ethyl acetate = 10:1-3:1), giving 1.1 g of compound **S16** as yellowish foam in a yield of 70% over two steps. $^{1}$H NMR (500 MHz, DMSO at 345K) δ 8.39 (s, 1H), 8.26 (s, 1H), 8.10 (d, *J* = 8.4 Hz, 1H), 8.03 (d, *J* = 7.1 Hz, 1H), 5.28 (d, *J* = 5.1 Hz, 1H), 5.18 (dd, *J* = 8.4, 2.8 Hz, 1H), 4.61 (dd, *J* = 7.1, 3.7 Hz, 1H), 4.41 – 4.24 (m, 4H), 4.22 – 4.17 (m, 1H), 4.10 – 4.03 (m, 2H), 1.68 (s, 3H), 1.54 (s, 3H), 1.42 – 1.29 (m, 10H), 1.25 (s, 9H), 1.15 (d, *J* = 6.2 Hz, 3H), 1.10 (d, *J* = 6.3 Hz, 3H), 0.98 (s, 9H); $^{13}$C NMR (126 MHz, DMSO at 345K) δ 174.05, 171.85, 169.79, 161.23, 160.47, 146.25, 129.10, 124.87, 94.71, 74.93, 74.42, 69.05, 67.04, 61.06, 60.11, 59.22, 57.80, 57.57, 28.61, 28.42, 28.37, 20.20, 18.74, 14.60; **HRMS**: calculated for $C_{35}H_{56}N_5O_9S_2^+$ [M+H$^+$]: 754.35140; found: 754.35260.

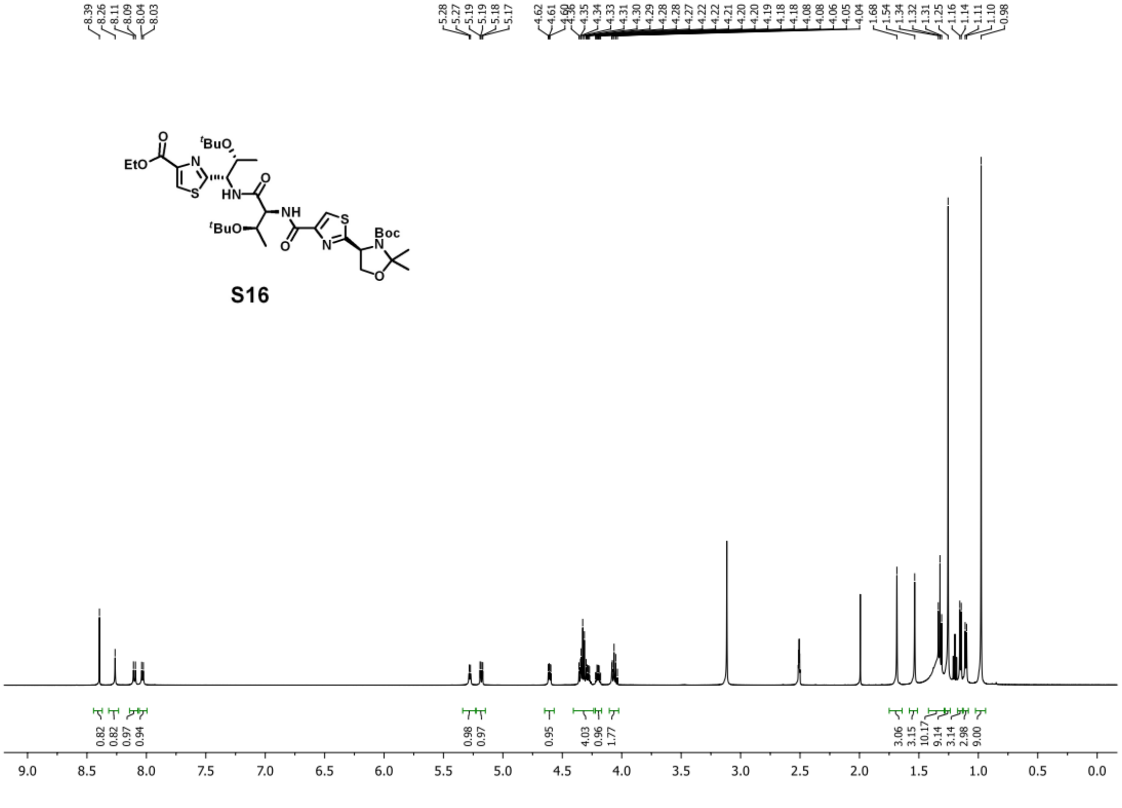

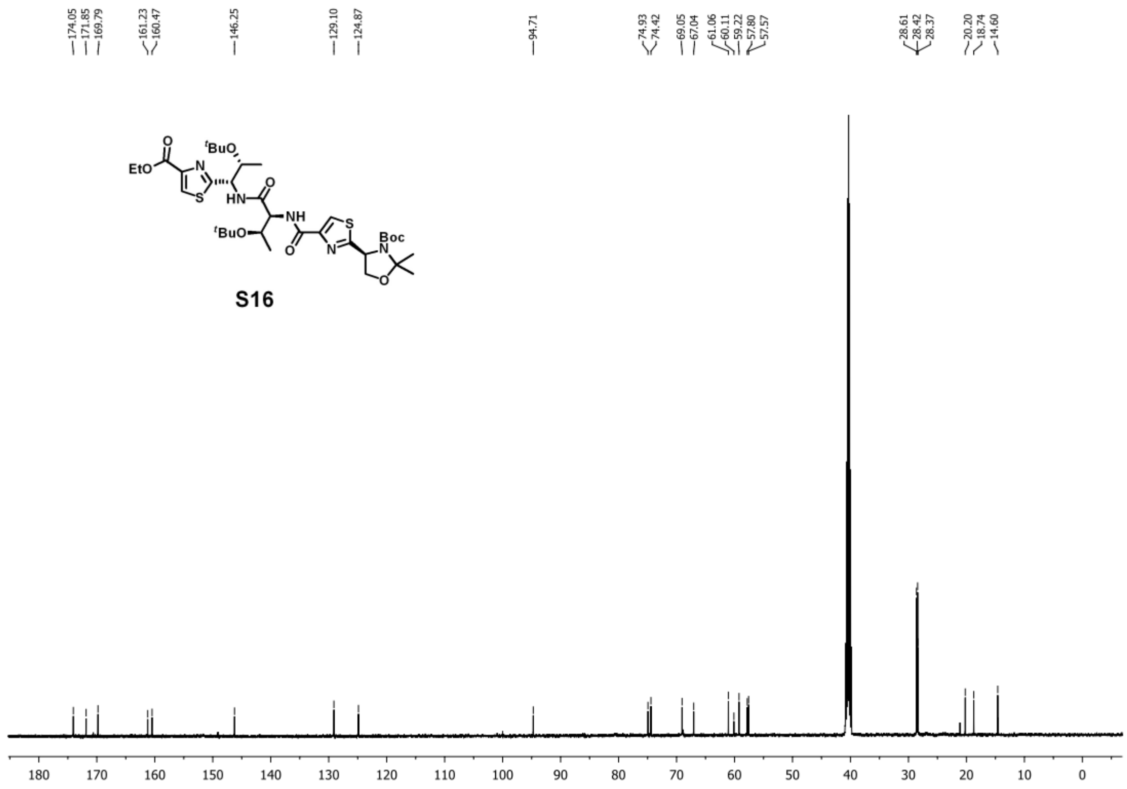


**Step m:**

To the stirred solution of **S16** (600 mg, 0.80 mmol, 1.0 equiv) in THF (10 mL)-MeOH (10 mL)-water (10 mL) was added sodium hydroxide (128 mg, 3.2 mmol, 4.0 equiv). The reaction mixture was stirred at room temperature for 2 h. After completion, the reaction mixture was diluted with water (100 mL) and washed with diethyl ether (100 mL). The aqueous layer was acidified by hydrochloric acid (1 M) to pH 3.0 and extracted with ethyl acetate (100 mL). The organic layer was washed with water and brine. The organic layer was dried over anhydrous sodium sulfate and concentrated *in vacuo*, giving 500 mg white foam.

The white foam from the previous step (100 mg, 138 μmol, 1.0 equiv) was dissolved in TFA (5 mL) and $H_2O$ (1 mL). The resulting reaction was stirred at room temperature for 1 h. After completion, the reaction mixture was concentrated *in vacuo*, and the resulting residue was used for the next step without further purification.

To an ice-water cooled solution of the residue from the previous step in acetonitrile (10 mL) and aqueous saturated sodium bicarbonate (10 mL) was added allyl chloroformate (59 µL, 550 µmol, 4.0 equiv). The reaction mixture was stirred at room temperature for 1 h. After completion, the reaction mixture was diluted with water (30 mL) and extracted with ethyl acetate (20 mL) three times. The aqueous layer was concentrated *in vacuo*, giving compound **S17** as a white powder. **S17** was used for the next step without purification.

**Step n:**

Compound **S13** (138 µmol, 1.0 equiv) and compound **S17** (138 µmol, 1.0 equiv) were dissolved in DMF (10 mL). HATU (52 mg, 138 µmol, 1.0 equiv) and TEA (100 µL, 690 µmol, 5.0 equiv) were added to the reaction mixture. The resulting reaction was stirred at room temperature for 1 h. After completion, the reaction mixture was diluted with ethyl acetate (100 mL) and washed with hydrochloric acid (0.1 M) and brine. The organic layer was dried over anhydrous sodium sulfate and concentrated *in vacuo*, and the resulting residue was purified by silica gel flash chromatography (DCM : methanol = 30:1-10:1), giving 87 mg of compound **S18** as a white powder in a yield of 55%. $^1$H NMR (500 MHz, DMSO at 345K) δ 11.65 (s, 1H), 9.00 (d, *J* = 8.6 Hz, 1H), 8.65 (d, *J* = 8.6 Hz, 1H), 8.49 (s, 1H), 8.35 (d, *J* = 8.4 Hz, 1H), 8.32 (s, 1H), 8.26 (s, 1H), 8.20 (s, 1H), 8.01 (d, *J* = 8.1 Hz, 1H), 7.74 (s, 1H), 7.47 (d, *J* = 8.3 Hz, 1H), 7.20 (t, *J* = 7.6 Hz, 1H), 7.07 (d, *J* = 7.0 Hz, 1H), 6.10 – 5.98 (m, 1H), 5.98 – 5.81 (m, 2H), 5.50 – 5.15 (m, 8H), 5.09 (dd, *J* = 11.1, 2.8 Hz, 1H), 4.95 (q, *J* = 12.4, 7.0 Hz, 1H), 4.84 – 4.73 (m, 3H), 4.64 (dd, *J* = 8.0, 3.6 Hz, 1H), 4.54 (d, *J* = 5.1 Hz, 2H), 4.35 – 4.28 (m, 1H), 4.27 – 4.19 (m, 1H), 3.90 (dd, *J* =

11.0, 5.0 Hz, 1H), 3.78 (dd, $J$ = 11.0, 6.8 Hz, 1H), 2.82 (s, 3H), 2.48 – 2.40 (m, 2H), 1.17 (d, $J$ = 6.3 Hz, 3H), 1.14 (d, $J$ = 6.2 Hz, 3H); $^{13}$C NMR (126 MHz, DMSO at 345K) δ 174.38, 172.89, 172.71, 171.93, 171.25, 170.81, 162.43, 161.48, 160.87, 160.69, 160.57, 149.72, 148.96, 148.61, 146.59, 138.12, 133.89, 132.92, 129.99, 129.89, 126.21, 125.92, 125.27, 124.76, 124.53, 124.50, 123.53, 118.83, 118.65, 117.55, 114.38, 68.20, 67.26, 66.44, 66.37, 65.59, 65.22, 63.79, 58.39, 57.29, 56.54, 55.22, 51.36, 50.60, 32.05, 29.52, 20.44, 20.39, 12.92; **HRMS**: calculated for $C_{49}H_{53}N_{10}O_{15}S_4^+$ [M+H$^+$]: 1149.25692; found: 1149.25880.

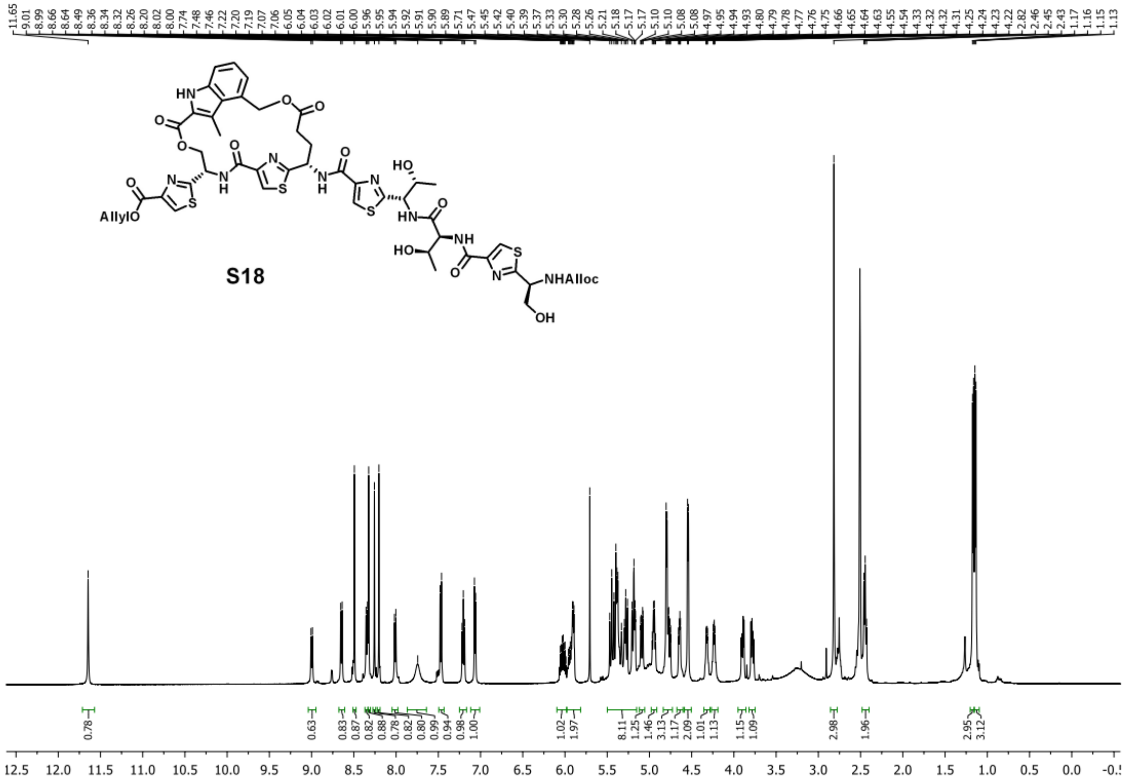

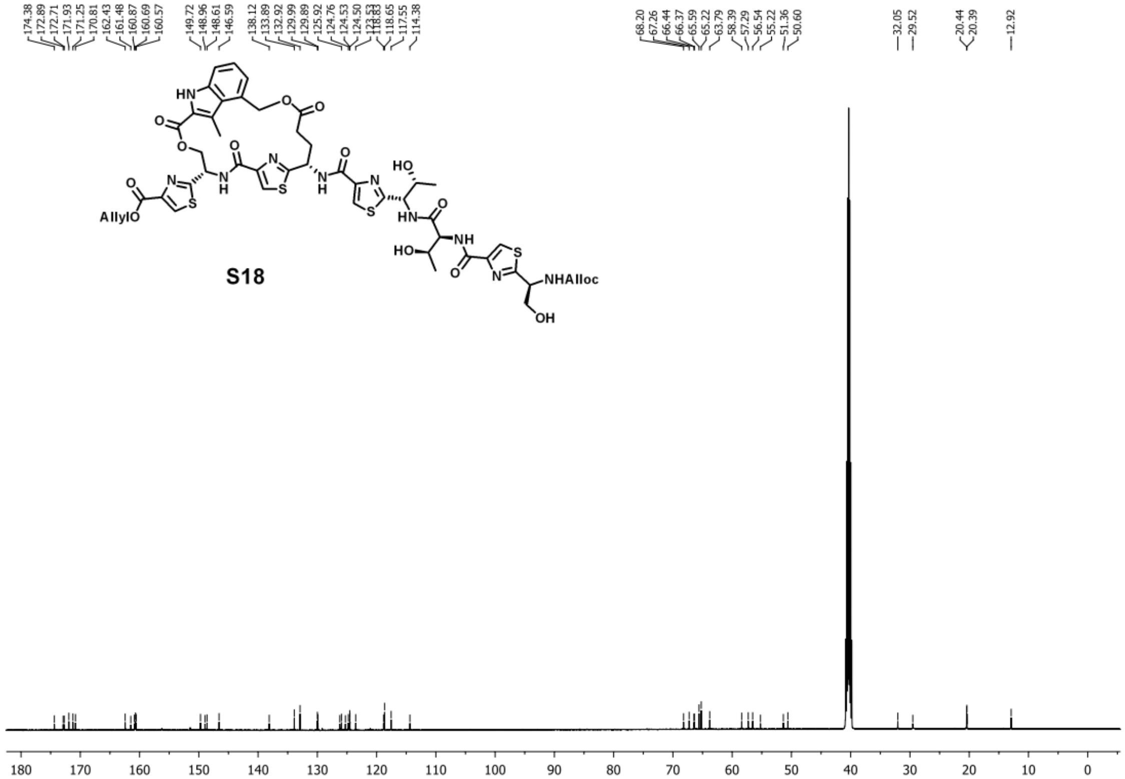


**Step o:**

To a stirred solution of **S18** (50 mg, 43 µmol, 1.0 equiv) in THF (10 mL) was added $Pd(PPh_3)_4$ (25 mg, 22 µmol, 0.5 equiv) and $PhSiH_3$ (53 µL, 430 µmol, 10.0 equiv) The resulting reaction was stirred at room temperature for 1 h. After completion, the reaction mixture was diluted with water (40 mL) and extracted with ethyl acetate (20 mL) three times. The aqueous layer was lyophilized. The resulting residue was dissolved in 50% $H_2O$-acetonitrile (5 mL) and purified using Agilent 5 Prep-C18 50×21.2 mm column on Agilent 1260 Infinity II Prep HPLC system. The HPLC purification was performed using solvent A (water) and solvent B (acetonitrile) as follows:

| Time (min) | A% | B% | Flow rate (ml/min) |
|---|---|---|---|
| 0.00 | 85.0 | 15.0 | 30.00 |
| 1.00 | 85.0 | 15.0 | 30.00 |
| 7.00 | 10.0 | 90.0 | 30.00 |

SRC eluted from 3.9 min to 4.7 min. The fractions containing SRC were pooled and lyophilized, giving 9 mg of SRC in a yield of 20%. $^{1}$H NMR (500 MHz, DMSO) δ 11.67 (s, 1H), 8.96 (d, *J* = 8.6 Hz, 1H), 8.53 (s, 1H), 8.39 (d, *J* = 8.4 Hz, 1H), 8.31 (s, 1H), 8.24 (s, 1H), 8.14 (s, 1H), 8.04 (d, *J* = 8.2 Hz, 1H), 7.46 (d, *J* = 8.3 Hz, 1H), 7.21 – 7.16 (m, 2H), 7.04 (d, *J* = 6.9 Hz, 2H), 5.46 (d, *J* = 11.8 Hz, 1H), 5.41 – 5.32 (m, 2H), 5.19 – 5.06 (m, 2H), 4.78 – 4.67 (m, 1H), 4.62 (dd, *J* = 8.2, 3.7 Hz, 1H), 4.37 – 4.28 (m, 1H), 4.27 – 4.15 (m, 2H), 3.76 (dd, *J* = 10.6, 4.3 Hz, 1H), 3.56 (dd, *J* = 10.6, 6.6 Hz, 1H), 2.70 (s, 1H), 2.46 – 2.38 (m, 2H), 1.15 (d, *J* = 6.3 Hz, 3H), 1.10 (d, *J* = 6.3 Hz, 3H); $^{13}$C NMR (126 MHz, DMSO) δ 178.10, 174.33, 173.17, 172.08, 171.05, 162.58, 161.47, 161.03, 160.45, 149.61, 148.87, 148.45, 137.94, 129.85, 126.17, 126.07, 125.52, 124.81, 124.46, 123.54, 118.61, 114.35, 68.19, 67.43, 66.63, 66.42, 63.55, 58.02, 57.10, 56.09, 51.06, 50.25, 32.07, 29.36, 20.66, 20.56, 13.00; **HRMS**: calculated for $C_{42}H_{45}N_{10}O_{13}S_4^{+}$ [M+H$^{+}$]: 1025.20449; found: 1025.20620.

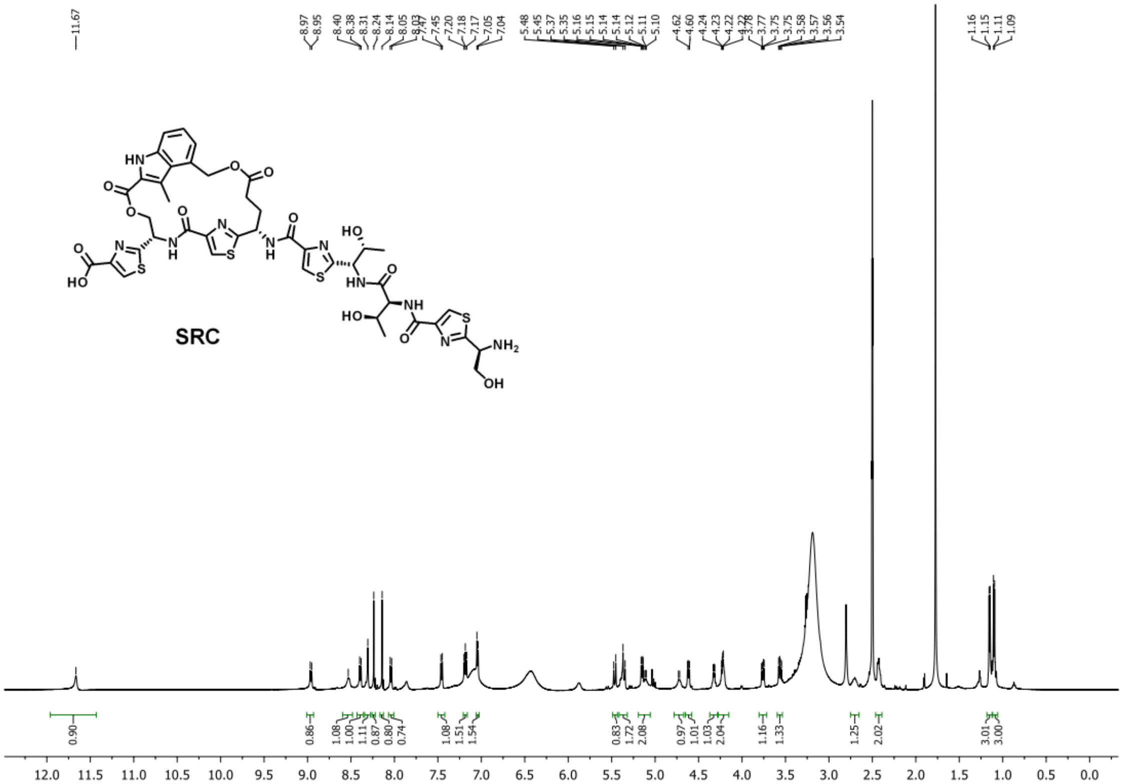
SRC

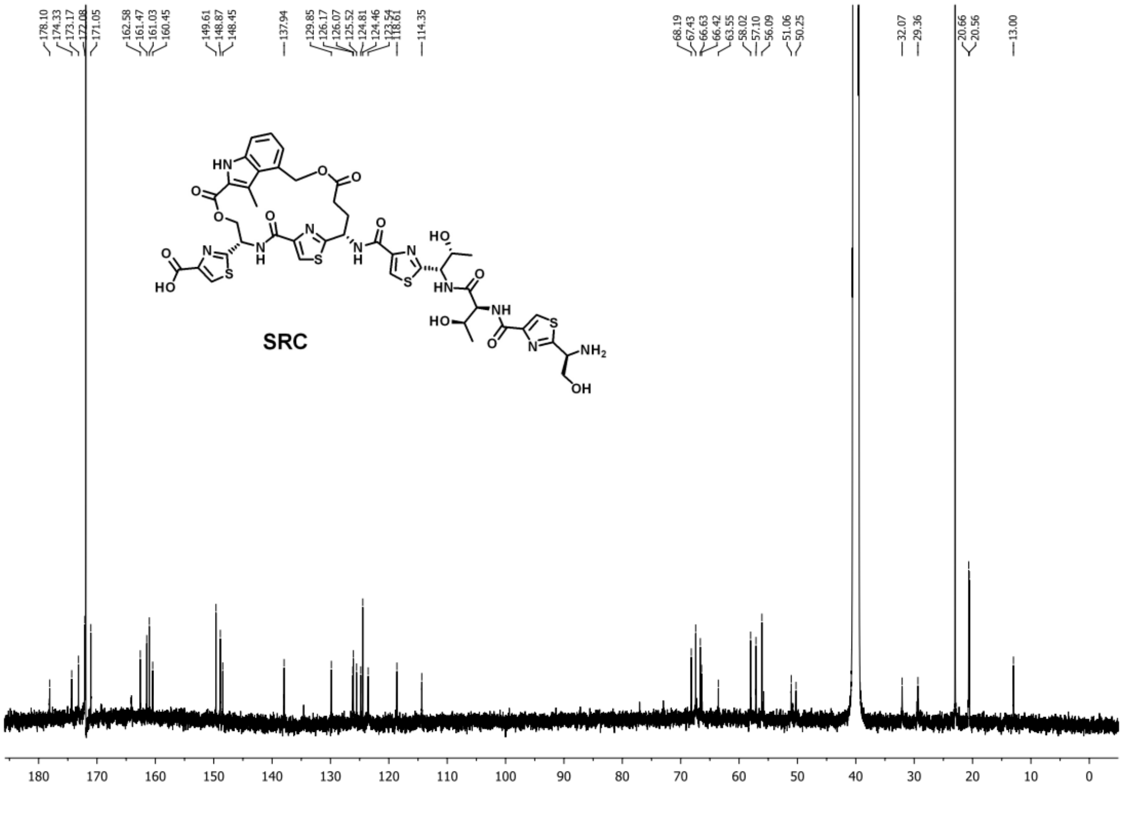
SRC

### 7. Synthesis of deuterated substrate peptides for EPR studies

#### 7.1. Synthesis of 1*H*-indole-2-carboxylic-5-*d* acid.

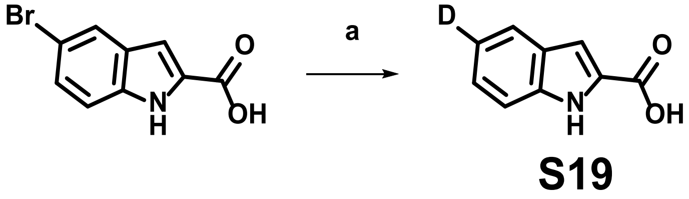


**Conditions and reagents:** a) Deuterium gas, Pd/C, MeOH, rt, quantitative.

To the solution of 5-bromo-1*H*-indole-2-carboxylic acid (1.0 g, 4.2 mmol, 1 equiv) in methanol (100 mL) was added Pd/C (10% on carbon, 100 mg). The reaction mixture was stirred vigorously under deuterium gas overnight. After filtration to remove Pd/C, the resulting solution was concentration *in vacuo* to give compound **S19** as a yellowish solid in a quantitative yield. $^{1}$H NMR (500 MHz, DMSO) δ 12.94 (s, 1H), 11.76 (s, 1H), 7.64 (s, 1H), 7.44 (d, $J$ = 8.3 Hz, 1H), 7.24 (d, $J$ = 8.3 Hz, 1H), 7.09 (d, $J$ = 1.3 Hz, 1H); $^{13}$C NMR (126 MHz, DMSO) δ 163.28, 137.71, 128.86, 127.31, 124.64, 122.29, 120.17 (t, $J_{C\text{-}D}$ = 23.9 Hz), 112.94, 107.76; **HRMS**: calculated for $C_9H_7DNO_2^+$ [M+H$^+$]: 163.06123; found: 163.06107.

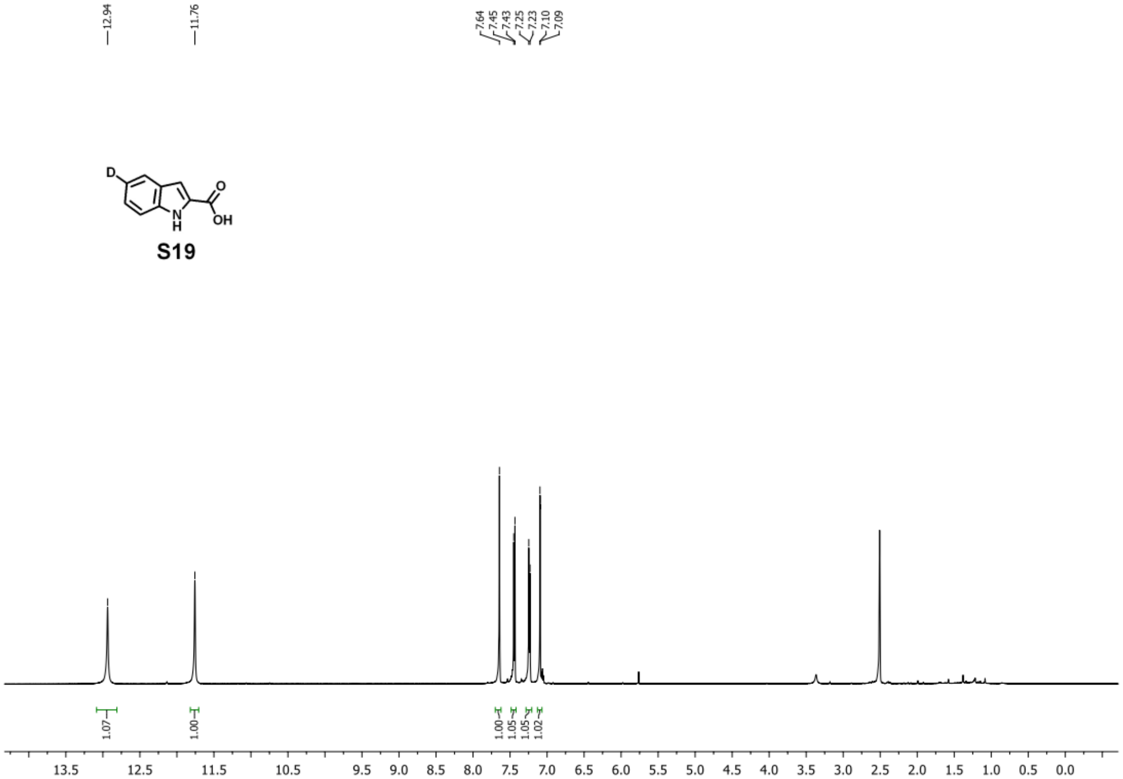
12.94
11.76
7.64
7.45
7.43
7.25
7.23
7.10
7.09
D
O
OH
N
H
1.07
1.00
1.00
1.05
1.05
1.02
13.5 12.5 11.5 10.5 9.5 9.0 8.5 8.0 7.5 7.0 6.5 6.0 5.5 5.0 4.5 4.0 3.5 3.0 2.5 2.0 1.5 1.0 0.5 0.0

S19

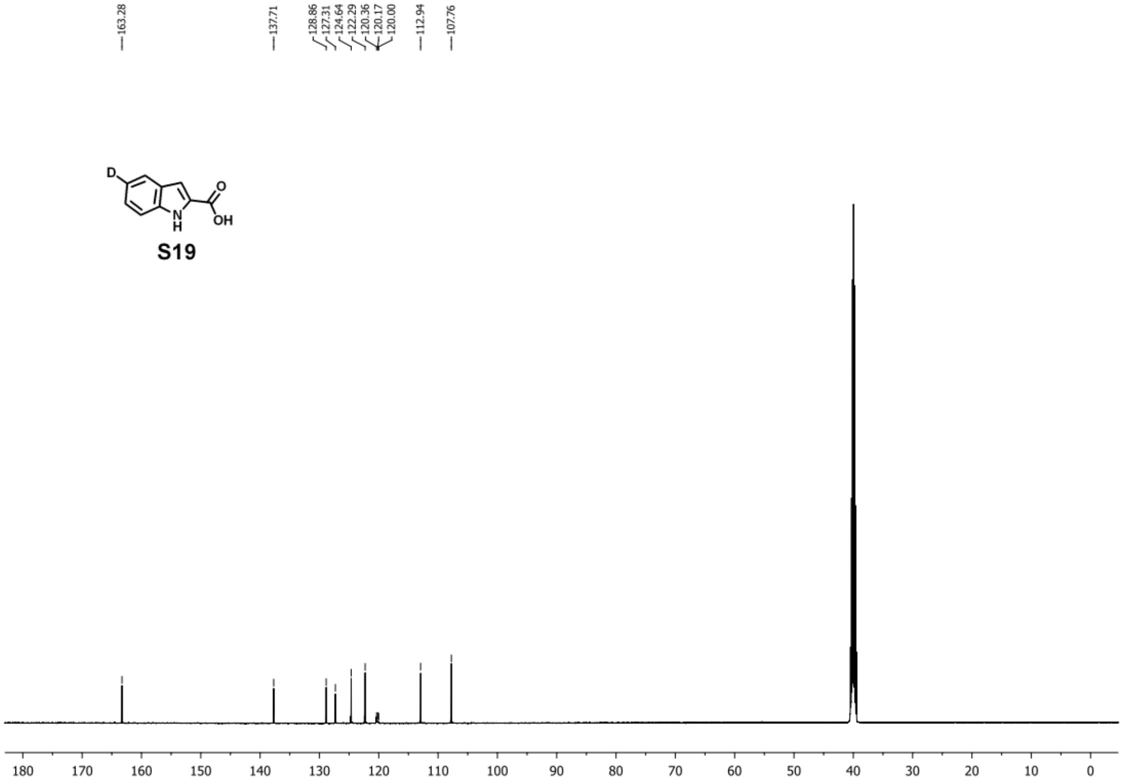
163.28
137.71
128.86
127.31
124.64
122.29
120.36
120.17
120.00
112.94
107.76
D
O
OH
N
H
180 170 160 150 140 130 120 110 100 90 80 70 60 50 40 30 20 10 0

S19

### 7.2. Synthesis of 1*H*-indole-2-carboxylic-4-*d* acid.

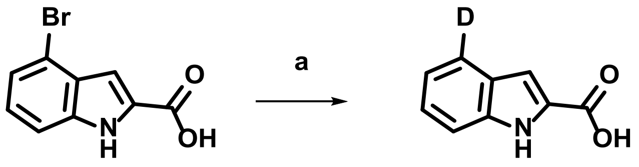

**S20**

**Conditions and reagents:** a) Deuterium, Pd/C, MeOH, rt, quantitative.

To the solution of 4-bromo-1*H*-indole-2-carboxylic acid (1.0 g, 4.2 mmol, 1 equiv) in methanol (100 mL) was added Pd/C (10% on carbon, 100 mg). The reaction mixture was stirred vigorously under deuterium gas overnight. After filtration to remove Pd/C, the resulting solution was concentrated *in vacuo* to give compound **S20** as a yellowish solid in a quantitative yield. $^{1}$H NMR shows that the isotopic purity is 90%, with 10% protium incorporated. $^{1}$H NMR (500 MHz, DMSO) δ 12.91 (s, 1H), 11.74 (s, 1H), **7.64 (d, *J* = 8.0 Hz, 0.1H),** 7.44 (d, *J* = 8.3 Hz, 1H), 7.30 – 7.21 (m, 1H), 7.14 – 7.03 (m, 2H); $^{13}$C NMR (126 MHz, DMSO) δ 163.28, 137.71, 128.90, 127.25, 124.72, 122.13 (t, $J_{C\text{-}D}$ = 23.9 Hz), 120.30, 112.94, 107.70; **HRMS**: calculated for $C_9H_7DNO_2^+$ [M+H$^+$]: 163.06123; found: 163.06097.

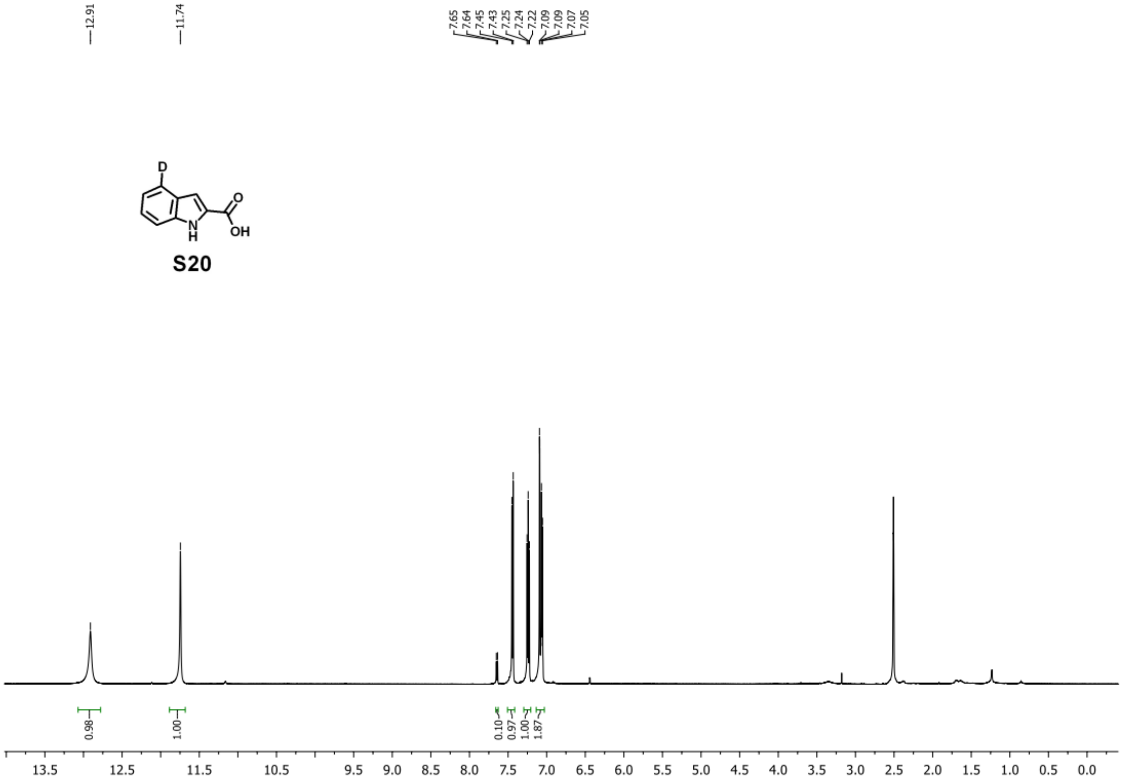
12.91
11.74
7.65
7.64
7.45
7.43
7.25
7.24
7.22
7.09
7.09
7.07
7.05
0.98
1.00
0.10
0.97
1.00
1.87
13.5
12.5
11.5
10.5
9.5
9.0
8.5
8.0
7.5
7.0
6.5
6.0
5.5
5.0
4.5
4.0
3.5
3.0
2.5
2.0
1.5
1.0
0.5
0.0
D
H
N
O
OH
S20

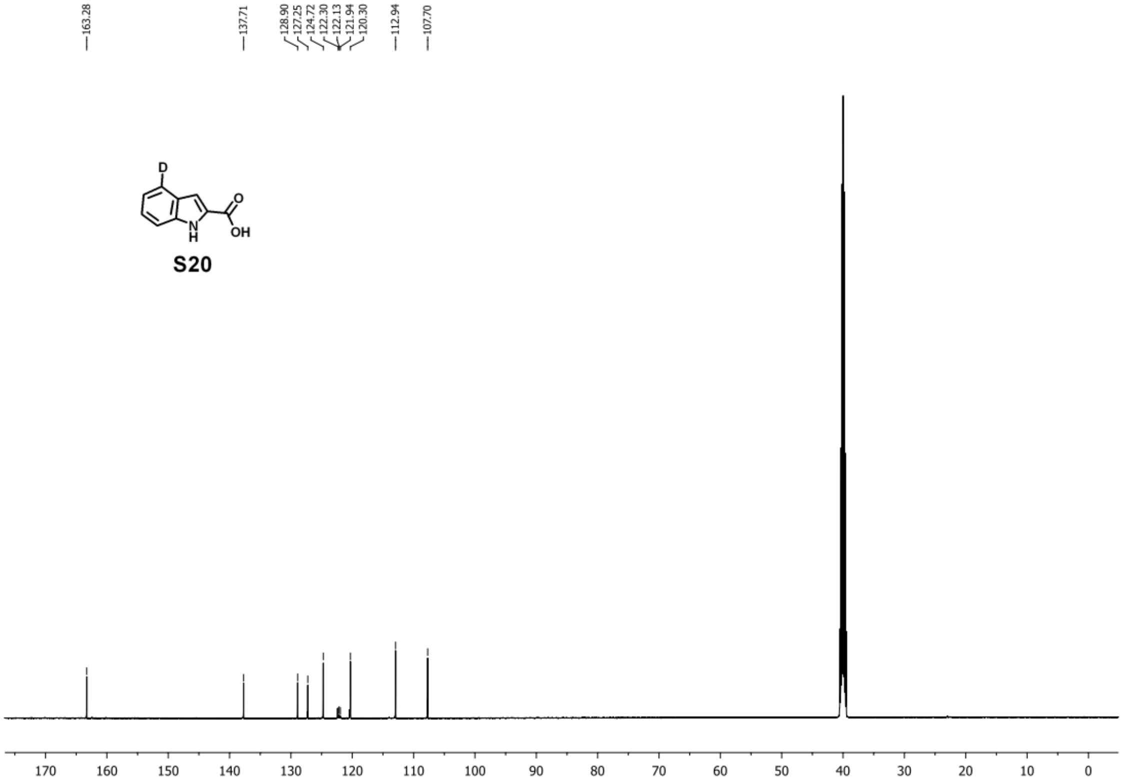
163.28
137.71
128.90
127.25
124.72
122.30
122.13
121.94
120.30
112.94
107.70
170
160
150
140
130
120
110
100
90
80
70
60
50
40
30
20
10
0
D
H
N
O
OH
S20

### 7.3. Synthesis of 1H-indole-2-carboxylic-4,5,6,7-$d_4$ acid.

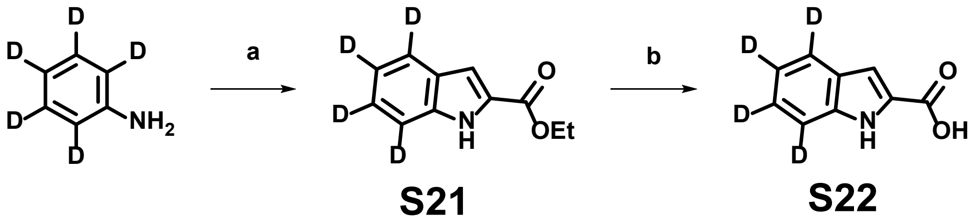

**Conditions and reagents:** a) ethyl pyruvate, $Pd(OAc)_2$, 4A MS, AcOH, DMSO, $O_2$, 70 ºC, 12h, 40%; b) NaOH, THF, MeOH, $H_2O$, rt, 1h.

**Step a:**

Scaling up the reaction will result in a lower yield. Thus, twenty reactions in the following scale were set up in parallel and pooled for workup and purification:

Per-deuterated aniline (39.3 mg, 0.4 mmol, 1.0 equiv), $Pd(OAc)_2$ (9.0 mg, 0.04 mmol, 0.1 equiv), ethyl pyruvate (92.8 mg, 0.8 mmol, 2.0 equiv), acetic acid (96.0 mg, 1.6 mmol, 4.0 equiv), and 4A MS (80 mg) were added to DMSO (2.0 mL) in a 5 mL vial. The reaction was bubbled with oxygen for 1 min and then sealed. The reaction mixture was stirred at 70 ºC for 12 h. After completion, the reaction mixture was combined and diluted with ethyl acetate (250 mL) and washed thoroughly with 0.1 M HCl, saturated aqueous sodium bicarbonate, water, and brine. The organic layer was dried over anhydrous sodium sulfate and concentrated *in vacuo*, and the resulting residue was purified by silica gel flash chromatography (hexanes : ethylacetate = 20:1-5:1), giving 620 mg of compound **S21** as a yellowish solid in a yield of 40%.$^1$H NMR (500 MHz, DMSO) δ 11.88 (s, 1H), 7.15 (d, *J* = 2.0 Hz, 1H), 4.35 (q, *J* = 7.1 Hz, 2H), 1.35 (t, *J* = 7.1 Hz, 3H); $^{13}$C NMR (126 MHz, DMSO) δ 161.80, 137.78, 127.81, 127.10, 124.60 (t, $J_{C-D}$ = 26.5 Hz), 122.12 (t, $J_{C-D}$ = 22.7 Hz), 120.13 (t, $J_{C-D}$ = 23.9 Hz), 112.67 (t, $J_{C-D}$ = 23.9 Hz), 108.10, 60.88, 14.77; **HRMS**: calculated for $C_{11}H_8D_4NO_2^+$ [M+H$^+$]: 194.11136; found: 194.11106.

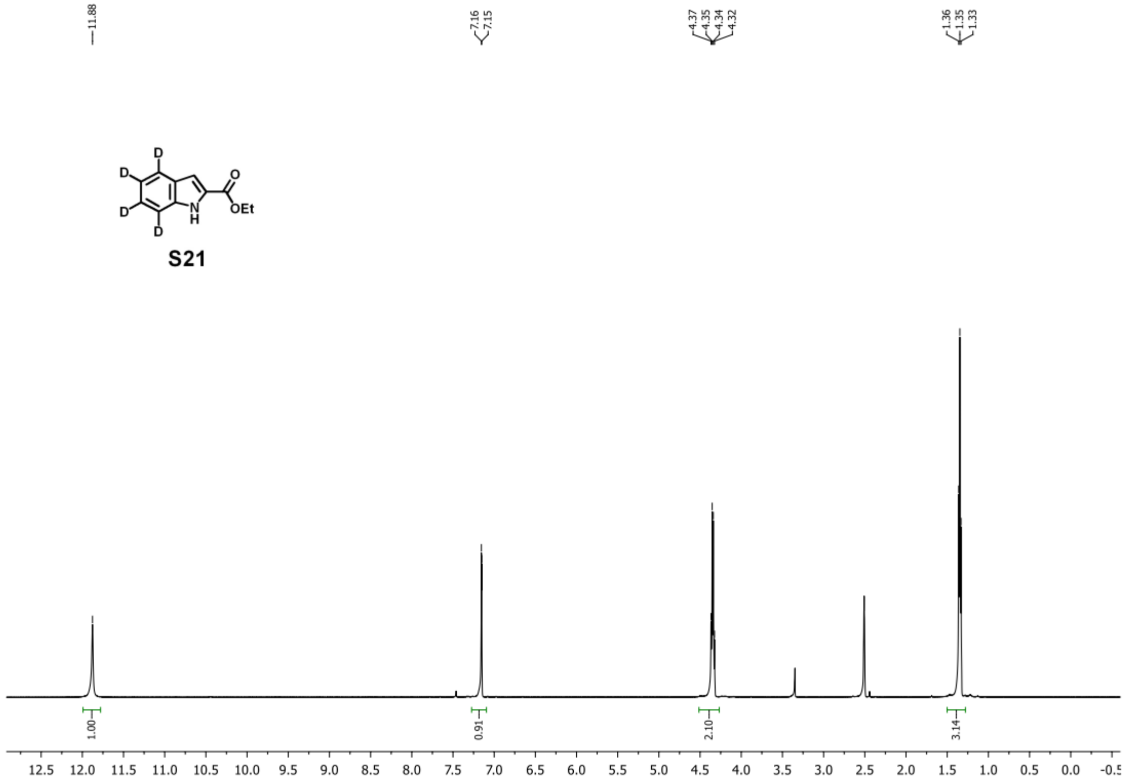
11.88
7.16
7.15
4.37
4.35
4.34
4.32
1.36
1.35
1.33
D
D
D
D
O
OEt
N
H
S21
1.00
0.91
2.10
3.14
12.5 12.0 11.5 11.0 10.5 10.0 9.5 9.0 8.5 8.0 7.5 7.0 6.5 6.0 5.5 5.0 4.5 4.0 3.5 3.0 2.5 2.0 1.5 1.0 0.5 0.0

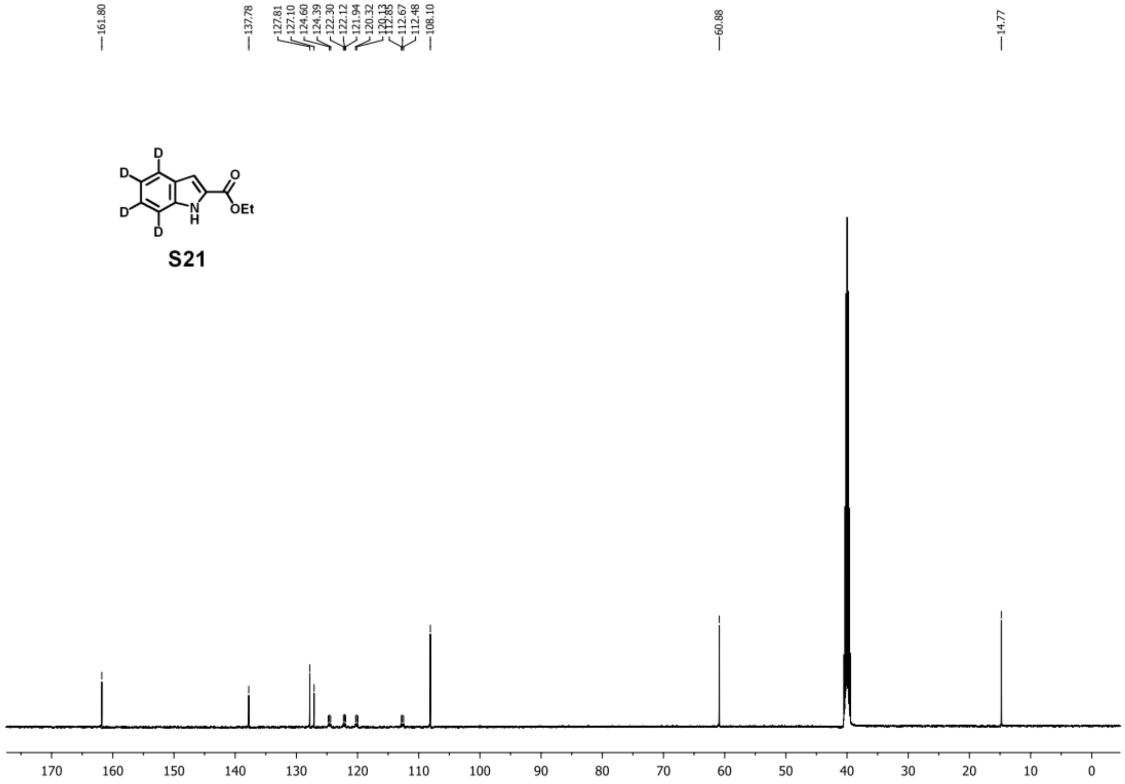
161.80
137.78
127.81
127.10
124.60
124.39
122.30
122.12
121.94
120.32
108.10
112.67
112.48
60.88
14.77
D
D
D
D
O
OEt
N
H
S21
170 160 150 140 130 120 110 100 90 80 70 60 50 40 30 20 10 0

**Step b:**

To the stirred solution of **S21** (300 mg, 1.6 mmol, 1.0 equiv) in THF (10 mL)-MeOH (10 mL)-water (10 mL) was added sodium hydroxide (320 mg, 8.0 mmol, 5 equiv). The reaction mixture was stirred at room temperature for 1 h. After completion, the reaction mixture was diluted with ethyl acetate (100 mL) and washed thoroughly with 0.1 M HCl and brine. The organic layer was dried over anhydrous sodium sulfate and concentrated *in vacuo*, and the resulting compound **S22** was used for the next step without further purification. $^{1}$H NMR (500 MHz, DMSO) δ 12.94 (s, 1H), 11.76 (s, 1H), 7.11 (d, $J$ = 2.2 Hz, 1H); $^{13}$C NMR (126 MHz, DMSO) δ 163.31, 137.67, 128.87, 127.26, 124.26 (t, $J_{C\text{-}D}$ = 22.5 Hz), 122.01 (t, $J_{C\text{-}D}$ = 23.8 Hz), 119.94 (t, $J_{C\text{-}D}$ = 21.3 Hz), 112.58 (t, $J_{C\text{-}D}$ = 22.5 Hz), 107.75; **HRMS**: calculated for $C_9H_4D_4NO_2^+$ [M+H$^+$]: 166.08006; found: 166.07985.

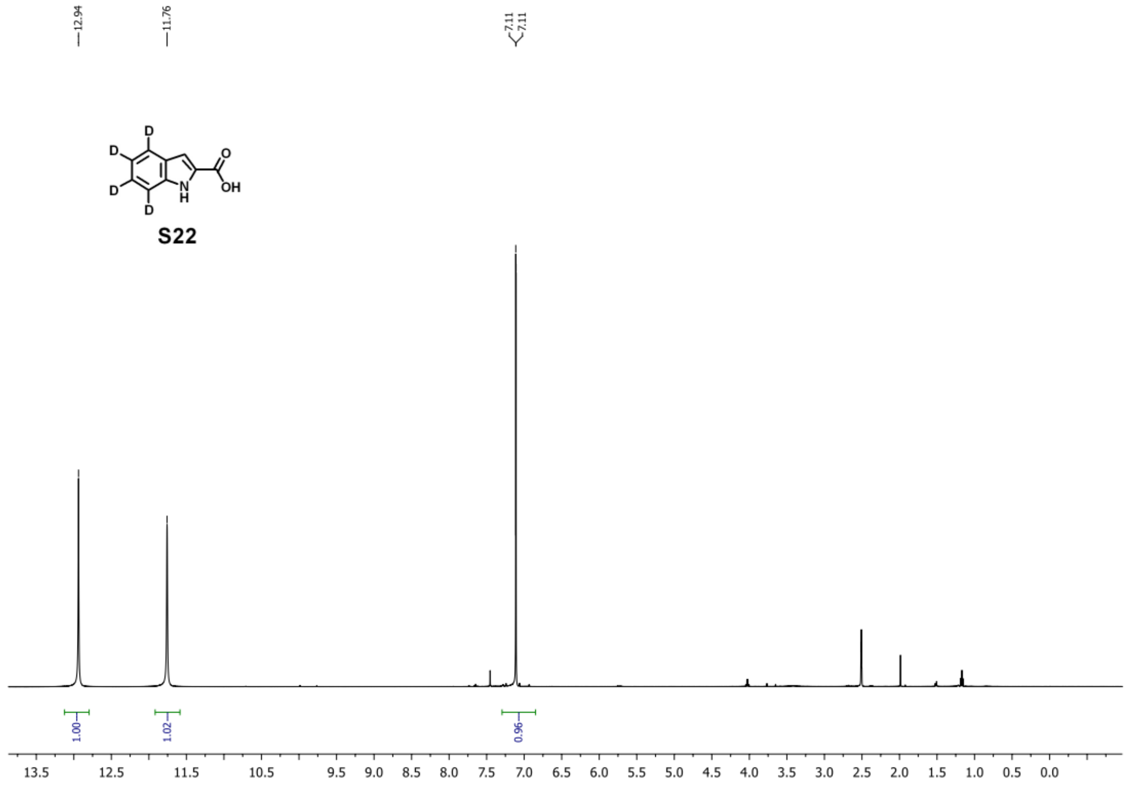

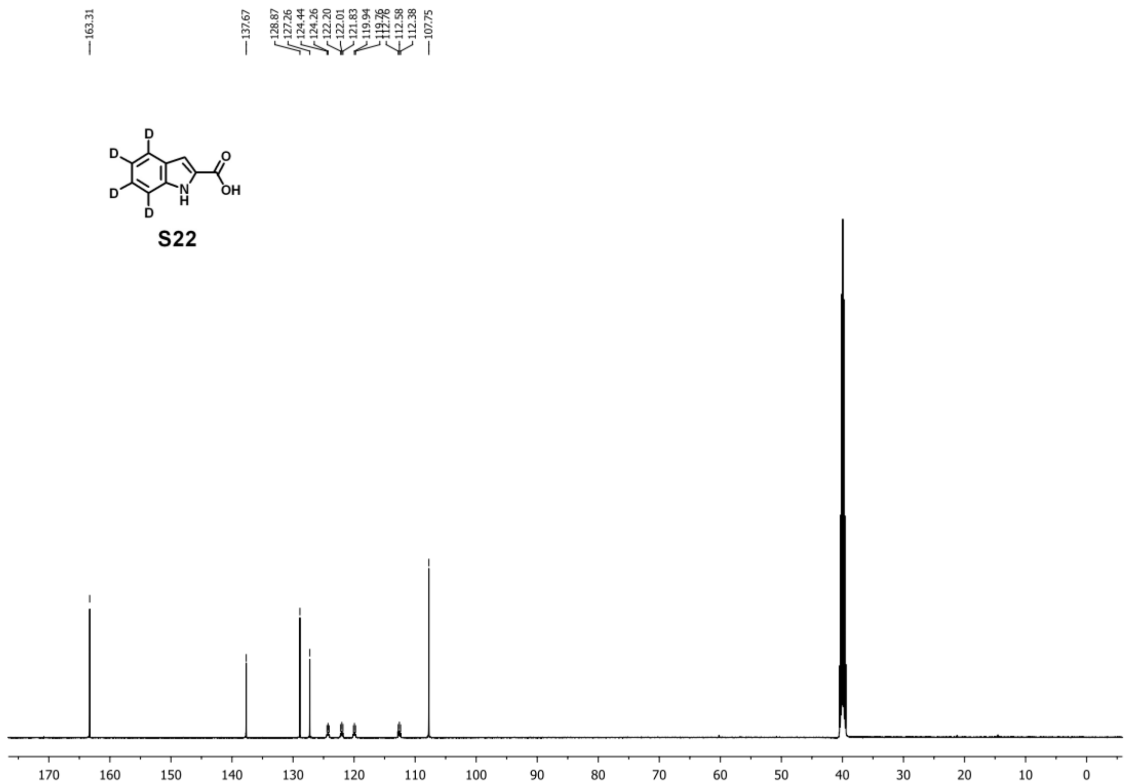


### 7.4. Synthesis of 1*H*-indole-2-carboxylic-4,6-$d_2$ acid.

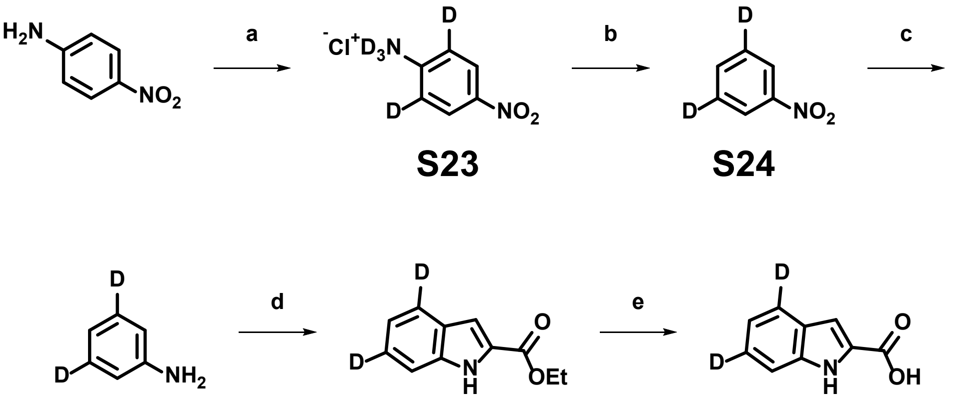


**Conditions and reagents:** a) DCl, $D_2O$, 110 °C, 24 h; b) $NaNO_2$, $CHCl_3$, $H_2O$, AcOH, rt, 2h; c) $H_2$, Pd/C, MeOH, rt, 3h; d) ethyl pyruvate, $Pd(OAc)_2$, 4A MS, AcOH, DMSO, $O_2$, 70 °C, 12h, 52%; e) NaOH, THF, MeOH, $H_2O$, rt, 1h.

**Step a:**

4-Nitroaniline (10.0 g, 72.5 mmol, 1 equiv) was added to deuterium chloride solution in $D_2O$ (35 wt.%, 5 mL) and $D_2O$ (20 mL). The resulting solution was sealed in a 100 mL heavy wall pressure vessel and stirred at 110 ºC for 24 hours. After cooling to the room temperature, the reaction mixture was lyophilized to give brownish solid **S23** without further purification in a quantitative yield. $^{1}H$ NMR (500 MHz, DMSO) δ 7.97 (s, 2H); $^{13}C$ NMR (126 MHz, DMSO) δ 152.77, 137.54, 126.53, 114.48; **HRMS**: calculated for $C_6H_5D_2N_2O_2^+$ [M+H$^+$]: 141.06276; found: 141.06247.

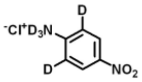


**S23**

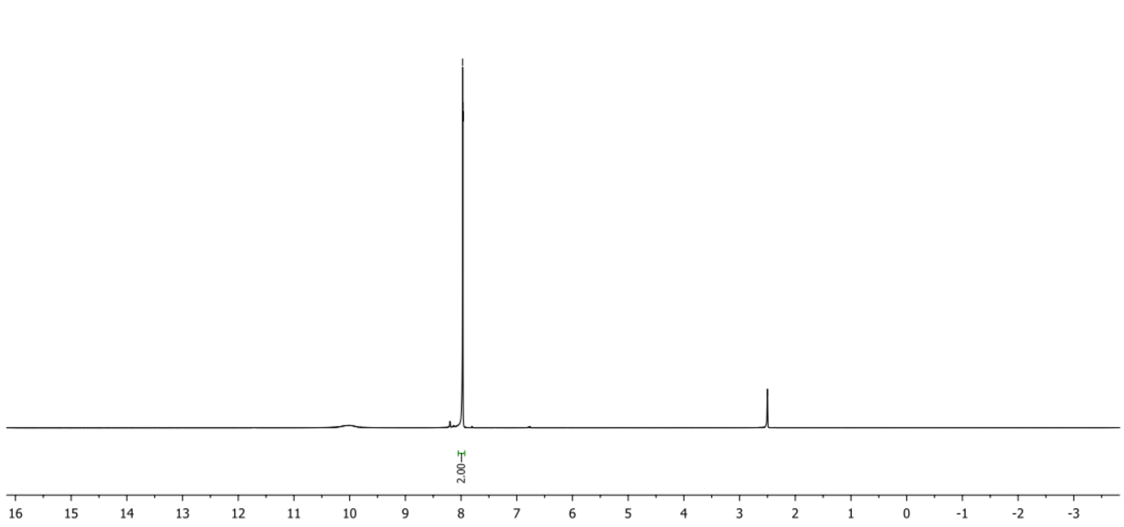

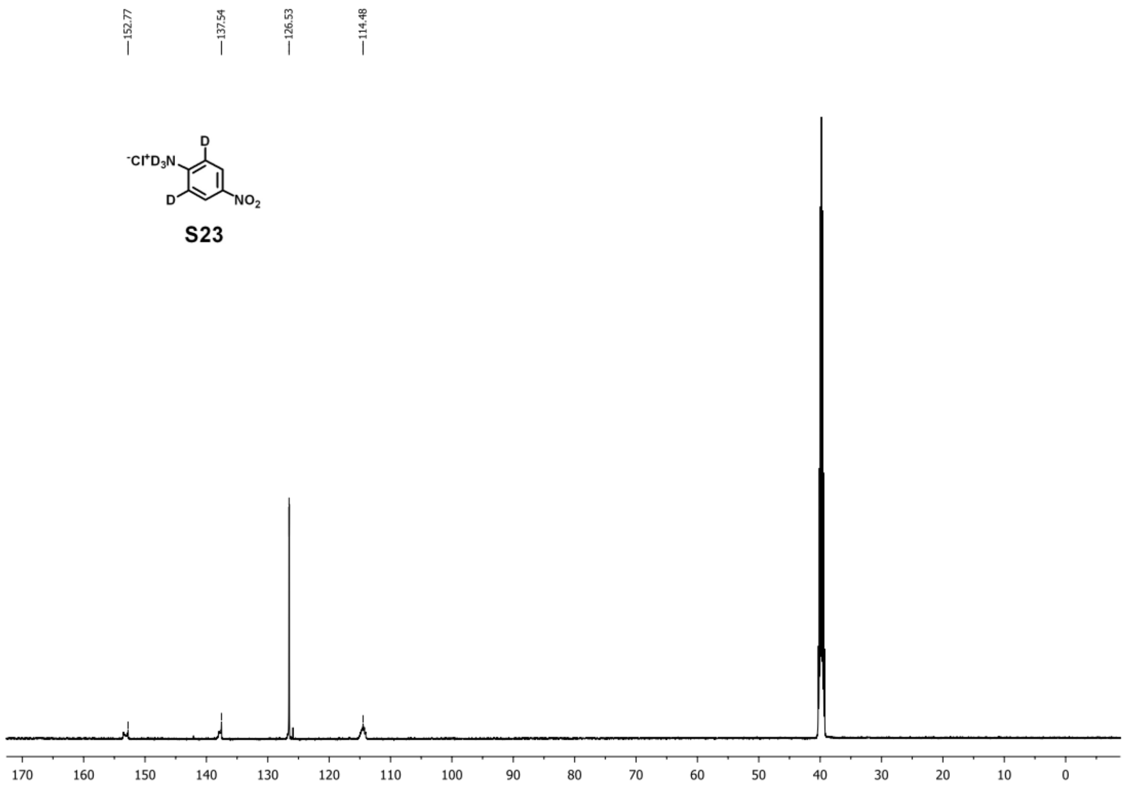


**Step b:**

To a stirred solution of **S23** (1.0 g, 5.6 mmol, 1.0 equiv) in chloroform (25 mL), water (25 mL), and acetic acid (5 mL) was added sodium nitrite (1.7 g, 24.6 mmol, 4.5 equiv). The reaction was stirred at room temperature for 2h. After completion, the reaction mixture was diluted with dichloromethane (100 mL) and washed thoroughly with 0.1 M HCl, saturated aqueous sodium bicarbonate, and brine. The organic layer was dried over anhydrous sodium sulfate and concentrated *in vacuo*, and the resulting compound **S24** was used for next step without further purification.

**Step c:**

To the solution of **S24** (500 mg, 4.0 mmol, 1 equiv) in THF (30 mL) was added Pd/C (10%, 50 mg). The reaction mixture was stirred vigorously under hydrogen gas overnight. After filtration to remove Pd/C, the resulting solution was concentrated *in vacuo* to give brown oil **S25** in a quantitative yield. $^{1}$H NMR (500 MHz, $CDCl_3$) δ 6.85 (s, 1H), 6.75 (s, 2H); $^{13}$C NMR (126 MHz, $CDCl_3$) δ 146.40, 129.10 (t, $J_{C\text{-}D}$ = 23.9 Hz), 118.44, 115.18; **HRMS**: calculated for $C_6H_6D_2N^+$ [$M+H^+$]: 96.07768; found: 96.07786.

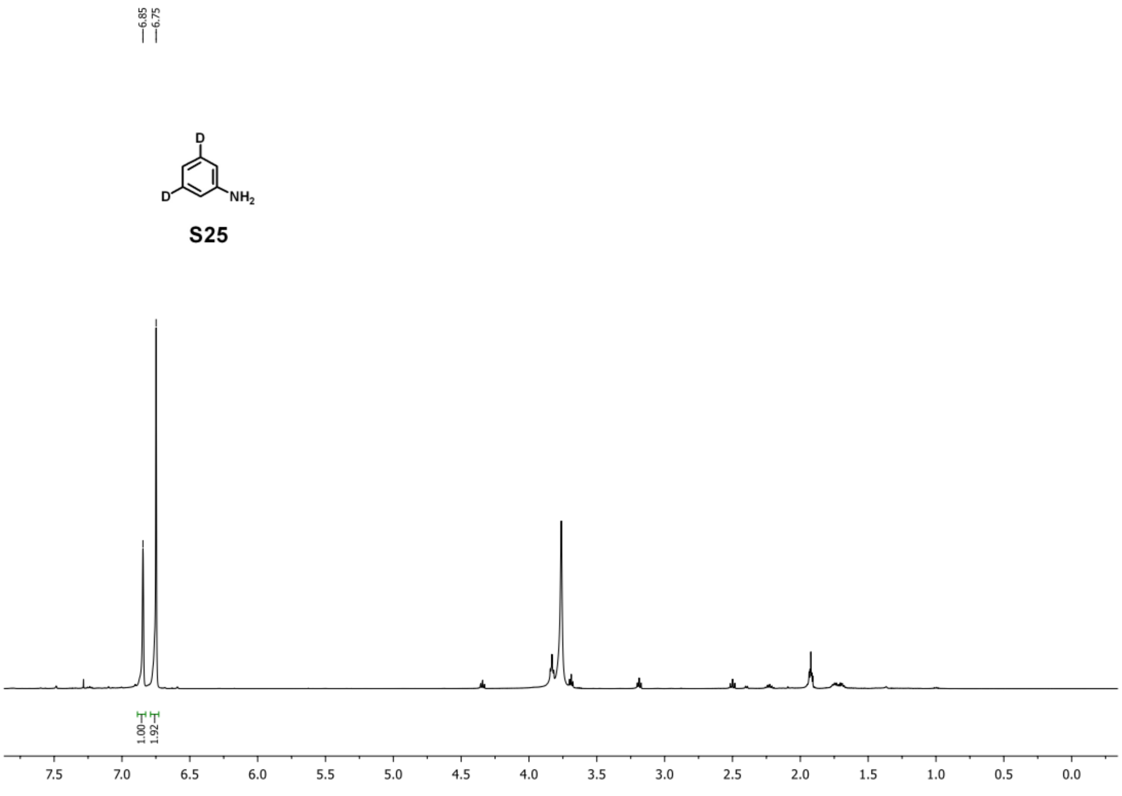

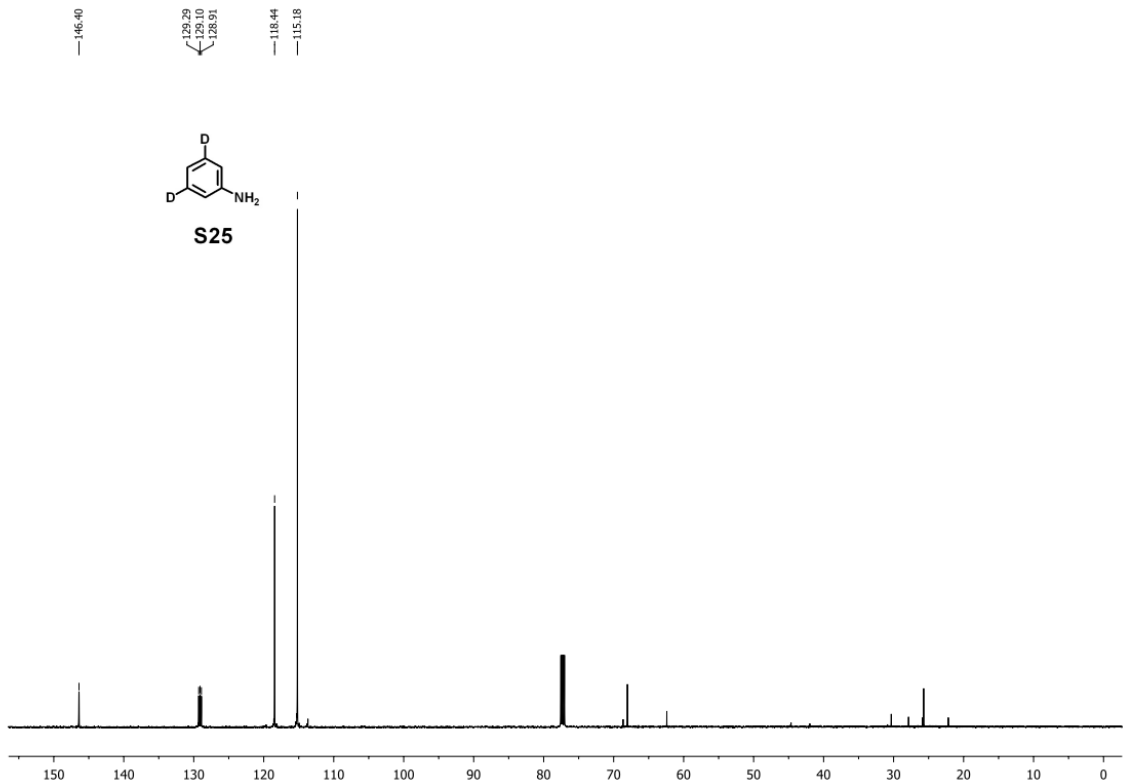


**Step d:**

<u>Scaling up the reaction will result in a lower yield. Thus, twenty reactions in the following scale were set up in parallel and pooled for work up and purification:</u>

To each reaction: **S25** (38 mg, 0.4 mmol, 1.0 equiv), $Pd(OAc)_2$ (9.0 mg, 0.04 mmol, 0.1 equiv), ethyl pyruvate (92.8 mg, 0.8 mmol, 2.0 equiv), acetic acid (96.0 mg, 1.6 mmol, 4.0 equiv), and 4A MS (80 mg) were added to DMSO (2.0 mL) in a 5 mL vial. The reaction was bubbled with oxygen for 1 min then sealed. The reaction mixture was stirred at 70 °C for 12 h. After completion, the reaction mixture was combined and diluted with ethyl acetate (250 mL) and washed thoroughly with 0.1 M HCl, saturated aqueous sodium bicarbonate, water, and brine. The organic layer was dried over anhydrous sodium sulfate and concentrated *in vacuo*, and the resulting residue was purified by silica gel flash chromatography (hexanes : ethylacetate = 20:1-

5:1), giving 400 mg of compound **S26** as yellowish solid in a yield of 52%. $^{1}H$ NMR (500 MHz, DMSO) δ 11.90 (s, 1H), 7.48 (s, 1H), 7.16 (s, 1H), 7.08 (s, 1H), 4.35 (q, $J$ = 7.1 Hz, 2H), 1.34 (t, $J$ = 7.1 Hz, 3H); $^{13}C$ NMR (126 MHz, DMSO) δ 161.81, 137.86, 127.82, 127.12, 124.79 (t, $J_{C\text{-}D}$ = 25.2 Hz), 122.24 (t, $J_{C\text{-}D}$ = 23.9 Hz), 120.39, 112.93, 108.10, 60.87, 14.76; **HRMS**: calculated for $C_{11}H_{10}D_2NO_2^+$ [M+H$^+$]: 192.09881; found: 192.09860.

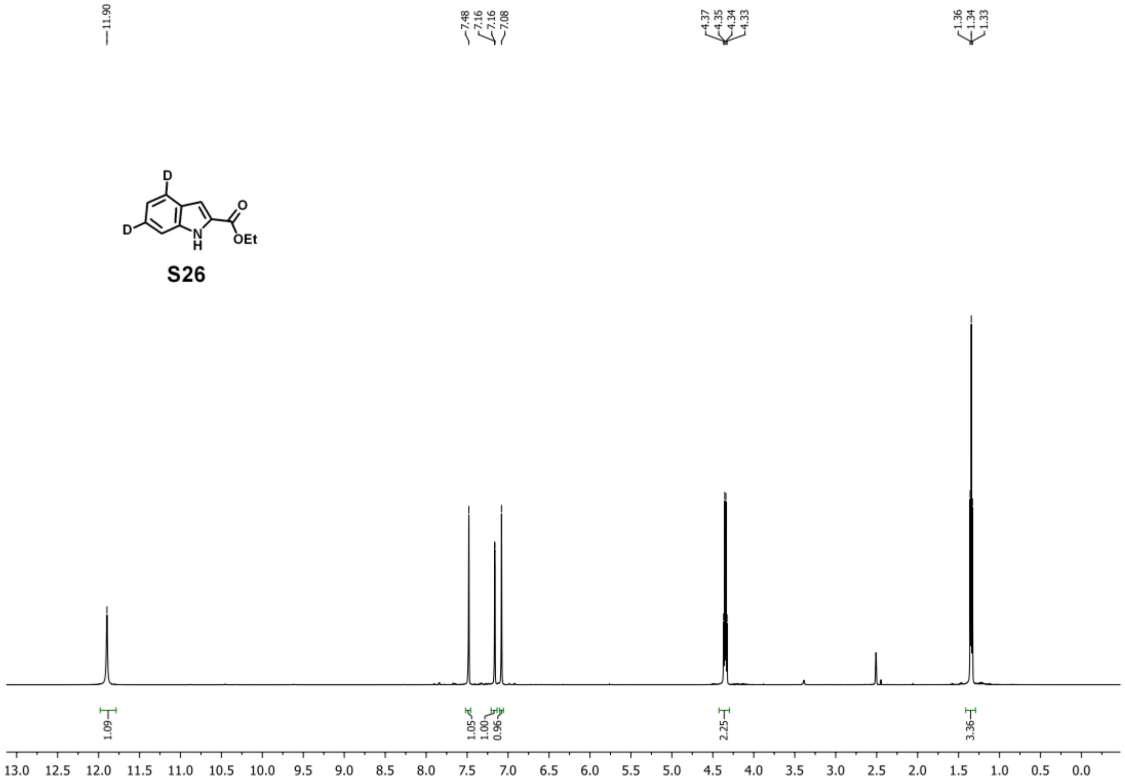

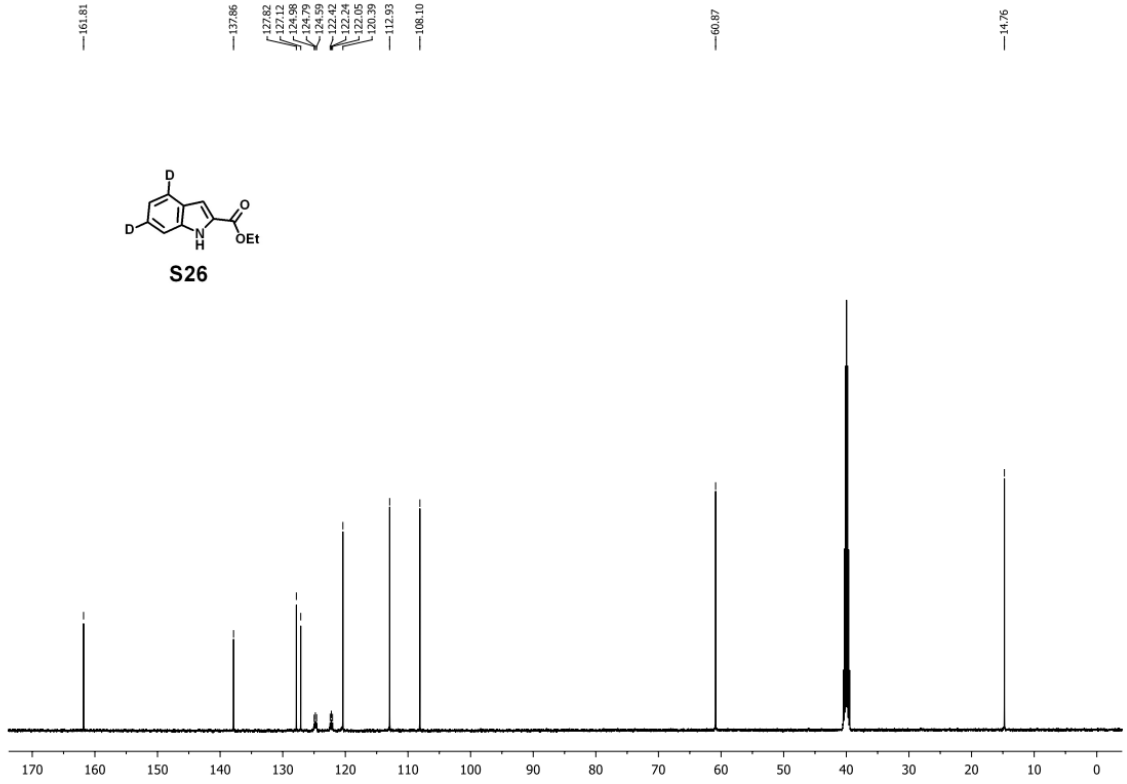


**Step e:**

To the stirred solution of **S26** (300 mg, 1.6 mmol, 1.0 equiv) in THF (10 mL)-MeOH (10 mL)-water (10 mL) was added sodium hydroxide (320 mg, 8.0 mmol, 5 equiv). The reaction mixture was stirred at room temperature for 1 h. After completion, the reaction mixture was diluted with ethyl acetate (100 mL) and washed thoroughly with 0.1 M HCl and brine. The organic layer was dried over anhydrous sodium sulfate and concentrated *in vacuo*, and the resulting compound **S27** was used for next step without further purification. $^{1}$H NMR (500 MHz, DMSO) δ 12.94 (s, 1H), 11.75 (s, 1H), 7.45 (s, 1H), 7.13 – 7.08 (m, 1H), 7.06 (s, 1H); $^{13}$C NMR (126 MHz, DMSO) δ 163.30, 137.71, 128.91, 127.25, 124.45 (t, $J_{C\text{-}D}$ = 20.0 Hz), 122.13 (t, $J_{C\text{-}D}$ = 22.5 Hz), 120.20, 112.83, 107.71; **HRMS**: calculated for $C_9H_6D_2NO_2^+$ [M+H$^+$]: 164.06751; found: 164.06719.

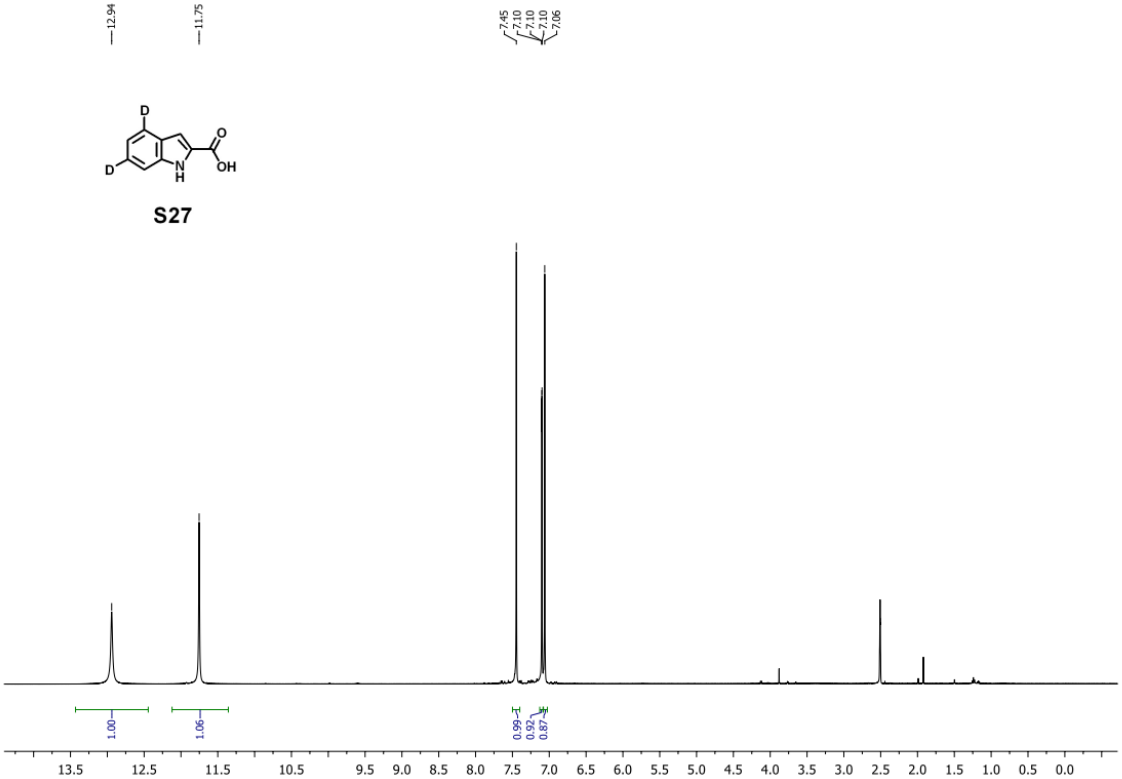
12.94
11.75
7.45
7.10
7.10
7.10
7.06
D
D
N
H
O
OH
S27
1.00
1.06
0.99
0.92
0.87
13.5
12.5
11.5
10.5
9.5
9.0
8.5
8.0
7.5
7.0
6.5
6.0
5.5
5.0
4.5
4.0
3.5
3.0
2.5
2.0
1.5
1.0
0.5
0.0

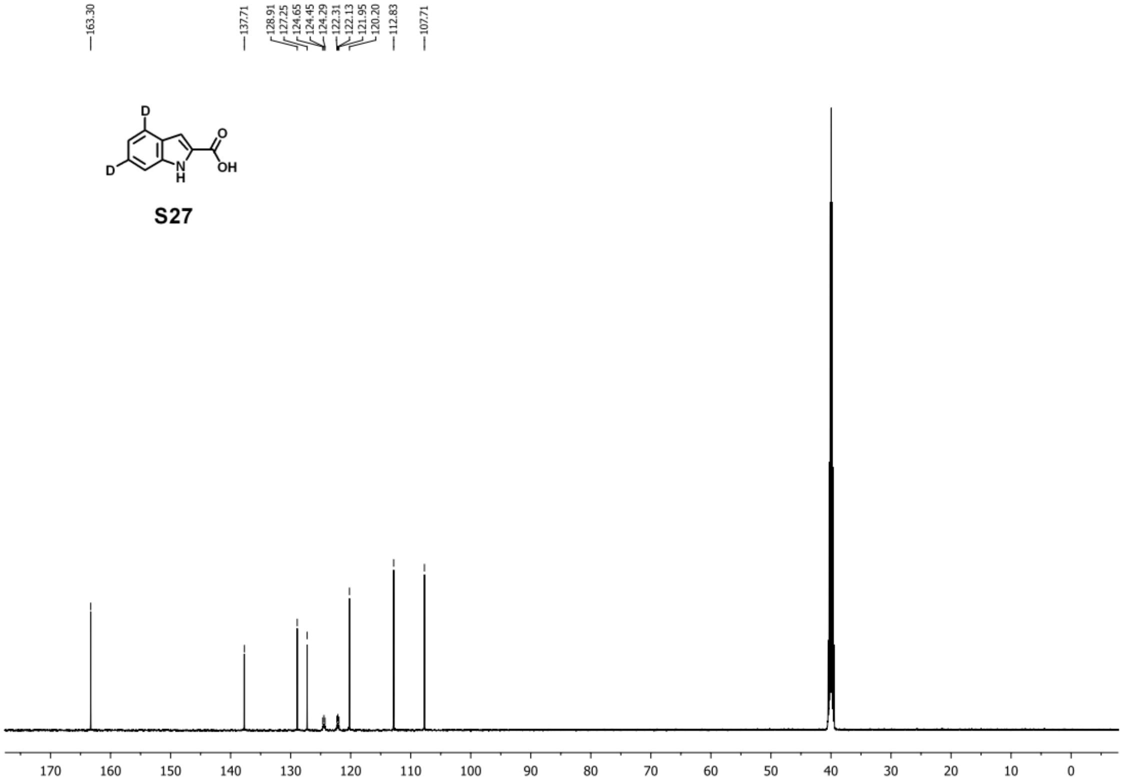
163.30
137.71
128.91
127.25
124.65
124.45
124.29
122.31
122.13
121.95
120.20
112.83
107.71
D
D
N
H
O
OH
S27
170
160
150
140
130
120
110
100
90
80
70
60
50
40
30
20
10
0

### 7.5. Synthesis of Fmoc-MIA-O-dipeptide-OH building blocks for SPPS

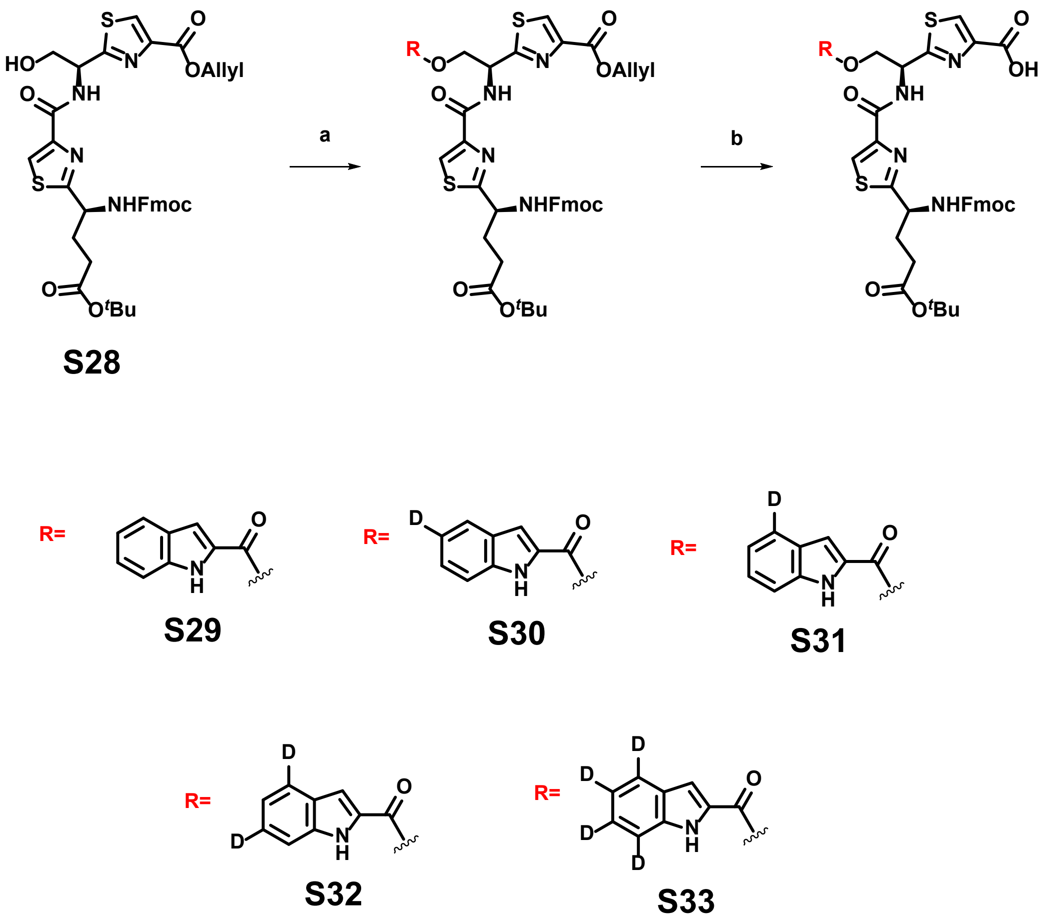


**Conditions and reagents:** a) 1*H*-indole-2-carboxylic acid or deuterated derivatives, DCC, DMAP, DCM, rt, 2h; b) $Pd(PPh_3)_4$, DMBA, THF, rt, 1h.

**Step a:**

Compound **S28** was synthesized according to the reported procedure.[24] To a stirred solution of **S28** (359 mg, 0.5 mmol, 1.0 equiv) and 1*H*-indole-2-carboxylic acid (161 mg, 1.0 mmol, 2.0 equiv) or deuterated derivatives (**S19**, **S20**, **S22**, or **S27**, 1.0 mmol, 2.0 equiv) in DCM (10 mL) was added DCC (124 mg, 0.6 mmol, 1.2 equiv) and DMAP (6.1 mg, 0.05 mmol, 0.1 equiv). The resulting reaction mixture was stirred at room temperature for 1h. After completion, the reaction mixture was diluted with dichloromethane (50 mL) and washed thoroughly with 0.1 M HCl, saturated aqueous sodium bicarbonate, and brine. The organic layer was dried over anhydrous sodium sulfate and concentrated *in vacuo*, and the resulting residue was subjected to next step without further purification.

**step b:**

To a stirred solution of the residue from previous step in THF (20 mL) was added $Pd(PPh_3)_4$ (29 mg, 0.025 mmol, 0.05 equiv) and 1,3-dimethylbarbituric acid (78 mg, 0.5 mmol, 1.0 equiv), and the resulting reaction mixture was stirred at room temperature for 1 h. Then, the reaction mixture was diluted with ethyl acetate (100 mL) and washed thoroughly with 0.1 M HCl, water, and brine. The organic layer was dried over anhydrous sodium sulfate and concentrated *in vacuo*, and the resulting residue was used for SPPS directly without further purification.

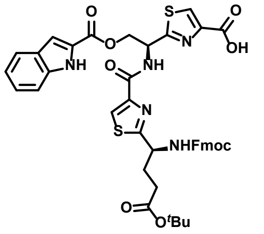


**S29** $^{1}$H NMR (400 MHz, DMSO) δ 11.99 (s, 1H), 9.31 – 9.04 (m, 1H), 8.32 (s, 1H), 8.29 (d, *J* = 8.0 Hz, 1H), 7.89 (d, *J* = 7.5 Hz, 2H), 7.75 (s, 1H), 7.70 (d, *J* = 7.3 Hz, 2H), 7.65 – 7.54 (m, 4H), 7.48 – 7.38 (m, 3H), 7.35 – 7.29 (m, 2H), 7.27 – 7.22 (m, 1H), 5.92 – 5.78 (m, 1H), 4.98 – 4.76 (m, 3H), 4.48 – 4.30 (m, 2H), 4.24 (t, *J* = 6.7 Hz, 1H), 2.33 – 2.25 (m, 2H), 1.77 – 1.56 (m, 2H), 1.39 – 1.28 (m, 9H); $^{13}$C NMR (101 MHz, DMSO) δ 175.22, 167.62, 164.68, 161.64, 161.25, 157.17, 156.41, 151.02, 149.53, 144.17, 141.24, 132.01, 131.92, 129.28, 129.17, 128.12, 127.54, 127.23, 127.11, 125.65, 122.59, 120.61, 120.41, 113.17, 108.52, 80.28, 66.04, 65.64, 53.02, 50.59, 47.98, 31.66, 29.93, 28.60; **HRMS**: calculated for $C_{42}H_{40}N_5O_9S_2^+$ [M+H$^+$]: 822.22620; found: 822.22477.

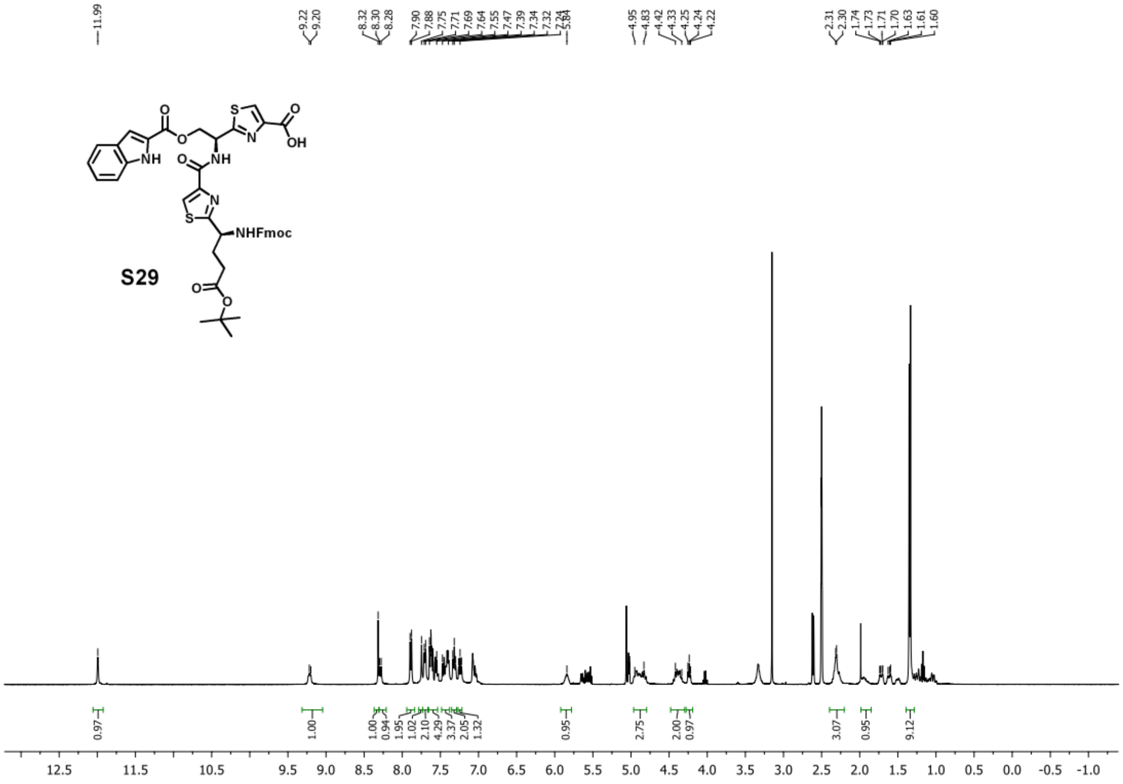
11.99
9.22
9.20
8.32
8.30
8.28
7.90
7.88
7.75
7.71
7.69
7.64
7.55
7.47
7.39
7.34
7.32
4.95
4.83
4.42
4.33
4.25
4.24
4.22
2.31
2.30
1.74
1.73
1.71
1.70
1.63
1.61
1.60
O
S
OH
N
NH
NH
O
N
S
NHFmoc
S29
O
O
0.97
1.00
1.00
0.94
1.95
1.02
2.10
4.29
3.37
2.05
1.32
0.95
2.75
2.00
0.97
3.07
0.95
9.12
12.5
11.5
10.5
9.5
9.0
8.5
8.0
7.5
7.0
6.5
6.0
5.5
5.0
4.5
4.0
3.5
3.0
2.5
2.0
1.5
1.0
0.5
0.0
-0.5
-1.0

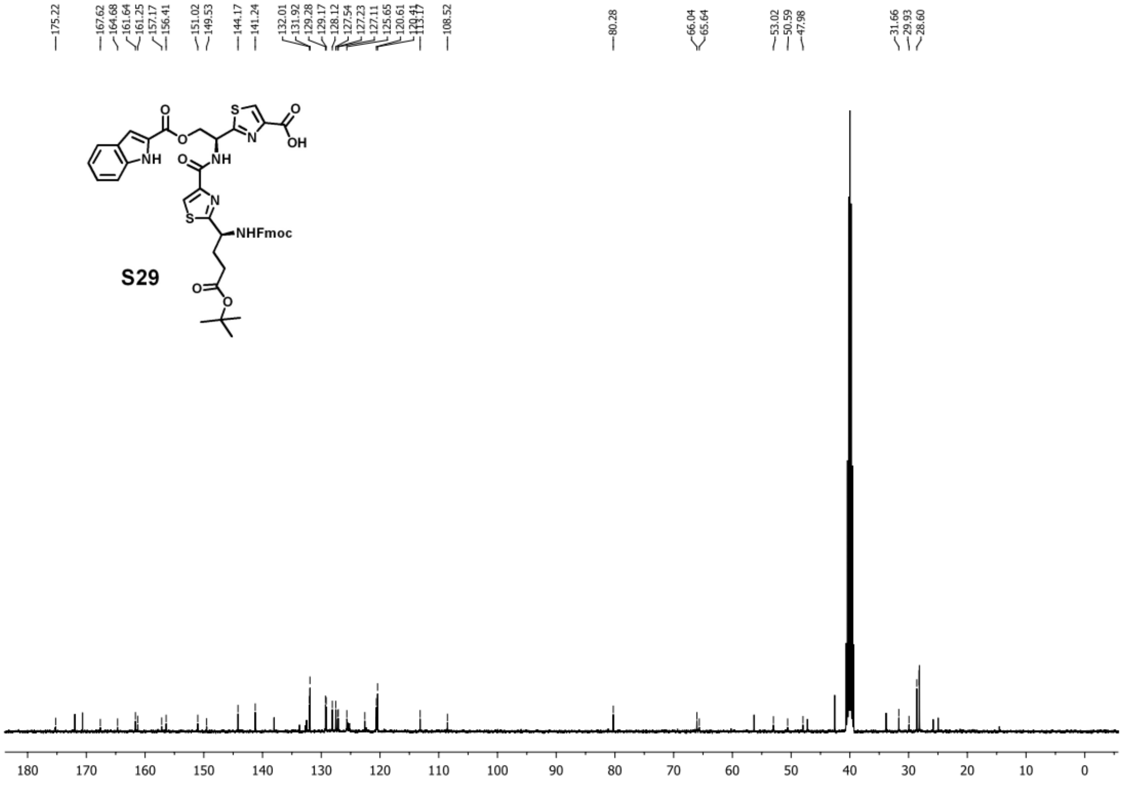
175.22
167.62
164.68
161.64
161.25
157.17
156.41
151.02
149.53
144.17
141.24
132.01
131.92
129.28
129.17
128.12
127.54
127.23
127.11
125.65
120.61
108.52
80.28
66.04
65.64
53.02
50.59
47.98
31.66
29.93
28.60
O
S
OH
N
NH
NH
O
N
S
NHFmoc
S29
O
O
180
170
160
150
140
130
120
110
100
90
80
70
60
50
40
30
20
10
0

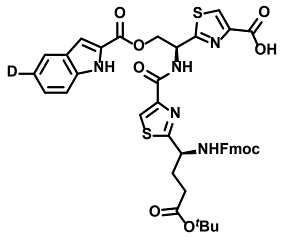

**S30** $^{1}H$ NMR (400 MHz, DMSO) δ 11.99 (s, 1H), 9.31 (s, 1H), 8.39 – 8.21 (m, 2H), 8.01 (s, 1H), 7.89 (d, *J* = 7.1 Hz, 2H), 7.70 (d, *J* = 5.8 Hz, 2H), 7.62 (s, 1H), 7.49 – 7.36 (m, 3H), 7.36 – 7.28 (m, 2H), 7.25 (d, *J* = 8.2 Hz, 1H), 7.09 (s, 1H), 5.90 (s, 1H), 4.88 (s, 2H), 4.50 – 4.28 (m, 2H), 4.28 – 4.18 (m, 1H), 2.39 – 2.17 (m, 3H), 1.94 (s, 1H), 1.41 – 1.26 (m, 9H); $^{13}C$ NMR (101 MHz, DMSO) δ 175.37, 171.95, 171.93, 161.55, 161.37, 156.41, 149.37, 144.15, 141.25, 138.01, 128.13, 127.53, 127.10, 125.59, 125.17, 122.50, 120.62, 113.08, 108.59, 80.27, 66.03, 65.63, 53.04, 50.92, 47.23, 31.64, 29.92, 28.13; **HRMS**: calculated for $C_{42}H_{39}DN_5O_9S_2^+$ [M+H$^+$]: 823.23247; found: 823.23045.

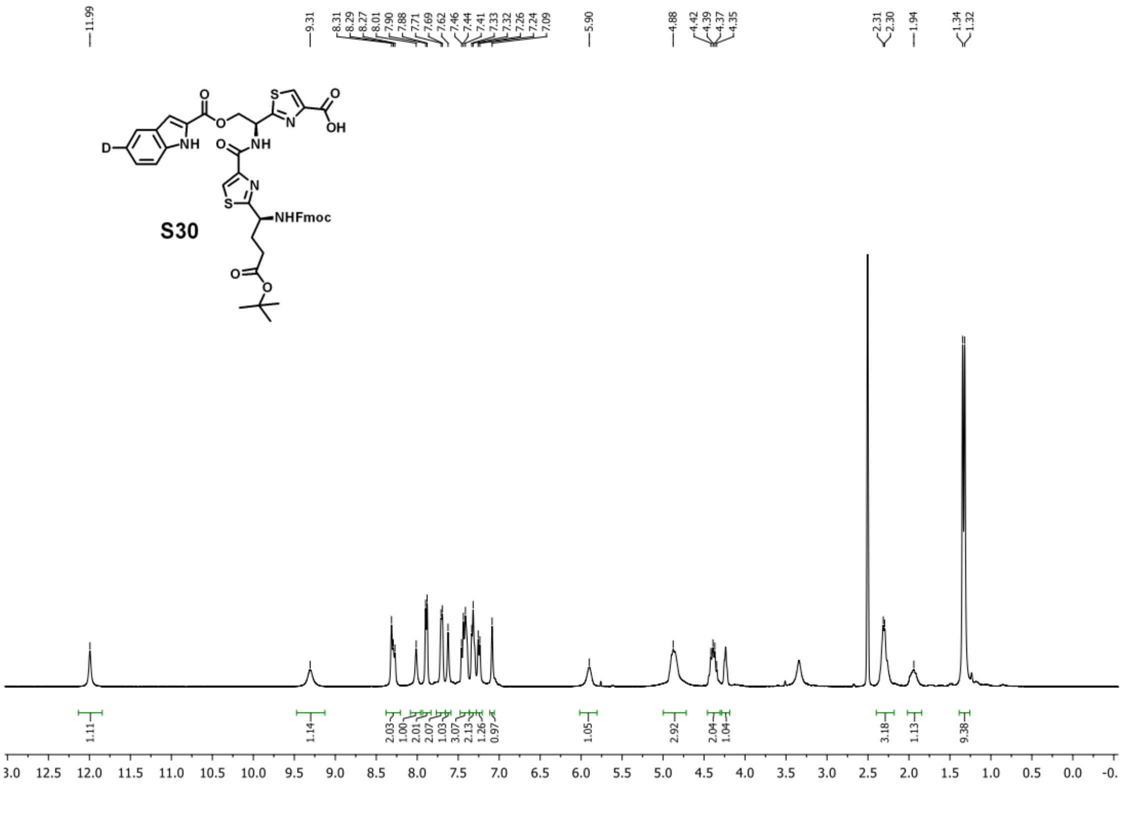

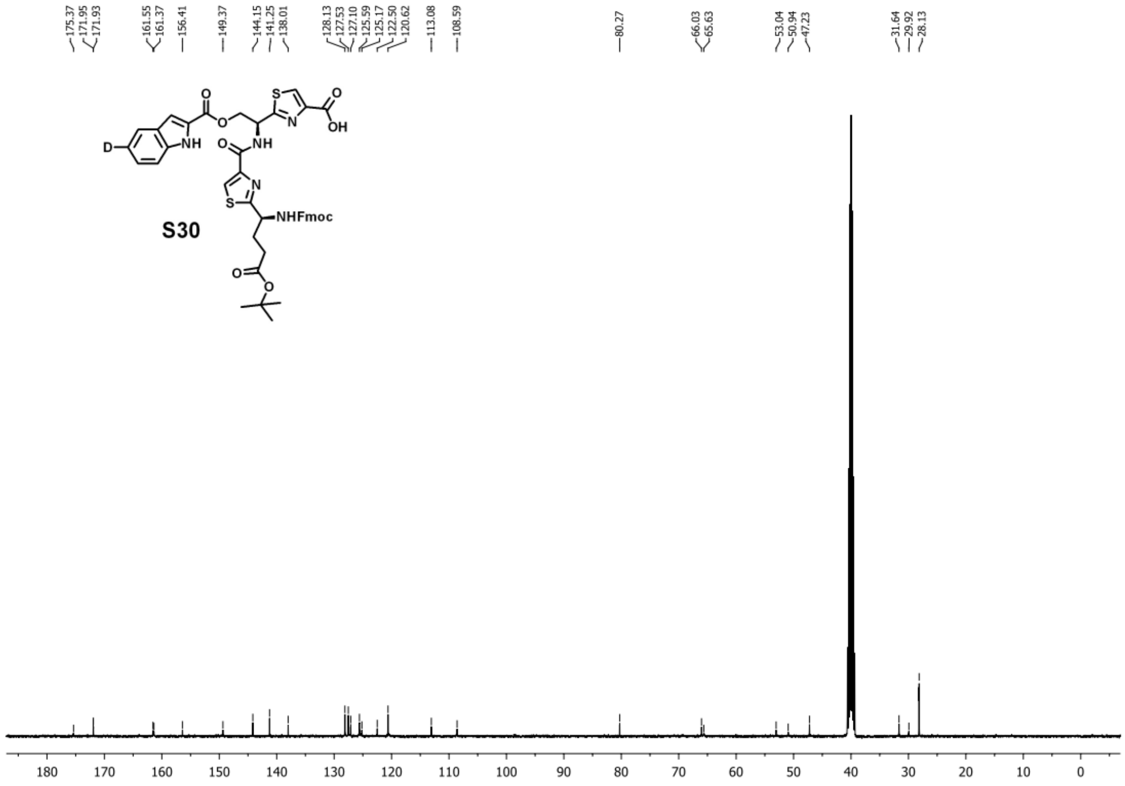


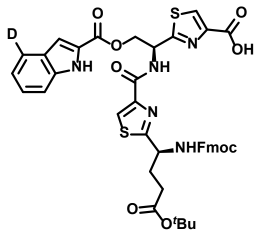

**S31** $^1$H NMR (400 MHz, DMSO) δ 12.00 (s, 1H), 9.23 (s, 1H), 8.37 – 8.22 (m, 2H), 7.89 (d, *J* = 7.5 Hz, 2H), 7.77 (s, 1H), 7.71 (d, *J* = 7.3 Hz, 2H), 7.66 – 7.59 (m, 3H), 7.58 – 7.52 (m, 2H), 7.48 – 7.39 (m, 3H), 7.36 – 7.29 (m, 2H), 7.27 – 7.21 (m, 1H), 5.85 (s, 1H), 4.99 – 4.77 (m, 3H), 4.46 – 4.31 (m, 2H), 4.24 (t, *J* = 7.2 Hz, 1H), 2.38 – 2.24 (m, 3H), 1.98 – 1.88 (m, 1H), 1.39 – 1.32 (m, 9H); $^{13}$C NMR (126 MHz, DMSO) δ 174.92, 167.37, 164.60, 161.58, 161.21, 157.29, 156.32, 151.06, 149.54, 144.24, 141.28, 131.99, 131.96, 129.18, 129.08, 128.05, 127.46, 127.33, 127.15, 125.53, 122.49, 120.48, 120.17, 113.18, 108.60, 80.29, 66.24, 65.64,

53.22, 50.85, 47.39, 31.84, 30.08, 28.52; **HRMS**: calculated for $C_{42}H_{39}DN_5O_9S_2^+$ [M+H$^+$]: 823.23247; found: 823.23083.

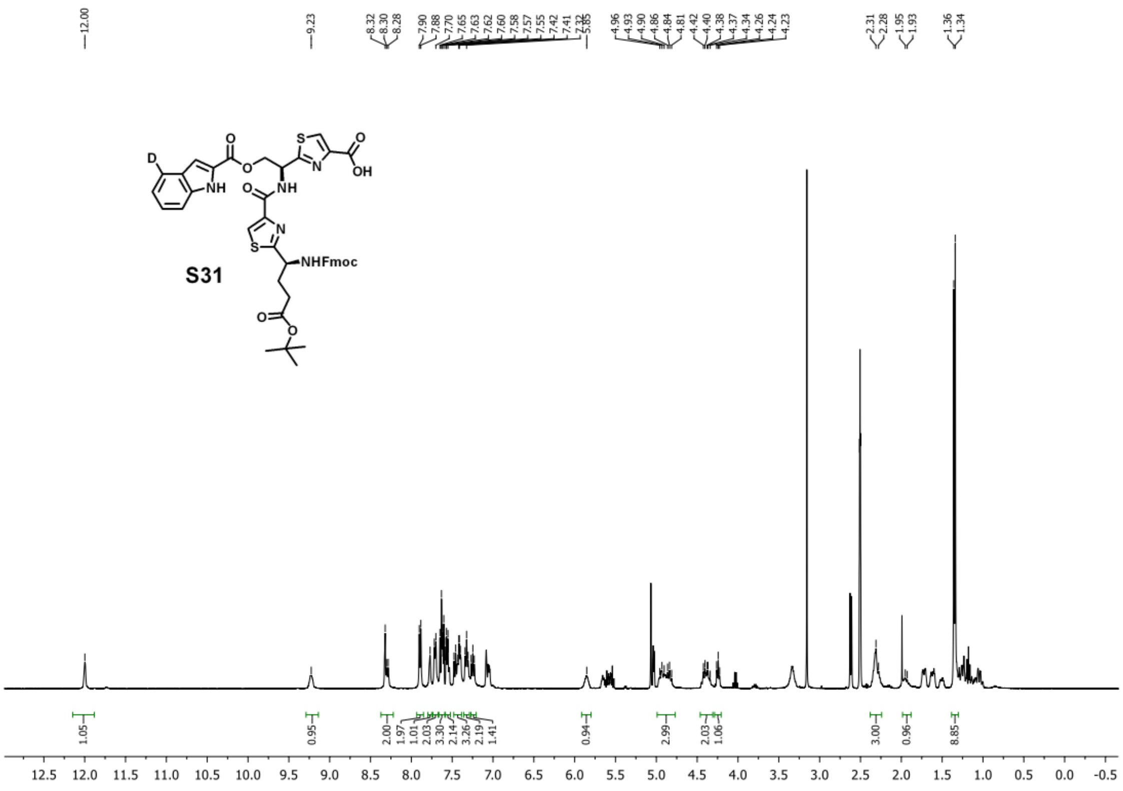

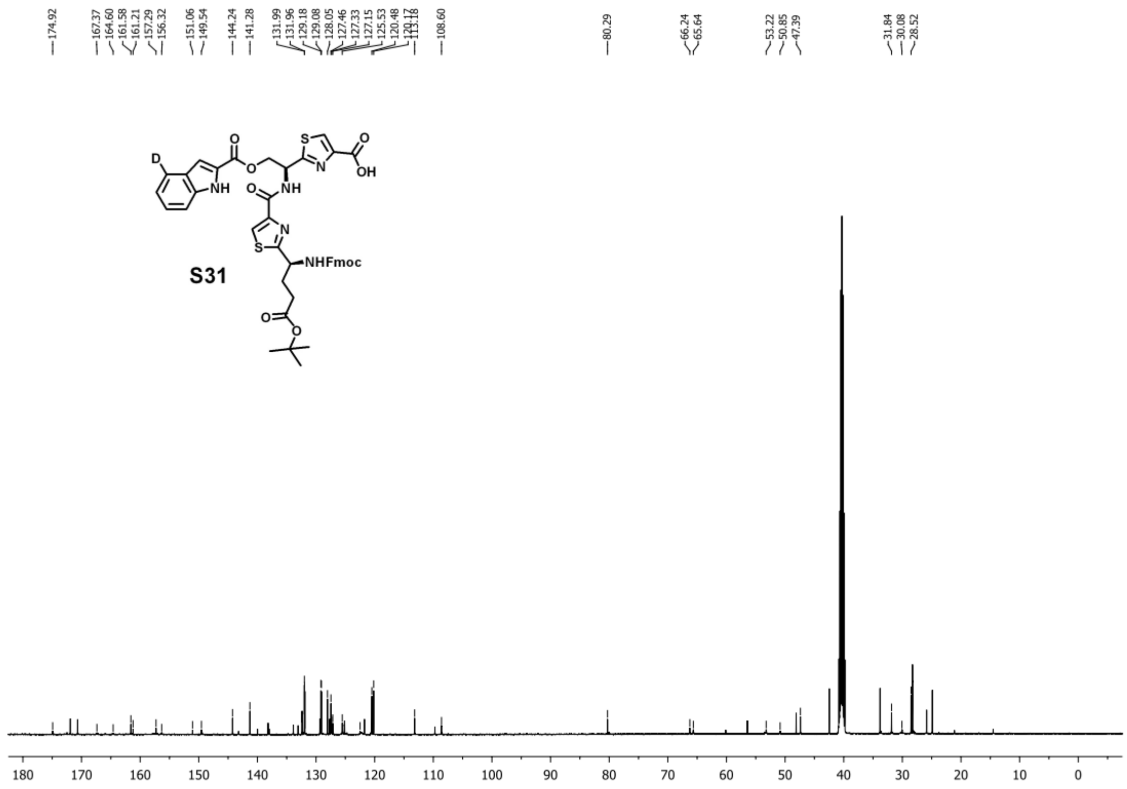


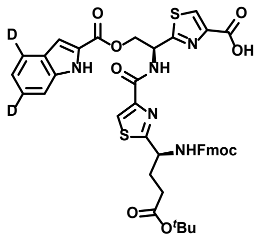

**S32** $^{1}$H NMR (400 MHz, DMSO) δ 11.97 (s, 1H), 9.19 (d, $J$ = 7.6 Hz, 1H), 8.37 – 8.19 (m, 2H), 7.89 (d, $J$ = 7.1 Hz, 2H), 7.77 (s, 1H), 7.75 – 7.67 (m, 2H), 7.66 – 7.59 (m, 2H), 7.59 – 7.53 (m, 1H), 7.48 – 7.36 (m, 5H), 7.36 – 7.28 (m, 3H), 5.83 (s, 1H), 4.96 – 4.79 (m, 2H), 4.47 – 4.29 (m, 2H), 4.28 – 4.19 (m, 1H), 2.39 – 2.22 (m, 3H), 1.98 – 1.90 (m, 1H), 1.38 – 1.32 (m, 9H); $^{13}$C NMR (101 MHz, DMSO) δ 175.30, 171.96, 161.24, 161.21, 156.41, 149.53, 144.17, 141.24, 138.04, 128.13, 127.54, 125.65, 125.60, 125.44, 122.03, 120.62, 113.04, 108.47, 80.28, 66.04, 65.67, 53.03, 50.52, 47.23, 31.66, 29.94, 28.60; **HRMS**: calculated for $C_{42}H_{38}D_2N_5O_9S_2^+$ [M+H$^+$]: 824.23875; found: 824.23788.

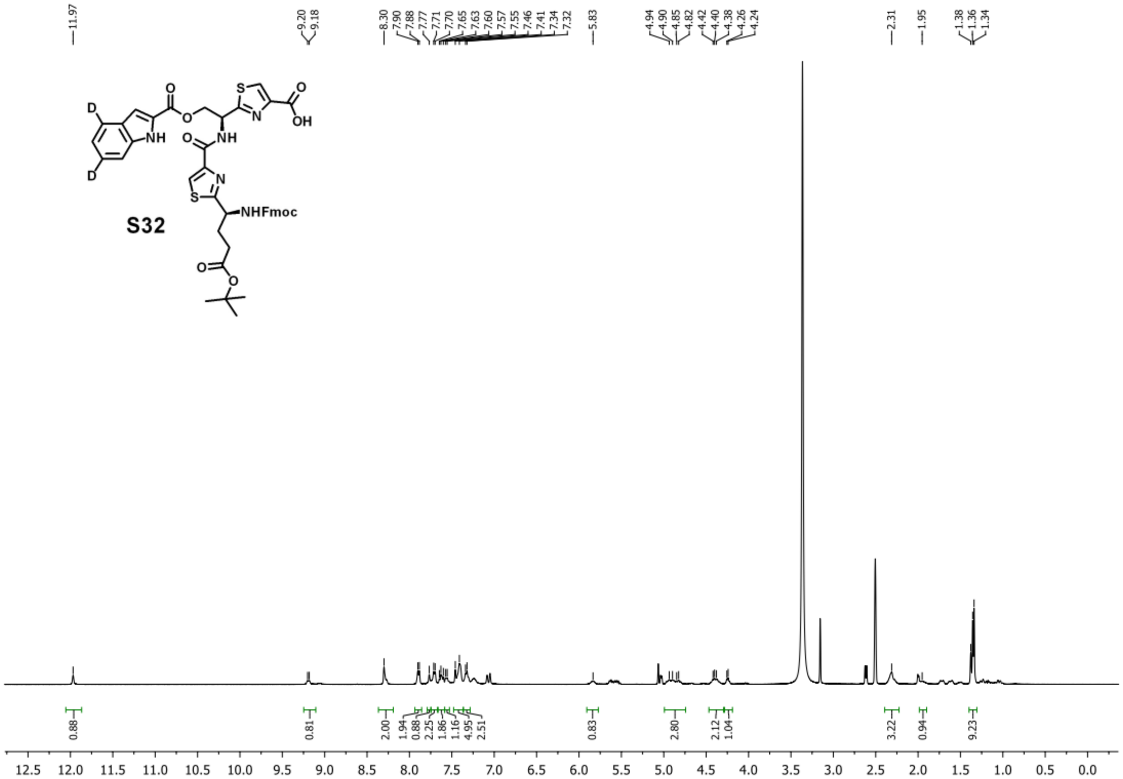
11.97
9.20 9.18
8.30 7.90 7.88 7.77 7.71 7.70 7.65 7.63 7.60 7.57 7.55 7.46 7.41 7.34 7.32
5.83
4.94 4.90 4.85 4.82 4.42 4.40 4.38 4.26 4.24
2.31
1.95
1.38 1.36 1.34
D
D
NH
O
NH
OH
S
N
NHFmoc
S32
0.88 0.81 2.00 1.94 0.88 2.25 1.86 1.16 4.95 2.51 0.83 2.80 2.12 1.04 3.22 0.94 9.23
12.5 12.0 11.5 11.0 10.5 10.0 9.5 9.0 8.5 8.0 7.5 7.0 6.5 6.0 5.5 5.0 4.5 4.0 3.5 3.0 2.5 2.0 1.5 1.0 0.5 0.0

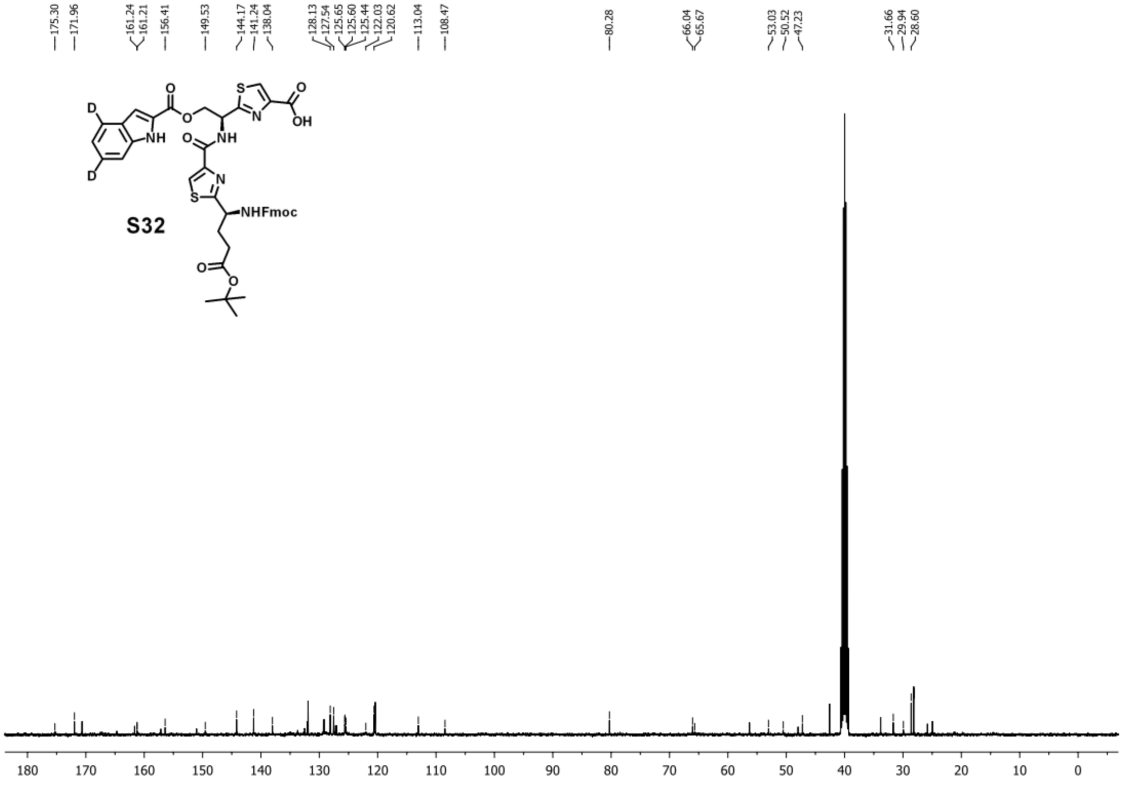
175.30 171.96
161.24 161.21 156.41
149.53
144.17 141.24 138.04
128.13 127.54 125.65 125.60 125.44 122.03 120.62
113.04
108.47
80.28
66.04 65.67
53.03 50.52 47.23
31.66 29.94 28.60
D
D
NH
O
NH
OH
S
N
NHFmoc
S32
180 170 160 150 140 130 120 110 100 90 80 70 60 50 40 30 20 10 0

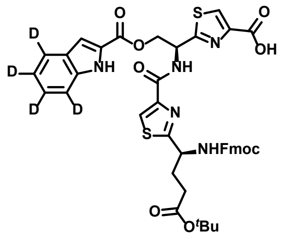

**S33** $^{1}$H NMR (400 MHz, DMSO) δ 11.98 (s, 1H), 9.21 (s, 1H), 8.37 – 8.21 (m, 2H), 7.89 (d, *J* = 7.3 Hz, 2H), 7.76 – 7.67 (m, 3H), 7.66 – 7.55 (m, 2H), 7.44 – 7.38 (m, 3H), 7.35 – 7.29 (m, 2H), 5.85 (s, 1H), 4.97 – 4.79 (m, 2H), 4.47 – 4.30 (m, 2H), 4.28 – 4.20 (m, 1H), 2.38 – 2.22 (m, 3H), 1.98 – 1.85 (m, 1H), 1.38 – 1.31 (m, 9H); $^{13}$C NMR (101 MHz, DMSO) δ 175.29, 171.96, 170.65, 161.66, 161.30, 156.42, 149.54, 144.17, 141.25, 137.99, 128.12, 127.54, 127.22, 125.65, 125.44, 122.08, 120.61, 108.48, 80.28, 66.04, 65.66, 53.02, 50.54, 47.23, 31.67, 29.94, 28.60; **HRMS**: calculated for $C_{42}H_{36}D_4N_5O_9S_2^+$ [M+H$^+$]: 826.25130; found: 826.25030.

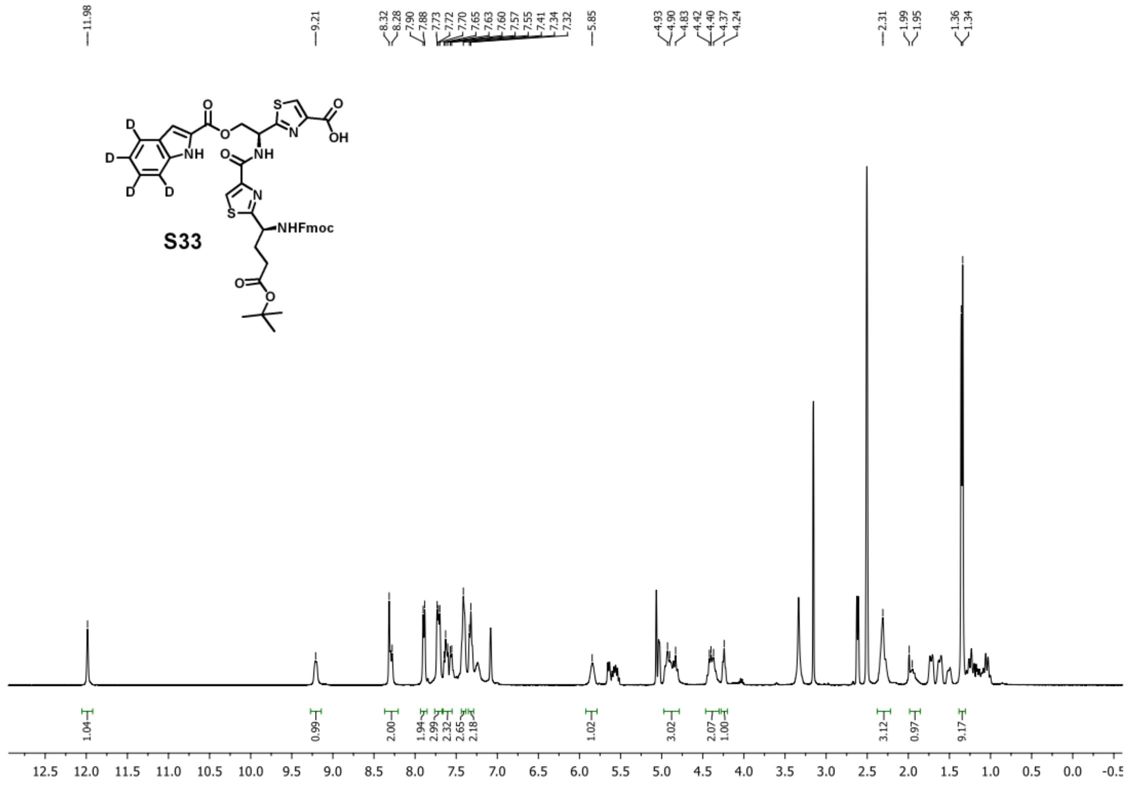

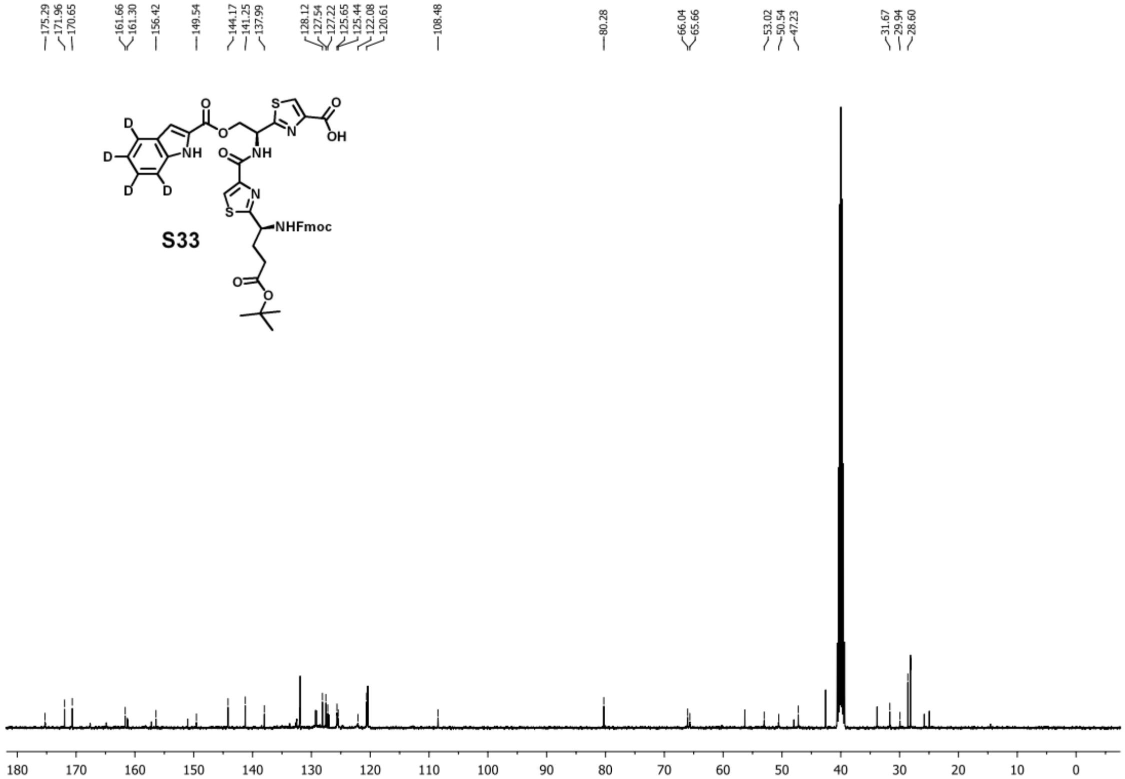


### 8.6. General protocol for solid phase peptide synthesis

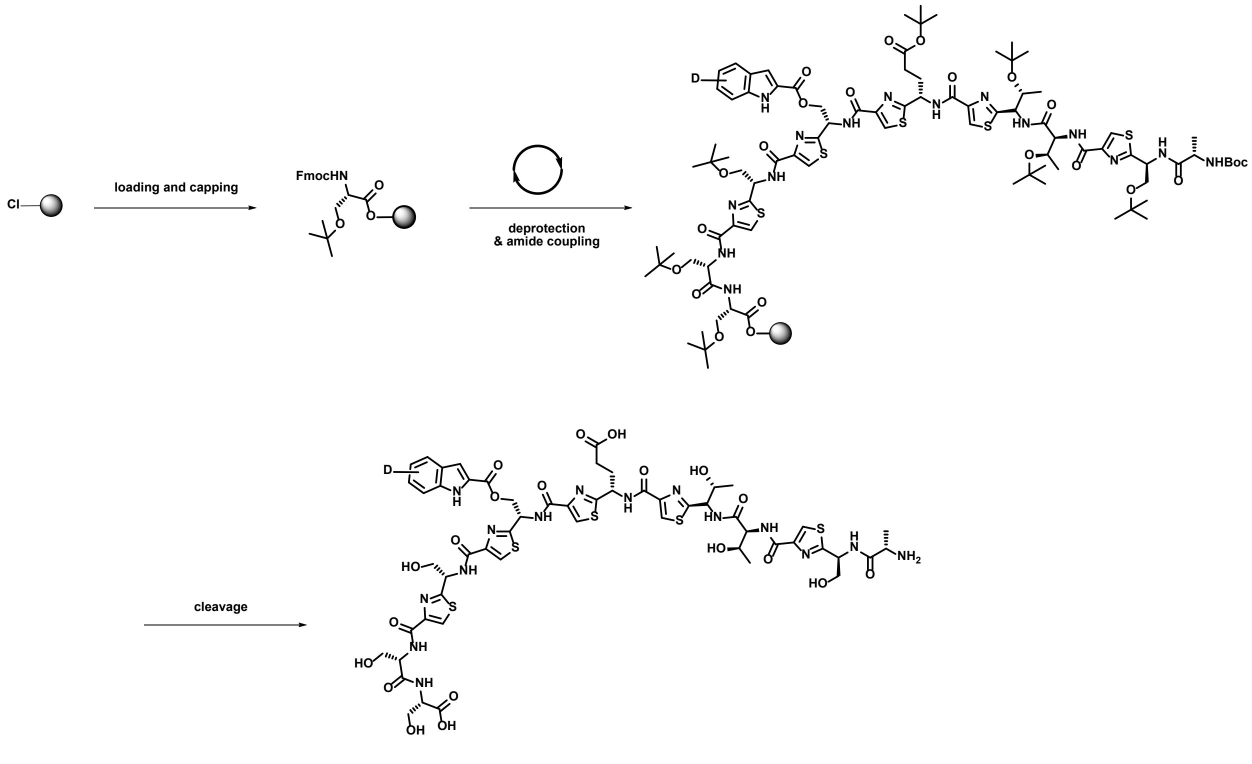

#### 7.6.1 Loading 2-chlorotrityl chloride resin

Weigh 2-chlorotrityl chloride resin (0.75 meq/g, 2 g, 1.5 mmol) in a 50 mL solid phase peptide synthesis vessel, then, swell the resin in DCM (20 mL) for 1 h. Drain the DCM, and dump a solution of Fmoc-L-Ser(*t*Bu)-OH (2.9 g, 7.5 mmol, 5 equiv) and 2,4,6-collidine (2 mL) in DCM (20 mL) into the vessel. Shake the vessel overnight, drain the reaction mixture and wash with DCM.

#### 7.6.2 Capping 2-chlorotrityl chloride resin

Dump the capping solution (DCM:MeOH:DIPEA=17mL:2mL:1mL) into the vessel and shake for 1h; drain the capping solution; wash the resin with DCM; dry the resin under nitrogen flow. The obtained resin is 0.7 mmol/g.

#### 7.6.3 SPPS

For the SPPS, we started from 0.1 mmol scale (equals to 145 mg resin from previous step).

Removal of Fmoc was achieved by treating the resin with 20% piperidine in DMF (4 mL) for 10 min. Then, the resin was washed with DMF and DCM.

For amide coupling, a solution of the corresponding amino acid building block (0.4 mmol, 4.0 equiv) and HATU (152 mg, 0.4 mmol, 4.0 equiv) in DMF containing 20% NMM (5 mL) was added to the vessel and shaken for 20 min. After completion, the reaction solution was dumped and washed with DMF followed by DCM.

Repeat the deprotecting and amide coupling step until all the amino acids are incorporated.

The last amino acid to be incorporated usually is Boc-L-Ala, instead of Fmoc-L-Ala, for the one-pot deprotection and cleavage from the resin.

#### 7.6.4 Cleavage and Deprotection

After all the amino acid building blocks were incorporated, a cleavage cocktail (TFA:$H_2O$:triisopropylsilane=9.5mL:0.25mL:0.25mL) was added to the resin, and the resulting reaction mixture was shaken for 3 hours. Then, the solution was collected in a 50 mL conical tube. The volume was reduced to ~4 mL under nitrogen flow, and cold *t*-butyl methyl ether (40 mL) was added to precipitate the peptide. The precipitation was collected by centrifuge and purified by HPLC.

#### 7.6.5 HPLC Purification

The purification of crude peptides is performed on an Agilent 1260 Infinity II Preparative HPLC system using an Agilent 5 Prep-C18 (50 × 21.2 mm) column. The solvents for purification include 0.1% TFA in water (solvent A) and 0.1% TFA in acetonitrile (solvent B), and detection is performed by UV monitoring at 254 nm. The detailed methods are listed below:

| Time (min) | A% | B% | Flow rate (ml/min) |
|---|---|---|---|
| 0.00 | 75 | 23 | 30.00 |
| 2.00 | 75 | 23 | 30.00 |
| 5.00 | 60 | 34.5 | 30.00 |
| 5.20 | 20 | 80 | 30.00 |
| 6.00 | 20 | 80 | 30.00 |
| 6.20 | 75 | 23 | 30.00 |
| 7.00 | 75 | 23 | 30.00 |

Each peptide elutes at around 3.5 min. Iterative injections were performed to purify all the crude peptides. Fractions are pooled and lyophilized to give a white powder as the final product. The powder is re-dissolved in water to make a stock solution with a concentration of 5 mM. The HPLC traces, chemical structures, and HRMS are shown below:

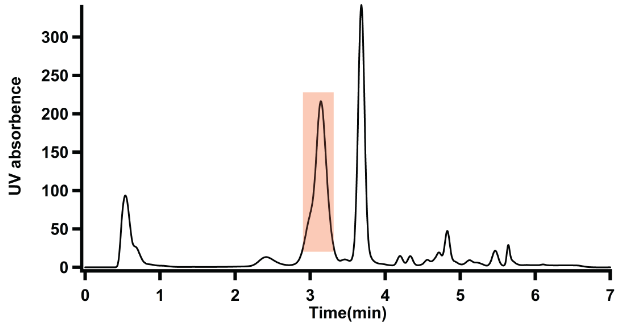


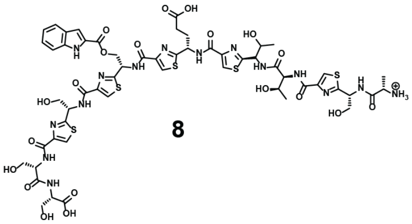

**8**

Calculated for $C_{55}H_{64}N_{15}O_{20}S_5^+$ [M+H$^+$]: 1414.30501; found: 1414.30387.

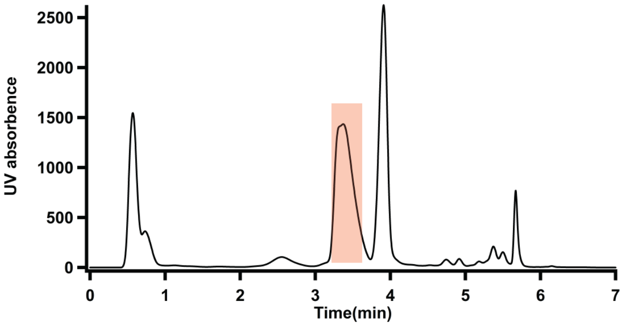


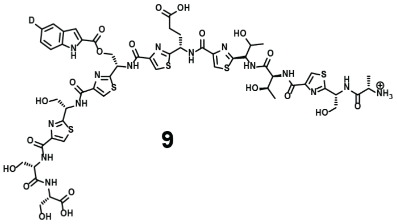

**9**

Calculated for $C_{55}H_{63}DN_{15}O_{20}S_5^+$ [M+H$^+$]: 1415.31129; found: 1415.31036.

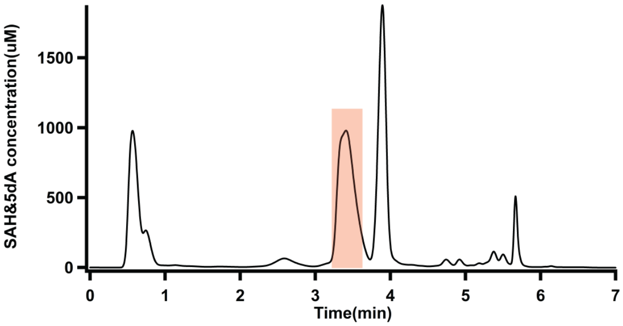


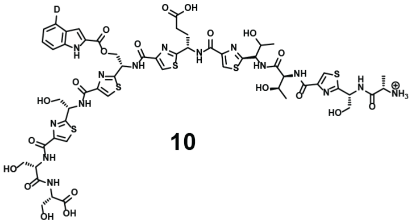

**10**

Calculated for $C_{55}H_{63}DN_{15}O_{20}S_5^+$ [M+H$^+$]: 1415.31129; found: 1415.31022.

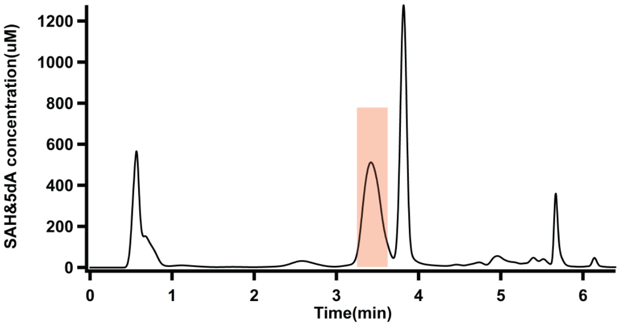


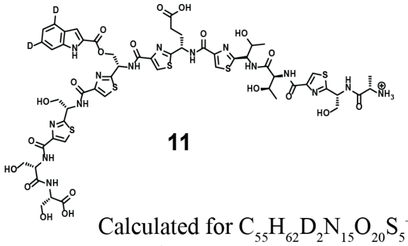

**11**

Calculated for $C_{55}H_{62}D_2N_{15}O_{20}S_5^+$ [M+H$^+$]: 1416.31756; found: 1416.31631.

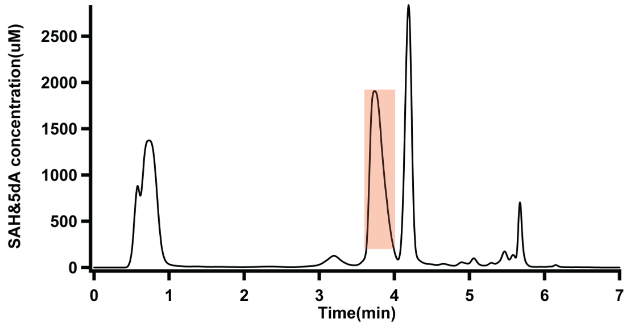


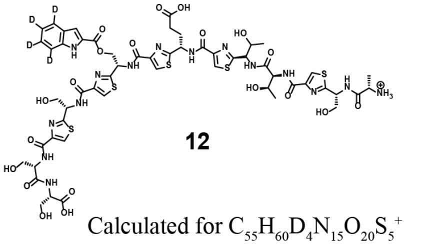

**12**

Calculated for $C_{55}H_{60}D_4N_{15}O_{20}S_5^+$ [M+H$^+$]: 1418.33012; found: 1418.32919.

1 Iwig, D. F. & Booker, S. J. Insight into the polar reactivity of the onium chalcogen analogues of S-adenosyl-L-methionine. *Biochemistry-Us* **43**, 13496-13509 (2004).

2 LaMattina, J. W. *et al.* NosN, a Radical S-Adenosylmethionine Methylase, Catalyzes Both C1 Transfer and Formation of the Ester Linkage of the Side-Ring System during the Biosynthesis of Nosiheptide. *J Am Chem Soc* **139**, 17438-17445 (2017).

3 Adams, P. D. *et al.* PHENIX: a comprehensive Python-based system for macromolecular structure solution. *Acta Crystallogr D* **66**, 213-221, doi:10.1107/S0907444909052925 (2010).

4 Bunkoczi, G. *et al.* Phaser.MRage: automated molecular replacement. *Acta Crystallogr D* **69**, 2276-2286, doi:10.1107/S0907444913022750 (2013).

5 Minor, W., Cymborowski, M., Otwinowski, Z. & Chruszcz, M. HKL-3000: the integration of data reduction and structure solution - from diffraction images to an initial model in minutes. *Acta Crystallogr D* **62**, 859-866, doi:10.1107/S0907444906019949 (2006).

6 Otwinowski, Z. & Minor, W. Processing of X-ray diffraction data collected in oscillation mode. *Macromolecular Crystallography, Pt A* **276**, 307-326, doi:Doi 10.1016/S0076-6879(97)76066-X (1997).

7 Emsley, P., Lohkamp, B., Scott, W. G. & Cowtan, K. Features and development of Coot. *Acta Crystallogr D* **66**, 486-501, doi:10.1107/S0907444910007493 (2010).

8 Chen, V. B. *et al.* MolProbity: all-atom structure validation for macromolecular crystallography. *Acta Crystallographica Section D-Structural Biology* **66**, 12-21, doi:10.1107/S0907444909042073 (2010).

9 Dunbar, K. L., Tietz, J. I., Cox, C. L., Burichart, B. J. & Mitchell, D. A. Identification of an Auxiliary Leader Peptide-Binding Protein Required for Azoline Formation in Ribosomal Natural Products. *J Am Chem Soc* **137**, 7672-7677 (2015).

10 Kung, Y., Doukov, T. I., Seravalli, J., Ragsdale, S. W. & Drennan, C. L. Crystallographic Snapshots of Cyanide- and Water-Bound C-Clusters from Bifunctional Carbon Monoxide Dehydrogenase/Acetyl-CoA Synthase. *Biochemistry-Us* **48**, 7432-7440, doi:10.1021/bi900574h (2009).

11 Wang, B., Silakov, A. & Booker, S. J. in *Methods in Enzymology* Vol. 666 (ed R. David Britt) 469-487 (Academic Press, 2022).

12 Neese, F. The ORCA program system. *WIREs Computational Molecular Science* **2**, 73-78, doi:https://doi.org/10.1002/wcms.81 (2012).

13 Neese, F. Software update: The ORCA program system—Version 5.0. *WIREs Computational Molecular Science* **12**, e1606, doi:https://doi.org/10.1002/wcms.1606 (2022).

14 Staroverov, V. N., Scuseria, G. E., Tao, J. & Perdew, J. P. Comparative assessment of a new nonempirical density functional: Molecules and hydrogen-bonded complexes. *The Journal of Chemical Physics* **119**, 12129-12137, doi:10.1063/1.1626543 (2003).

15 Tao, J., Perdew, J. P., Staroverov, V. N. & Scuseria, G. E. Climbing the density functional ladder: nonempirical meta-generalized gradient approximation designed for molecules and solids. *Phys Rev Lett* **91**, 146401, doi:10.1103/PhysRevLett.91.146401 (2003).

16 Weigend, F. & Ahlrichs, R. Balanced basis sets of split valence, triple zeta valence and quadruple zeta valence quality for H to Rn: Design and assessment of accuracy. *Phys Chem Chem Phys* **7**, 3297-3305, doi:10.1039/b508541a (2005).

17 Weigend, F. Accurate Coulomb-fitting basis sets for H to Rn. *Phys Chem Chem Phys* **8**, 1057-1065, doi:10.1039/b515623h (2006).

18 Stephens, P. J., Devlin, F. J., Chabalowski, C. F. & Frisch, M. J. Ab Initio Calculation of Vibrational Absorption and Circular Dichroism Spectra Using Density Functional Force Fields. *The Journal of Physical Chemistry* **98**, 11623-11627, doi:10.1021/j100096a001 (1994).

19 Helmich-Paris, B., de Souza, B., Neese, F. & Izsák, R. An improved chain of spheres for exchange algorithm. *The Journal of Chemical Physics* **155**, doi:10.1063/5.0058766 (2021).
20 Grimme, S., Antony, J., Ehrlich, S. & Krieg, H. A consistent and accurate ab initio parametrization of density functional dispersion correction (DFT-D) for the 94 elements H-Pu. *Journal of Chemical Physics* **132**, 154104, doi:10.1063/1.3382344 (2010).
21 Cossi, M., Rega, N., Scalmani, G. & Barone, V. Energies, structures, and electronic properties of molecules in solution with the C-PCM solvation model. *Journal of Computational Chemistry* **24**, 669-681, doi:https://doi.org/10.1002/jcc.10189 (2003).
22 Kendall, R. A. & Früchtl, H. A. The impact of the resolution of the identity approximate integral method on modern ab initio algorithm development. *Theoretical Chemistry Accounts* **97**, 158-163, doi:10.1007/s002140050249 (1997).
23 Liu, Y. *et al.* One-Pot Enantiomeric Synthesis of Thiazole-Containing Amino Acids: Total Synthesis of Venturamides A and B. *J Org Chem* **83**, 3897-3905, doi:10.1021/acs.joc.8b00244 (2018).
24 Wang, B., LaMattina, J. W., Marshall, S. L. & Booker, S. J. Capturing Intermediates in the Reaction Catalyzed by NosN, a Class C Radical S-Adenosylmethionine Methylase Involved in the Biosynthesis of the Nosiheptide Side-Ring System. *J Am Chem Soc* **141**, 5788-5797, doi:10.1021/jacs.8b13157 (2019).
